# SARDINE unlocks EWAS summary statistics for large-scale epigenetic biomarker analysis

**DOI:** 10.64898/2026.11.05.763758

**Authors:** Yuchang Wu, Yushan Wang, Ryan Moreno, Sarah M. Merrill, Varun B. Dwaraka, Kirsten B. Seale, Ryan Smith, Lauren L. Schmitz, Qiongshi Lu

## Abstract

Epigenetic clocks are increasingly used to study biological aging, yet their interpretation and utility remain uncertain. Meanwhile, the growing wealth of epigenome-wide association study (EWAS) summary statistics remains largely untapped for evaluating these biomarkers. Here, we introduce SARDINE, a statistical framework that estimates epigenetic clock-phenotype associations using EWAS summary statistics, clock weights, and reference methylation covariance, without individual-level data. SARDINE accurately reproduced individual-level association estimates across 38 epigenetic biomarkers and four phenotypes in 15,677 individuals. Applying SARDINE to diverse EWAS datasets revealed substantial heterogeneity in biomarker associations with aging, diseases, and lifestyle factors. By transforming CpG-level EWAS results into interpretable biomarker-level associations, SARDINE unlocks a largely underutilized data resource and establishes a scalable paradigm for systematic biomarker evaluation and discovery in aging research.

## Introduction

Epigenetic clocks have emerged as powerful molecular biomarkers of biological aging^1–6^. Constructed from DNA methylation (DNAm) measurements at tens to thousands of cytosine-phosphate-guanine (CpG) sites, these models have evolved from predictors of chronological age^7,8^ to biomarkers of healthspan^9,10^, mortality risk^11^, and the pace of aging^12^. More broadly, DNAm-based predictors have been developed for diverse health^13^, behavioral^14^, and socioeconomic^14,15^ phenotypes, expanding the potential applications of epigenetic biomarkers in human research. These advances have established DNAm-based biomarkers as valuable predictive tools for investigating the effects of socioeconomic conditions^16,17^, lifestyle factors^9,10,18^, environmental exposures^19^, disease processes^10,11,20^, and interventions targeting healthy aging^4,21,22^.

Despite their widespread use, important questions remain regarding the biological interpretation, specificity, and practical utility of epigenetic clocks^4,6,23^. Epigenetic clocks are most often assessed via epigenetic age acceleration (EAA), which reflects discrepancies in DNAm-predicted age relative to chronological age. However, the increasing diversity of DNAm-based biomarkers complicates this interpretation, as does the fact that different clocks often exhibit modest correlations with one another^24,25^ and can show substantially divergent associations of EAA with the same phenotype, raising questions about whether they capture shared dimensions of biological aging or distinct biological processes. Systematic benchmarking of epigenetic clocks in a large sample across diverse phenotypes is therefore essential for understanding what these models measure and what modifiable factors they are sensitive to, evaluating their potential utility, and guiding future biomarker development. Recent comparative studies have demonstrated the value of this approach^20^, but comprehensive benchmarking remains challenging. Conventional analyses require individual-level DNAm data, extensive phenotypic measurements, and substantial computational resources. Access to such datasets is often restricted. Some cohorts release precomputed clock values, but these are rarely updated to incorporate newly developed models. Furthermore, no single cohort captures the full spectrum of phenotypes relevant to biological aging. These barriers substantially limit the scale, scope, and reproducibility of DNAm-based biomarker research.

Meanwhile, the rapidly expanding collection of epigenome-wide association studies (EWAS) represents a largely untapped resource for addressing these challenges. Thousands of EWAS have investigated DNAm associations with diverse human phenotypes, including lifestyle factors, environmental exposures, socioeconomic conditions, and diseases^18,26–33^. Increasingly, EWAS summary statistics are made publicly available through publications^33^ and centralized repositories^34^, providing opportunities to investigate epigenetic biomarker associations across studies and populations without accessing individual-level data. However, a fundamental methodological gap prevents their widespread use for this purpose: EWAS quantify phenotype associations at individual CpG sites, whereas epigenetic clocks integrate information across many CpGs. To our knowledge, no existing statistical approach enables direct estimation of epigenetic clock-phenotype associations from EWAS summary statistics. Consequently, translating CpG-level associations into interpretable clock-level biological insights remains difficult. This disconnect leaves a rapidly expanding public data resource for addressing key questions in biological aging largely underutilized.

Here, we introduce SARDINE (Summary-statistics-based Association Regression for Deciphering INtegrated CpG Effects), a statistical framework that bridges this methodological gap by estimating epigenetic clock-phenotype associations from EWAS summary statistics. By integrating CpG-level association estimates, published biomarker weights, and reference DNAm covariance matrices for adult whole blood, SARDINE eliminates the need for individual-level DNAm data and enables scalable association analysis across many clock models, phenotypes, and cohorts. The framework accommodates diverse EWAS designs, supports covariate-adjusted inference, and applies broadly to DNAm-based predictors constructed as weighted combinations of CpG measurements. We validate SARDINE against conventional individual-level analyses and demonstrate its utility through systematic benchmarking of epigenetic clocks and related biomarkers across numerous phenotypes. By transforming publicly available EWAS resources into a powerful platform for aging research, SARDINE establishes a scalable framework for evaluating existing and future DNAm-based biomarkers, revealing shared and distinct phenotypic association patterns, and guiding the development of future epigenetic aging measures.

## Results

### Method overview

Epigenetic aging studies commonly assess the association between an epigenetic clock and a phenotype, a term we use broadly for any exposure or outcome of interest. The SARDINE framework can be applied to any linear DNAm-based predictor. Indeed, some models included in later analyses are DNAm surrogates for complex traits rather than conventional epigenetic clocks that track chronological or biological aging. Throughout this paper, we use the term “clock” broadly to encompass both epigenetic clocks and other DNAm-based predictors.

Existing studies focus on (1) whether an exposure (e.g., smoking, supplement intake) influences epigenetic aging, and/or (2) whether epigenetic aging influences health outcomes (e.g., disease diagnosis, mortality). These questions are investigated through the regressions

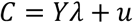

and

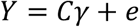

where *C* denotes an epigenetic aging clock, *Y* is the phenotype of interest, and *u* and *e* are noise terms in the regression. Both *C* and *Y* are centered at zero so intercepts are omitted. The main inference task is to estimate and perform hypothesis testing on effect parameters *λ* and *γ*, which quantify the association between the clock and the phenotype.

A key methodological contribution of this work is demonstrating that *λ* and *γ* can be robustly estimated from (i) EWAS summary statistics 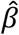 (i.e., DNAm-phenotype associations at the CpG level), (ii) CpG weight values *w* in an epigenetic clock model, and (iii) variance-covariance matrix *Σ* of DNAm across CpGs included in the clock model (**Figure 1** and **Supplementary Figure 1**). The point estimates of *λ* and *γ* are (**Methods** and **Supplementary Note**)

**Figure 1.**
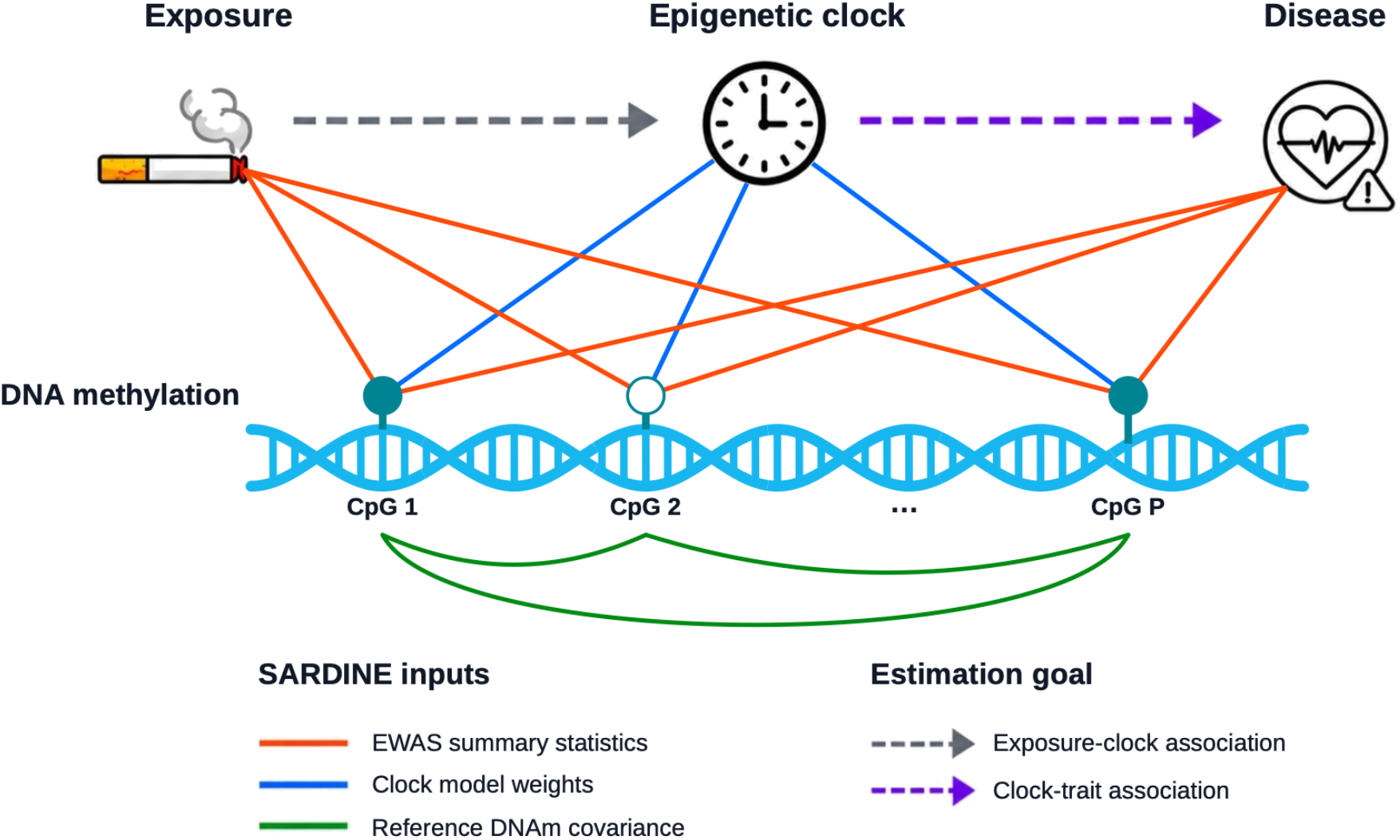
SARDINE overview. Our approach uses EWAS summary statistics, clock weights, and DNAm variance-covariance among CpG sites to estimate two types of clock-phenotype associations: (i) the effect of an exposure on epigenetic aging; and (ii) the effect of epigenetic aging on a phenotype.

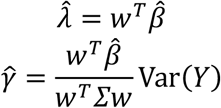

CpG-phenotype associations 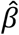 (and their standard errors) and clock weights *w* are often publicly released upon EWAS and clock method publication. We also show that matrix *Σ* can be reliably estimated from a reference DNAm dataset and phenotypic variance Var(*Y*) can be approximated using EWAS summary statistics and the estimated *Σ*. Therefore, clock-phenotype association analysis can be performed without using individual-level measures of *C* or *Y*. This enables systematic and scalable evaluation of DNAm-based biomarkers using increasingly available summary-level data resources.

Details of the SARDINE statistical framework are described in **Methods** and **Supplementary Note**. There, we also discuss and provide solutions to important technical issues, including approaches for estimating standard errors, adjusting for covariates, handling complex phenotypes (e.g., binary and survival outcomes), handling EWAS associations obtained from transformed DNAm measures, and reporting *γ* on interpretable scales (e.g., log odds ratio or log hazard ratio).

### Simulations validate summary-statistics-based inference

We conducted extensive simulations to evaluate SARDINE’s performance based on real DNAm data from a subset of 8,880 TruDiagnostic participants (**Methods**). We simulated continuous and binary phenotypes based on a variety of realistic model assumptions and parameter settings (**Methods**). We estimated *λ* and *γ* using SARDINE with EWAS summary statistics in which DNAm was either the outcome (M ~ Y) or the predictor (Y ~ M), and compared the results with conventional individual-level regression results. To assess the impact of using an external DNAm reference, we used DNAm covariance estimated either in-sample or from independent holdout datasets and compared the results. Each setting was repeated 2,000 times. Details on these simulations are described in **Methods** and **Supplementary Note**.

Results based on individual-level data and summary statistics were near-identical across all simulation settings. SARDINE maintained well-controlled type-I error rates under the null and achieved high statistical power under the presence of real clock-phenotype associations (**Figure 2A** and **Supplementary Figure 2**). SARDINE also provided unbiased effect size estimates for both continuous and binary phenotypes within a reasonable and expected effect size range (**Figure 2B**), given two types of input EWAS treating DNAm as either the outcome or the predictor in regression (**Methods**). In addition, both in-sample and external DNAm reference yielded similar estimation results, and the results were also robust to varying sample sizes in the DNAm reference dataset (**Supplementary Figures 3-4**).

**Figure 2.**
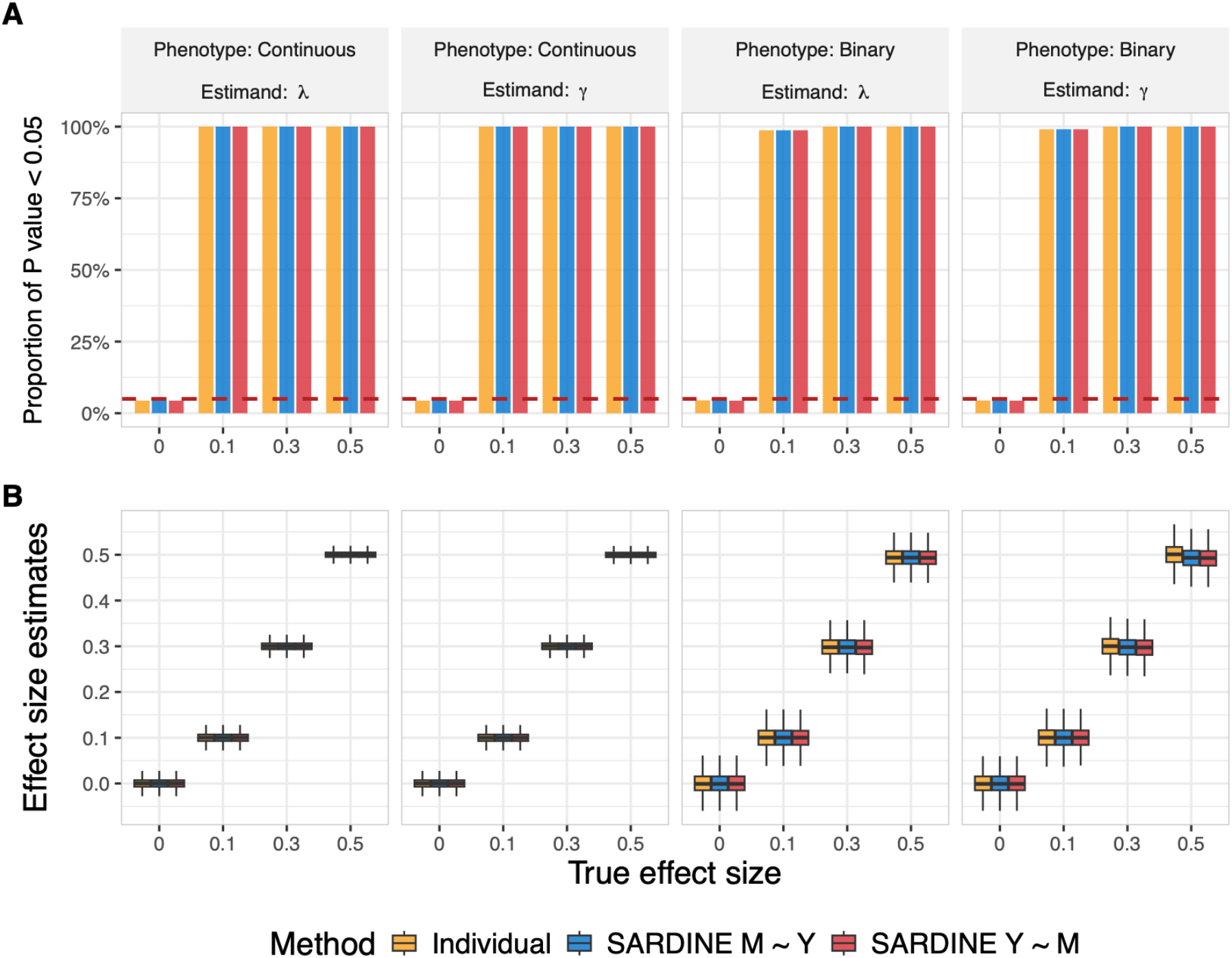
Simulation results. SARDINE and a conventional individual-level regression approach were used to estimate clock-phenotype associations under various simulation settings. SARDINE used an in-sample DNAm covariance matrix and EWAS summary statistics from one of the two specifications: DNAm regressed on the phenotype (M ~ Y) or the phenotype regressed on DNAm (Y ~ M). Both *λ* (i.e., the effect of phenotype on the clock) and *γ* (the effect of the clock on phenotype) were estimated; for binary phenotypes, *γ* is on the log odds ratio scale. (A) Type-I error rate and statistical power, measured as the proportion of simulation repeats with P < 0.05. The red dashed line marks the 5% level. (B) Boxplots of the effect size estimates across replicates. Results for a broader range of settings, including the use of an external DNAm covariance matrix, are shown in **Supplementary Figures 2-4**.

### SARDINE produces robust association estimates from EWAS on four complex phenotypes

We next evaluated SARDINE using EWAS of real phenotypes in the TruDiagnostic cohort (N = 15,677; **Methods**). We included 38 epigenetic clocks in the analysis, spanning multiple model categories defined by their training targets: chronological age, gestational age, aging biomarkers and mortality, pace of aging, cumulative cell division, socioeconomic status, and specific complex traits and diseases (**Supplementary Table 1**). We performed EWAS on two continuous phenotypes, height (N = 14,578) and body-mass index (BMI; N = 14,133), and two binary phenotypes, anxiety (N = 1,965 cases and 13,712 controls) and high blood pressure (N = 2,312 cases and 13,365 controls), to produce CpG-level DNAm-phenotype association summary statistics. We also estimated DNAm covariance using the same EWAS sample. We then benchmarked SARDINE estimates of epigenetic clock-phenotype associations against those obtained from the conventional individual-level data (**Supplementary Table 2**).

Consistent with our simulation results, we obtained near-identical epigenetic clock-phenotype associations estimated from individual-level data and summary statistics (**Figure 3A-B** and **Supplementary Figure 5**). Unsurprisingly, adjusting for sex, age, age-squared, and cell type composition substantially altered the association results, highlighting the importance of controlling for potential confounding variables in epigenetic clock applications (**Supplementary Table 3** and **Supplementary Figure 6**). Importantly, when we controlled for covariates in both EWAS and DNAm covariance (**Methods**), SARDINE again successfully recovered clock-phenotype associations, showing highly consistent results compared to estimates obtained from individual-level data (**Figure 3C-D** and **Supplementary Figure 7**).

**Figure 3.**
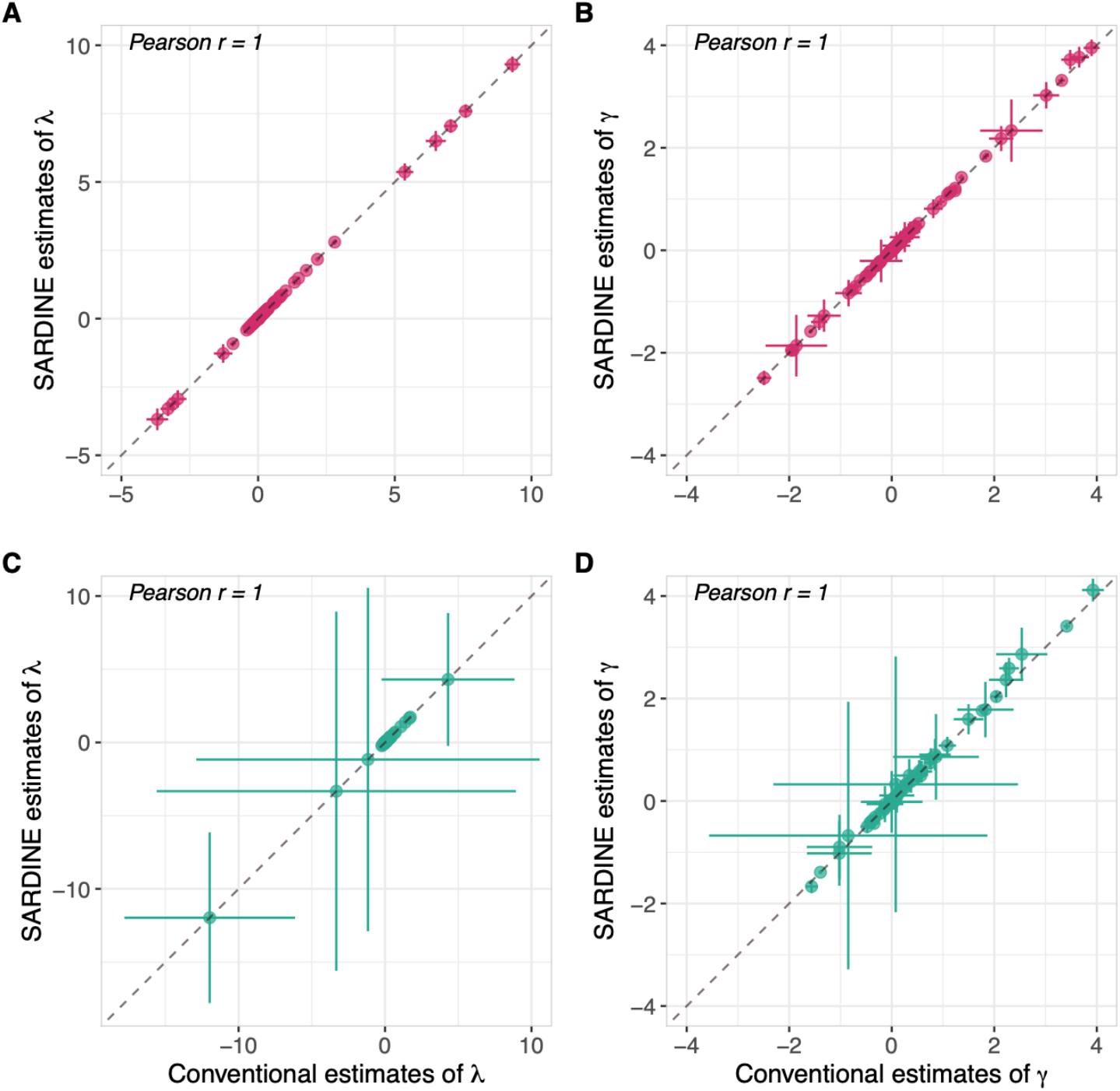
Validating SARDINE estimates of epigenetic clock-phenotype associations using four EWAS. Associations of 38 epigenetic clocks with 4 phenotypes were estimated using both SARDINE and a conventional regression approach. (A) Estimates of *λ* without adjusting for covariates. (B) Estimates of *γ* without adjusting for covariates. (C) Covariates-adjusted estimates of *λ*. (D) Covariates-adjusted estimates of *γ*. In all panels, each dot represents a clock-phenotype pair, error bars denote +/-one standard error, and the dashed line indicates y = x. The regression coefficients with extreme values are omitted from the (A) and (B) for better visualization. Covariates include sex, age, age-squared, and cell type composition. The complete results are presented in **Supplementary Figures 5** and **7** and **Supplementary Tables 2** and **3**.

Although we provide a solution for covariates-adjusted inference in SARDINE, the approach is less flexible compared to the conventional regression analysis which can easily adjust any covariates based on the scientific question. In practice, given the summary statistics of an EWAS, a reasonable and achievable goal for SARDINE would be to obtain clock-phenotype associations while controlling for the same covariates included in this EWAS. To achieve this, the DNAm covariance must adjust for the same set of covariates (**Methods** and **Supplementary Note**). This creates a challenge in practice, because some EWAS covariates could be missing in the reference dataset used for DNAm covariance estimation. To empirically assess this issue, we applied SARDINE to EWAS of height, BMI, anxiety, and high blood pressure which controlled for sex, age, age-squared, and cell type composition, in conjunction with a DNAm covariance matrix that only adjusted for a subset of these covariates. We found that SARDINE’s point estimate for *λ* is not affected by covariate adjustment in the DNAm reference, and partially-adjusted SARDINE results for both *λ* and *γ* showed lower statistical power compared to fully-adjusted results (**Supplementary Figures 8-9**). This provides an important guidance for the practical use of our method. If some covariates are missing in the DNAm reference dataset, we expect SARDINE to yield conservative inferences with reduced statistical power in the association results, but not false positive findings.

Next, we evaluated the impact of sample size and mis-matched demographics in the DNAm reference dataset (**Methods**). Clock-phenotype association estimates showed negligible changes as the reference sample size increased from 100 to 2,000 (**Supplementary Figures 10-11**). We also selected non-representative reference samples based on sex, age, and ancestry. The association estimates remained highly consistent given a sex- and ancestry-biased reference sample. However, we found biases in association results when the DNAm reference sample substantially differ from the EWAS sample in chronological age. Such biases were effectively removed when covariates were adjusted for in both the EWAS and the reference (**Supplementary Figures 12-13**). Overall, when confounders were appropriately adjusted for, the reference panels drawn from different demographic subgroups provided comparable results.

### Benchmarking epigenetic clock associations with chronological age and smoking

Having established that SARDINE produces robust association results from summary statistics, we next applied our method to publicly available EWAS data resources. We focused on chronological age and smoking, two phenotypes that are known to correlate with substantial DNAm changes across CpGs. Specifically, we collected summary statistics from four EWAS on age^8,35–37^, three of which used chronological age as a continuous variable in EWAS (but DNAm was measured in different tissues, i.e., whole blood, breast, and colorectal tissues). The fourth compared nonagenarians with younger adults (aged 19-30 years; median 22.5) in peripheral blood mononuclear cells. For smoking, we used four EWAS from two studies, comparing various categories of smokers and non-smokers in whole blood^38^ and peripheral blood mononuclear cells^39^. Summary statistics were obtained from EWAS Catalog re-analyses adjusted for batch and cell type composition. Further details on these public EWAS are in **Methods** and **Supplementary Table 4**.

Despite modest sample sizes (111 to 656; **Figure 4A** and **Supplementary Table 4**), these EWAS showed substantial clock-age and clock-smoking associations (**Figure 4B** and **Supplementary Table 5**). We identified 49 significant clock-age associations at FDR < 0.05, of which 42 remained significant after Bonferroni correction. For smoking, we found 45 associations at FDR < 0.05, of which 29 remained significant after Bonferroni correction (multiple testing corrections were applied across the 38 clocks x 4 EWAS = 152 tests for each phenotype).

**Figure 4.**
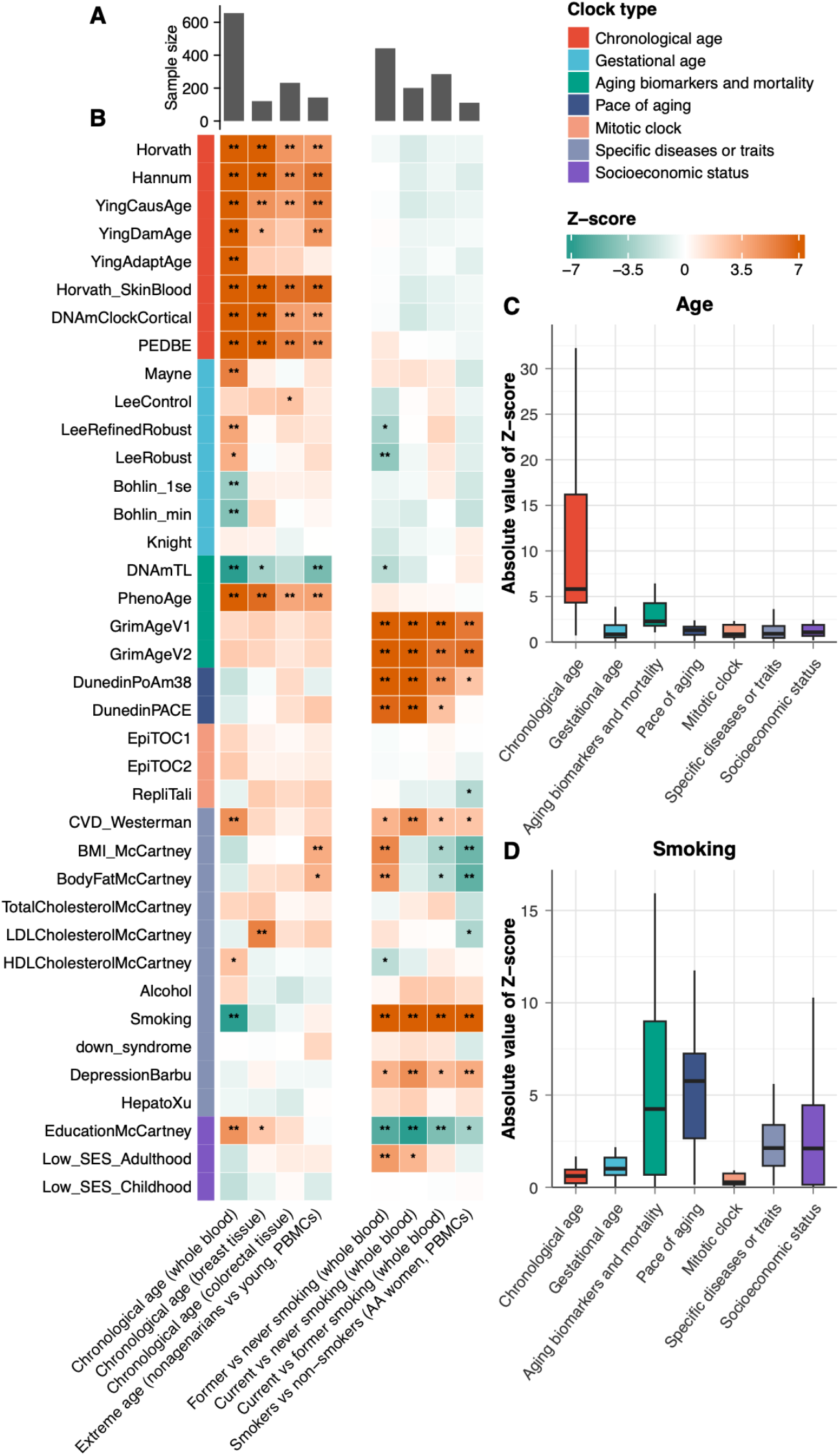
Associations of 38 DNAm clocks with chronological age and smoking. (A) Sample size of each EWAS. (B) Heatmap of clock associations with chronological age across four studies, and with four smoking outcomes in two studies. Color indicates the sign and magnitude of the association Z-scores in the heatmap. One asterisk denotes FDR<0.05 and two asterisks denote significance after Bonferroni correction. Z-scores are truncated at 7 and −7 to improve visualization. Full association results are shown in **Supplementary Table 5**. Boxplots of the absolute value of Z-scores for clock associations with chronological age (C) and smoking (D), grouped by clock category.

As shown in prior research, age and smoking were associated with largely non-overlapping groups of epigenetic biomarkers. Clock models trained on chronological age showed the strongest associations with age in our analysis (**Figure 4C**). Six out of eight chronological age clocks showed significant age associations in all four EWAS. These associations were the strongest in whole blood, which may be attributed to the larger EWAS sample size, but were also identified in breast- and colorectal-tissue EWAS. Clock models trained in adult samples (i.e., Hannum, and YingCausAge), pediatric samples (i.e., PedBE), and various tissue types (i.e., brain cortex in DNAmClockCortical and multiple tissues in Horvath clocks) all showed significant and robust associations with chronological age. In contrast, no pace-of-aging or mitotic clocks were significantly associated with chronological age in these smaller EWAS, although both showed significant associations in the larger TruDiagnostic sample (**Supplementary Figure 14** and **Supplementary Table 6**).

Smoking showed a different association pattern. Unsurprisingly, among all biomarker models included in the analysis, the DNAm smoking score, which was trained to predict smoking pack-years, showed the strongest associations with smoking. Additionally, six other models (i.e., GrimAge V1 and V2; DunedInPoAm38; Westerman_CVD; EducationMcCartney; and DepressionBarbu) showed significant associations at FDR < 0.05 in all four smoking EWAS. The associations of GrimAge V1 and V2 with smoking were expected by construction: both scores are weighted sum of DNAm-based surrogate biomarkers, one of which (“DNAm PACKYRS”) was trained to predict self-reported smoking pack-years. Overall, pace of aging clocks and clocks trained on mortality showed much stronger associations with smoking compared to other biomarker categories (**Figure 4D**). No chronological age clock or mitotic clock showed any significant associations with smoking after Bonferroni correction.

Some of these associations need to be interpreted with caution. The whole blood age EWAS dataset (i.e., GSE40279) was used for CpG selection or model training for Hannum, Horvath, EpiTOC1, EpiTOC2, and HepatoXu clocks. Such sample overlap can cause overfitting and inflate association results. Further, GrimAge clocks are functions of chronological age and sex in addition to DNAm, where SARDINE uses only their DNAm component. When the EWAS adjusts for age and sex, SARDINE estimates are identical to those from the full clock and correspond to age acceleration effects of the GrimAge clocks^10,40^. When the EWAS does not adjust for age or sex, as in the age and smoking EWAS in **Figure 4** which adjusted only for the estimated proportions of 12 cell types, the SARDINE estimates reflect only the DNAm component. They can therefore differ from estimates obtained by analyzing the full clock with individual-level data. We discuss these issues further in the **Supplementary Note**.

Due to these potential technical issues, we replicated clock-age and clock-smoking associations in the independent TruDiagnostic sample (**Methods**; **Supplementary Table 6**). We found much stronger and more significant associations across many models due to the larger sample size, but the overall association pattern with various clock model categories persisted (**Supplementary Figure 14**). Chronological age clocks showed strongest associations with age, while clocks trained on smoking, mortality, and pace of aging showed stronger associations with cigarette smoking.

### Epigenetic clocks are associated with prevalent and incident disease outcomes

We applied SARDINE to public EWAS summary statistics of 14 incident and 10 prevalent disease outcomes from Generation Scotland^33^, each with at least 100 cases (**Supplementary Table 7**). Prevalent outcomes were conditions present at the baseline blood draw. Most were self-reported on the baseline health questionnaire on participants, except Alzheimer’s disease was self-reported parental history and chronic kidney disease status was defined based on the estimated glomerular filtration rate at baseline. Incident outcomes were first diagnoses after the baseline ascertained through linkage to primary- and secondary-care records. Prevalent and incident outcomes were analyzed by logistic and Cox regression, respectively, with DNAm on the M-value scale measured at baseline; SARDINE accommodates both specifications (**Methods** and **Supplementary Note**). A recent study has assessed the performance of 14 epigenetic clocks using many disease outcomes in the same cohort^20^. Here, using only publicly available EWAS summary statistics, we evaluated 29 epigenetic biomarkers. Nine biomarkers of our original 38 were trained on the Generation Scotland sample and were thus excluded from these analyses to avoid overfitting (**Methods**). Age, sex (when applicable), batch, and estimated cell type composition were adjusted for in these EWAS. The reported associations are therefore between each pair of phenotype and clock conditioning on these covariates.

We identified 129 significant clock associations with prevalent disease outcomes at FDR<0.05, of which 83 remained significant after Bonferroni correction (**Figure 5A** and **Supplementary Table 8**). Similarly, 155 associations were found at FDR<0.05 for incident disease outcomes and 98 remained significant after Bonferroni correction (**Figure 5B** and **Supplementary Table 9**). Both corrections were applied jointly across 696 tests for 24 outcomes x 29 clocks. For the 10 diseases analyzed in both prevalent and incident outcomes, clock association Z-scores were highly concordant (Pearson correlation r = 0.81; **Figure 5C**).

**Figure 5.**
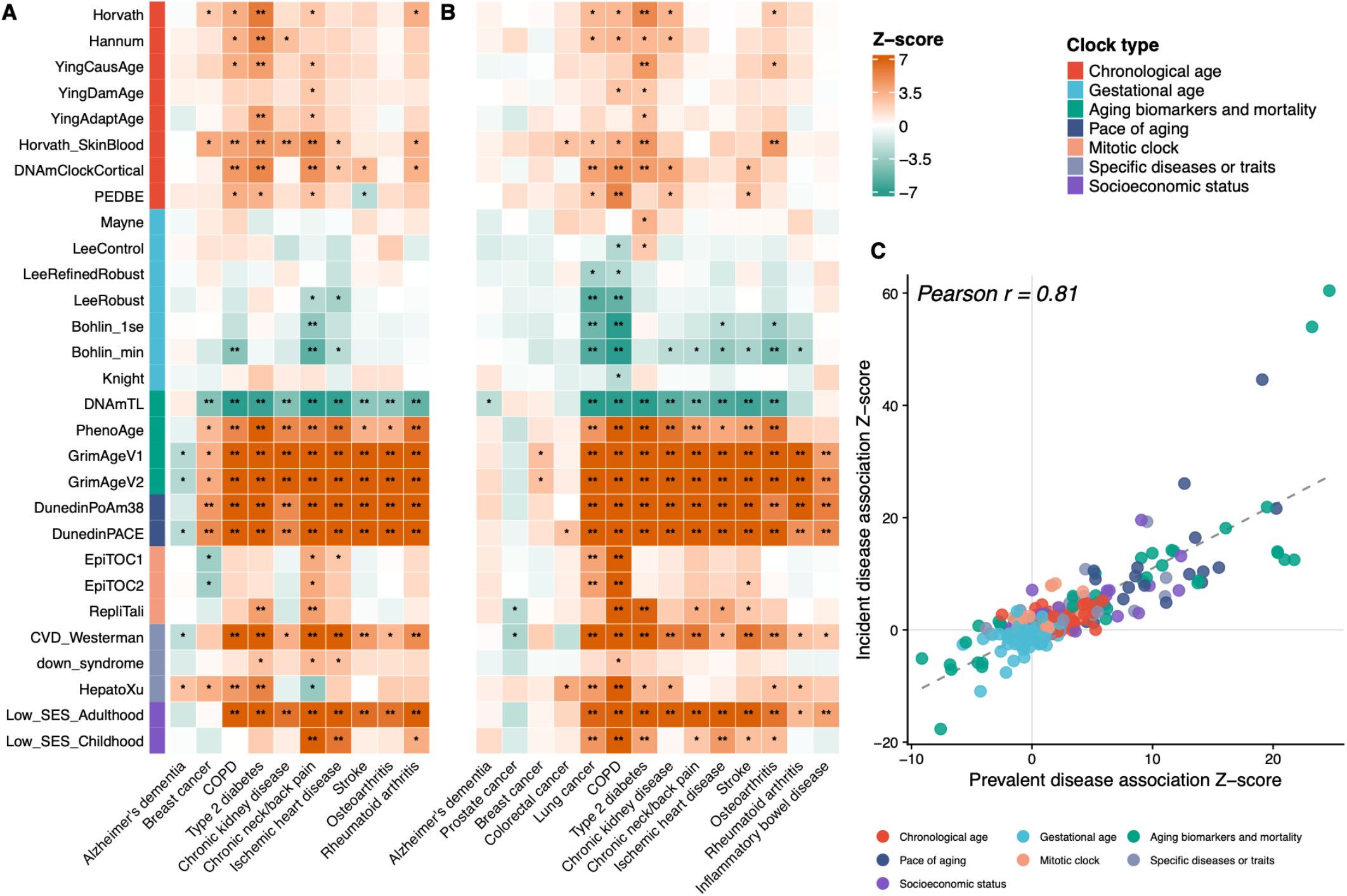
Associations of 29 DNAm clocks with 10 prevalent and 14 incident disease outcomes in Generation Scotland. Heatmap of clock associations Z-scores with 10 prevalent (A) and 14 incident (B) disease outcomes. Color indicates the sign and magnitude of the Z-score. One asterisk denotes FDR<0.05 and two asterisks denote significance after Bonferroni correction. Z-scores are truncated at 7 and −7 to improve visualization. Full association results are in **Supplementary Tables 8** and **9**. (C) Scatter plot comparing association Z-scores for 10 diseases analyzed in both prevalent and incident outcomes. The dashed line represents the fitted linear regression.

We found substantially different disease association patterns across biomarker categories. Epigenetic clocks trained on aging phenotypes, biomarkers, or mortality and pace of aging clocks were associated with most disease outcomes, while chronological age clocks showed substantially weaker associations (**Supplementary Figure 15**). These findings are consistent with previous results produced from individual-level data analysis^20^. Chronic obstructive pulmonary disease (COPD) and type 2 diabetes stood out as exceptions: six of the eight chronological age clocks were associated with COPD, and seven with type 2 diabetes, at FDR < 0.05 in each of the prevalent and incident analyses. Smoking was not adjusted for in these disease EWAS and may contribute to the COPD associations with smoking-related clocks such as GrimAge. However, unadjusted smoking is unlikely to explain the COPD associations with chronological age clocks which showed little association with smoking (**Figure 4** and **Supplementary Figure 14**). We also note that epigenetic biomarkers trained on specific disease outcomes lacked specificity in their prediction target. For example, the Westerman CVD epigenetic biomarker was strongly associated with most prevalent and incident disease outcomes in our analysis. Its associations with incident COPD (Z = 19.3) and type 2 diabetes (Z = 9.3) were stronger than its association with incident ischemic heart disease (Z = 3.4). Similarly, HepatoXu clock, which was trained to distinguish patients with hepatocellular carcinoma (the predominant form of liver cancer) from healthy controls, showed significant associations with COPD and diabetes in addition to cancer outcomes.

Compared to cardiovascular, pulmonary, immune, and metabolic disease outcomes, some cancers (e.g., incident prostate, breast, and colorectal cancers) and Alzheimer’s disease (AD) showed weaker associations with epigenetic clocks. This cannot be fully explained by lower sample size on these outcomes. For example, incident breast, prostate, and colorectal cancer and incident AD had case counts comparable to or larger than incident rheumatoid arthritis. Yet rheumatoid arthritis showed substantial associations with GrimAge and pace-of-aging clocks despite having the lowest incident case counts (N = 120) in our analysis (**Supplementary Table 7**). Mitotic clocks were developed for predicting the cumulative number of stem cell divisions related with cancer risk. However, apart from incident lung cancer, they showed no significant positive associations with cancer outcomes. Similarly, the cortical clock, trained on brain DNAm, did not show associations with either prevalent or incident AD. This suggests that future biomarker development is still needed in order to capture the DNAm signatures underlying the etiology and progression of these important diseases.

Finally, we note that phenotype quality is an important factor that can affect these analyses. AD parental history, instead of own AD diagnosis, was used as the prevalent AD outcome in EWAS (**Supplementary Table 7**). This is a common approach in genetic epidemiological studies leveraging mid-aged cohorts^41^, but is also known to cause complex biases in association analysis^42^. In fact, AD was the only disease showing discordant epigenetic clock associations in prevalent and incident outcome analyses (Pearson *r* = −0.31; **Supplementary Figure 16**). The FDR-significant yet paradoxical associations between AD (parental history) and slower epigenetic aging (e.g., quantified by GrimAge, DunedinPACE, and CVD clocks) are likely driven by biases in offspring-reported parental diagnosis^42^.

### Quantifying epigenetic clock associations with modifiable lifestyle factors

Finally, we investigated how lifestyle factors are associated with epigenetic aging biomarkers (**Figure 6**). We performed EWAS on six lifestyle phenotypes (drinks per week, coffee per week, sleep hours, exercise per week, fish oil intake, and herbal supplement intake), while controlling for sex, age, age-squared, and cell type composition using the TruDiagnostic sample (**Methods**). We applied SARDINE to the summary statistics of these EWAS, identifying 104 clock-phenotype associations at FDR<0.05. Sixty of these associations remained significant after Bonferroni correction (**Supplementary Table 10**). We also reproduced these associations using a conventional regression approach, and obtained virtually identical results compared to SARDINE estimates (**Supplementary Table 10** and **Supplementary Figure 17**).

**Figure 6.**
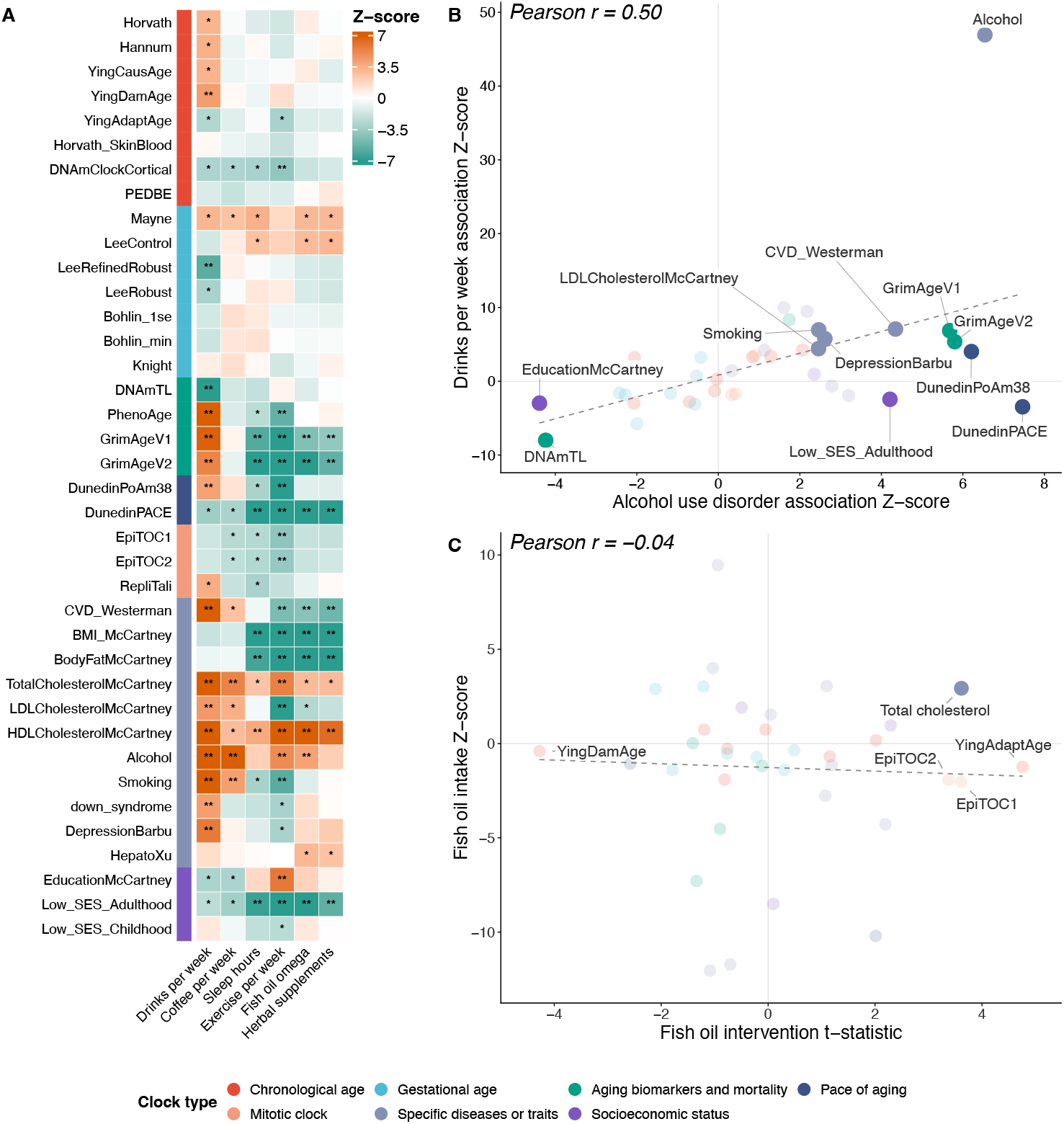
Associations of epigenetic clocks with lifestyle factors. (A) Heatmap of associations Z-scores between 38 clocks and 6 lifestyle factors in the TruDiagnostic cohort. Color indicates the sign and magnitude of the Z-scores. One asterisk indicates statistical significance at FDR<0.05 and two asterisks denote significance after Bonferroni correction. Z-scores are truncated at 7 and −7 to improve visualization. Full association results are in **Supplementary Table 10**. (B) Scatter plot comparing association Z-scores for alcohol use disorder versus drinks per week. Clocks reached FDR<0.05 in both analyses are highlighted and labeled. (C) Scatter plot comparing fish-oil intervention t-statistics with observational fish-oil intake association Z-scores across 36 epigenetic clocks. Clocks reached FDR<0.05 in both analyses are highlighted and labeled, clocks reached FDR<0.05 only in fish oil intervention are labeled. In (B) and (C), the dashed line represents the fitted linear regression.

The identified associations were concentrated in epigenetic measures targeting aging biomarker/mortality, pace of aging, socioeconomic status, and complex traits. One epigenetic aging measure and three epigenetic surrogates or biomarkers (DunedinPACE, TotalCholesterolMcCartney, HDLCholesterolMcCartney, and Low_SES_Adulthood) were associated with all six lifestyle phenotypes. Notably, the cortex clock derived from brain DNAm samples was the only chronological age clock showing broader associations across lifestyle factors. Exercise, sleep hours, fish oil intake, and herbal supplement intake showed similar associations with decelerated epigenetic aging. In contrast, drinks per week generally showed strong associations with increased DNAm age, including biomarkers predicting chronological aging. The McCartney alcohol consumption DNAm biomarker showed highly significant association (Z = 46.95) with drinks per week; it was nonetheless also associated with coffee consumption, exercise, and fish oil intake indicating limited specificity of complex trait epigenetic predictors and possibly revealing socioeconomic confounding in these associations.

We sought replication of the association findings for alcohol consumption using a publicly available EWAS on alcohol use disorder (AUD)^43^ (**Figure 6B** and **Supplementary Table 11**). Across the 38 epigenetic clocks, their association with two phenotypes (i.e., drinks per week and AUD) were moderately correlated (Pearson r = 0.50). Twelve epigenetic measures showed significant associations at FDR < 0.05 in both analyses; of these, ten showed concordant directions whereas DunedinPACE and Low_SES_Adulthood showed opposite directions. Differences in phenotype definition and cohort sample selection may contribute to these discrepancies. Overall, these results suggest partial consistency of alcohol-related DNAm clock associations in two independent datasets.

Our results suggested that self-reported fish oil intake is associated with healthy aging, which is quantified by numerous epigenetic clocks (e.g., decelerated DunedinPACE, Z = −10.2, p = 1.88e-24; **Supplementary Table 10**). A reasonable hypothesis derived from these associations is that consuming fish oil could be causally beneficial, leading to decelerated aging. To assess this, we sought replication of clock-phenotype associations in an intervention study^44^ focusing on the effect of 6 weeks of fish oil supplementation on DNAm in overweight and obese individuals, using 33 paired pre- and post-intervention samples. RepliTali and down syndrome were excluded from this comparison because only 30/87 (34.5%) and 317/652 (48.6%) of their CpGs were found in the intervention dataset, respectively.The associations of fish oil intake with various epigenetic clocks showed low concordance between the observational and intervention studies (Pearson r = −0.04; **Figure 6C**), reminding us of an old but nevertheless important statistical lesson: cross-sectional associations cannot be interpreted as causal evidence. Only one epigenetic biomarker, i.e., TotalCholesterolMcCartney, showed significant associations at FDR < 0.05 in both analyses (**Supplementary Table 12**).

## Discussion

Most epidemiological studies of epigenetic aging investigate associations between human phenotypes and the deviation of biological age as estimated by epigenetic clocks and chronological age. Conventionally, these analyses require individual-level DNAm data to construct clock measures and estimate their associations with phenotypes. Here, we introduced SARDINE, a summary-statistics-based framework that overcomes this requirement and enables scalable association analysis of DNAm-based biomarkers. Our approach makes two important contributions. First, the rapid proliferation of epigenetic clocks and related biomarkers has created an urgent need for systematic evaluation and benchmarking. SARDINE transforms publicly available yet largely underutilized EWAS resources into a powerful platform for comparing established and emerging biomarkers across diverse phenotypes. Second, SARDINE provides important tools for post-EWAS analysis, translating CpG-level associations into interpretable clock-level estimates that can be compared across models, phenotypes, and cohorts. By eliminating the need for individual-level data, SARDINE breaks important barriers to data access and computational scalability, enabling analyses that would otherwise be difficult or infeasible.

Several important features support the broad applicability of SARDINE. The framework estimates two types of associations: (i) the effects of exposures on epigenetic clocks and (ii) the effects of epigenetic clocks on downstream outcomes. It accommodates continuous, binary, and survival EWAS outcomes, DNAm measured on either beta- or M-value scale, and EWAS that treat DNAm as either the outcome or predictor. SARDINE also supports covariate-adjusted inference when the reference DNAm covariance matrix is adjusted for the same covariates as the EWAS. When some covariates are unavailable in the reference panel, our empirical results suggest that partially adjusted analyses can yield conservative inference results with reduced statistical power. Furthermore, we demonstrate that relatively small reference panels of a few hundred individuals can provide robust association estimates, even when their demographic characteristics differ from those of the EWAS cohort, provided that relevant confounders are appropriately controlled. Together, these features make SARDINE a flexible and practical tool for large-scale epigenetic aging research.

Employing SARDINE to public and restricted access EWAS datasets, we identified numerous associations between DNAm-based biomarkers and human phenotypes. There are several key take-aways from these association findings. First, epigenetic predictors developed for different purposes exhibit substantially divergent phenotypic associations. As shown in prior research, chronological age clocks strongly tracked age, whereas pace-of-aging and mortality clocks were more strongly associated with smoking, disease outcomes, and lifestyle factors. Second, epigenetic predictors trained on specific complex traits often showed broad associations with other phenotypes, raising questions about their biological specificity. For example, the HepatoXu biomarker, developed to predict hepatocellular carcinoma, showed strong associations with COPD. Similarly, a DNAm-based alcohol consumption predictor was associated not only with alcohol intake but also with coffee consumption, exercise, and fish oil supplementation. These findings may reflect shared DNAm signatures across phenotypes, but could also arise from confounding factors such as smoking, socioeconomic status, and correlated behaviors. Third, our analyses identified important gaps in existing biomarkers. No evaluated clock showed a Bonferroni-significant association with incident colorectal cancer, breast cancer, prostate cancer, or Alzheimer’s disease, though these are all diseases for which blood would be peripheral and not the main tissue of interest. An important future direction is to investigate whether these diseases have unique epigenetic signatures not effectively captured by current models, which would suggest that new biomarkers are needed for these important health outcomes. Finally, our findings highlight the important lesson that strong cross-sectional associations do not always translate into causal effects. Although self-reported fish oil intake was associated with slower epigenetic aging across several biomarkers, these associations showed little concordance with results from a short-term intervention study. Study population (overweight and obese individuals), smaller sample size, and short study duration (6 weeks) of the interventional study may contribute to this discrepancy^44^. More recent, comprehensive trials have revealed some causal effect of longer-term fish oil consumption on healthy aging, especially in conjunction with other lifestyle interventions including exercise and vitamin D supplementation^45^. Nevertheless, these findings underscore the need for rigorous causal evaluation before translating biomarker associations into interventions targeting healthy aging.

Perhaps most importantly, this study has broader implications for how epigenetic aging research is designed and conducted. Over the past 15 years, the field of complex trait genetics has been transformed by large-scale genome-wide association study (GWAS) meta-analyses and the widespread sharing of summary statistics^46^. These resources have enabled the development of statistical methods for causal variant fine-mapping^47^, variance component analysis^48,49^, and genetic risk prediction^50^, substantially accelerating discovery without requiring individual-level genetic data. Our study demonstrates that a similar summary-statistics-based paradigm can also benefit biological aging research. We envision a future in which releasing full summary statistics after performing EWAS becomes a routine in the field. Increasing the accessibility of EWAS summary data resources will enable consortium-scale meta-analysis and improve the statistical power and reproducibility of association results. Importantly, SARDINE illustrates how CpG-level association results, even when individually underpowered, can be integrated into biologically interpretable biomarker-level analyses. Further development of summary-statistics-based methods could substantially expand the scientific value of existing and future EWAS resources. Finally, the underlying statistical framework of SARDINE is applicable to any biomarker that is defined as a weighted sum of molecular features, so the framework naturally extends to other types of clock models including proteomic^51^, transcriptomic^3^, and histological^52^ aging clocks, although marginal omic variable-phenotype association analysis is not yet the current norm in these other areas of biological aging research.

Our study has some limitations. First, SARDINE is not robust to sample overlap between the training cohort for clock models and the EWAS study cohort. Such overlap can lead to overfitting which will inflate association estimates. Second, SARDINE requires DNAm-based predictors that can be represented as weighted sums of CpG methylation measurements. Non-linear models (e.g., MAPLE^53^) or models based on order statistics (e.g., stemTOC^54^) are therefore not directly supported. Although principal component (PC) clocks^55^ satisfy the linearity assumption, their large numbers of CpGs can impose substantial computational demands. Therefore, we have also excluded them from the analyses. For biomarkers that incorporate additional covariates or nonlinear transformations, SARDINE estimates associations with the weighted sum component, which may differ from associations with the final reported biomarker. Third, when in-sample DNAm covariance is not available, SARDINE relies on an external reference panel to approximate the DNAm covariance in the EWAS dataset. Systematic differences between reference panels and EWAS populations, particularly those not adequately addressed through covariate adjustment, may introduce bias (**Supplementary Figures 12-13**; **Supplementary Note**). Further, differences in tissue type, cellular composition, experimental batch, and DNAm processing may influence covariance between CpG sites and consequently affect association estimates and their uncertainty. Although our sensitivity analyses demonstrated robustness to differences in reference sample size and demographic composition, these analyses primarily used reference panels derived from the same cohort and processing pipeline. Therefore, careful selection and evaluation of reference panels remain important for reliable inference.

Taken together, this work addresses a fundamental methodological bottleneck that currently limits the application, validation, and interpretation of epigenetic aging biomarkers. The methodological innovations unlock the full value of publicly available EWAS resources and enable well-powered and comprehensive studies that are currently infeasible. We expect the statistical framework and software produced from our study to provide the aging research community with paradigm-shifting analytic tools to accelerate discovery and improve understanding of the biological mechanisms that shape human aging.

## Methods

### EWAS model

We consider a common model for EWAS:

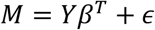

Here, *M* is an *N* × *J* matrix containing DNAm measurements for *N* individuals across *J* CpG sites, *Y* is an *N* × 1 column vector containing each individual’s phenotype value, *β*^*T*^ = (*β*_1_, *β*_2_, …, *β*_J_) parameterizes the effect sizes of *Y* on DNAm at *J* CpG sites, and *ϵ* is an *N* × *J* matrix containing random noise with Var 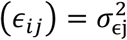. We assume that *Y* and each column of *M* are centered at 0. Noise *ϵ* and phenotype *Y* are also assumed to be independent.

For simplicity, here we introduce our statistical formulation using a setting where DNAm is the outcome in EWAS. This is a common design for studying the effect of an exposure (e.g., smoking) on DNAm. We note that generalizing this framework to a setting in which *Y* is the outcome is straightforward (**Supplementary Note**). Depending on the focus of an EWAS, *Y* may represent any variable ranging from socioeconomic conditions, lifestyle factors, environmental exposures, disease, or time-to-event outcomes. In this work, we broadly refer to all of these as phenotypes, and *β* denotes the true association between a phenotype with DNAm at CpG sites.

The OLS estimate for the j-th CpG is

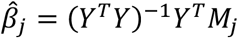

and the variance of 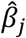 can be estimated as

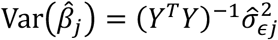

where 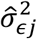 is the mean squared error obtained from regression. Point estimate 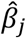 and its variance Var 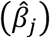are typically available in the EWAS summary statistics.

### Methylation clock models and DNAm covariance matrix

Most existing methylation clock models can be denoted as a weighted sum of DNAm across CpG sites:

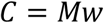

where *w* = (*w*_1_, *w*_2_, …, *w*_J_)^*T*^ denote the CpG weight values in the clock model. Typically, these weight values are publicly released alongside the paper that introduces the model. If a CpG site is not included in a model, we assign the corresponding weight *w*_*J*_ to be 0.

We note that some epigenetic clock models also include an intercept term. It is a constant across individuals and does not affect clock-phenotype association inference. Therefore, for simplicity, we ignore the intercept in our derivations below. Some clocks contain contributions from sex and age; when both sex and age are included in the covariates, ignoring them from the clock construction does not affect clock-phenotype association inference (**Supplementary Note**).

Another input to SARDINE is the *J* × *J* variance-covariance matrix for DNAm across CpG sites:

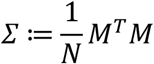

Ideally, this matrix would be obtained from the EWAS study sample. However, since individual-level EWAS data are often unavailable in practice, we can use an external reference DNAm dataset to approximate *Σ*:

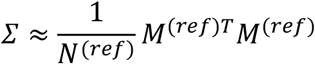

Here, *M*^(*ref*)^ is the DNAm matrix from the reference dataset and *N*^(*ref*)^ is its sample size.

### Estimating the effect of a phenotype on epigenetic aging

A common application of an epigenetic aging clock is to use it as an outcome biomarker to quantify the effect of a phenotype (e.g., smoking, trauma, exposure to toxicants, economic hardship, anti-aging intervention) on biological aging. This is commonly achieved through estimating *λ* in the following regression

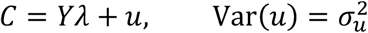

The OLS estimate for *λ* is

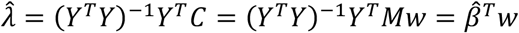

An important observation here is that 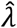, which is the estimated effect of a phenotype *Y* on an epigenetic clock *C*, can be obtained from a weighted sum of CpG-level associations (i.e.,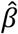) in an EWAS of *Y*. This is how our approach bypasses the requirement of individual-level data, and estimating clock-phenotype associations from EWAS summary statistics.

The variance of 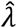 is

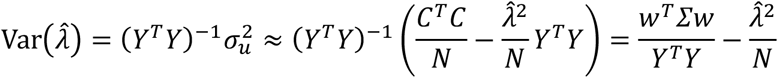

Here, in order to be free of any dependence on individual-level EWAS data, we need an approach for estimating Var(*Y*) (or equivalently, *Y*^*T*^*Y*) without using the observed values of *Y*. Recall that in the EWAS model, we had

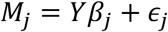

Given the independence of *Y* and *ϵ*_*J*_, we have

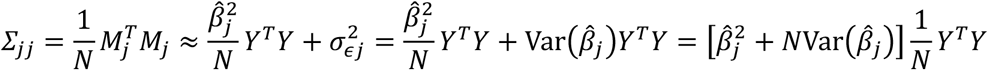

Thus, Var(*Y*) can be approximated by

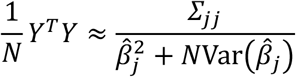

That is, we can estimate Var(*Y*) from diagonal terms of the DNAm covariance matrix *Σ*_*JJ*_, point estimate 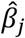 in an EWAS, and its variance Var 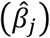 which are available in EWAS summary statistics. Further, any arbitrary CpG j can provide an estimate for Var(*Y*). For numerical stability, we use the median estimate across CpG sites:

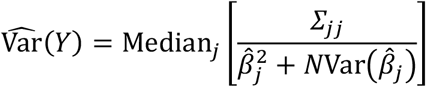

Based on this, Var 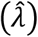 can be estimated as

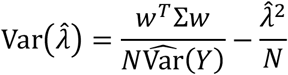

Hypothesis testing for *λ* = 0 can be performed using test statistic

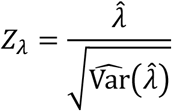

which allows for a standard approach for two-sided *P* value calculation.

The important insight is that statistical inference on the effect of a phenotype on an epigenetic clock can be performed without individual-level data. The point estimate 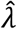 is the inner product of the pre-trained clock model weights and EWAS effect estimates across the CpGs. Its variance can also be approximated using EWAS summary statistics and covariance matrix *Σ*.

### Estimating the effect of epigenetic aging on a phenotype

Another common application is to estimate the effect of an epigenetic aging clock on an outcome of interest (e.g., disease diagnosis, complex traits, and other types of aging biomarkers). This is achieved through the following regression

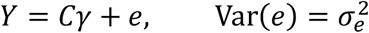

The OLS estimate of *γ* is

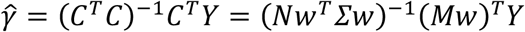

Recall that EWAS estimates can be denoted as 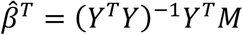, therefore

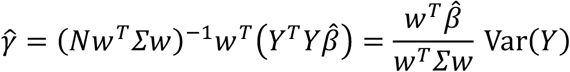

Thus, we can also approximate the OLS estimate 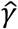, with EWAS summary statistics, clock model weights *w*, and DNAm covariance matrix *Σ*.

Similarly, we can estimate the variance of 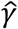,

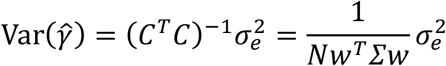

where 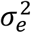 can be estimated as

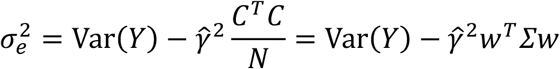

Combining these formulas, we obtain the estimate for Var 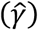

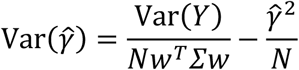

P-value calculation and hypothesis testing can be based on the following test statistic.

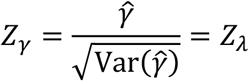

Detailed discussions regarding covariate adjustments, DNAm in M-value, and the incorporation of EWAS summary statistics derived from alternative models such as regressing phenotype *Y* on *M*, logistic regressions and Cox proportional hazards models can be found in the **Supplementary Note**. Another technical note is that SARDINE assumes the epigenetic clock to be a weighted sum of CpGs. For clocks whose published algorithm adds additional terms (e.g., GrimAge) or transformations (e.g., Horvath, PEDBE, DunedinPACE) to a weighted sum of DNAm, SARDINE estimates associations with the weighted sum component, and the estimates should be interpreted on that scale. Since GrimAge clocks have additional contributions from sex and age, in the Supplementary Note, we showed that SARDINE’s estimates are unbiased when sex and age are included in the clock-phenotype regression covariates. For Horvath, PEDBE, and DunedinPACE clocks, empirically we found that SARDINE’s estimates are highly concordant with those based on individual-level data where the clock was computed allowing for the non-linear transformations (**Supplementary Figures 18-21**).

### TruDiagnostic data details

The TruDiagnostic cohort is a US-based commercial cohort. Participants provided blood samples for epigenetic testing and completed a survey on their demographics, health, and lifestyle. DNA methylation was assayed with Illumina Infinium Methylation EPIC v1 array and the data that passed quality control (QC) was provided by TruDiagnostic. Detailed QCs have been reported previously^56^. 16,051 samples have DNAm data and 15,677 passed TruDiagnostic’s internal phenotype QCs.

### Estimation of cell-type compositions

We estimated blood cell-type proportions from the DNAm data using the Robust Partial Correlation (RPC) method implemented in the EpiDISH R package^57^. Specifically, we estimated a 12-cell-type composition using the “cent12CT.m” reference, and a 19-cell-type composition using the “centUniLIFE.m” reference. To avoid collinearity in the regression analyses, we excluded cell types that exhibited little variation or were the most abundant reference cell type. Therefore, we excluded “Eosinophils” (sparsity) and “Neutrophils” (most abundant), which are both as expected for adult whole blood samples, when adjusting for 12 cell-type compositions.

### DNAm references

To establish external reference panels for DNAm variance-covariance estimation, we computed a baseline matrix using a randomly selected subset of 2,000 individuals from the TruDiagnostic cohort. These individuals self-identified as having European or Caucasian ancestry and reported their biological sex as male or female. To evaluate the effect of reference sample size, we generated additional matrices from randomly down-sampled subsets of 1,000, 500, and 100 individuals from this baseline group. To assess demographic and ancestry effects, we generated matrices from stratified subsets of the 2,000 European individuals, specifically including 500 males, 500 females, the 500 youngest, and the 500 oldest participants. For cross-ancestry comparisons, we additionally generated matrices using randomly selected subsets of 500 Asian and 500 Latino or Hispanic individuals. None of these samples used for deriving the reference panels overlap with the simulation sample.

Using each subset of individuals, we first regressed out specified covariates from the DNAm Beta-values and then used the resulting residuals to compute covariate-adjusted DNAm reference matrices. To accommodate various covariate adjustments in downstream EWAS, we computed matrices adjusting for several covariate combinations: age and age-squared alone; sex alone; 12 cell-type compositions alone; sex, age, and age-squared; and sex, age, age-squared, and 12 cell-type compositions.

To facilitate the transformation of effect sizes from M-values to Beta-values in downstream analyses, we calculated the mean DNAm Beta-value for each CpG site across the complete sample pool and within each generated subset. Empirically, we found that the mean values from the European N = 500 reference cohort demonstrated high Pearson correlations with those derived from the other subsets (**Supplementary Figure 22**).

### Simulation settings

The simulation study was conducted using a random subset (N = 8,880) of TruDiagnostic samples of self-reported European ancestry. To reduce computational burden while preserving realistic correlation structures among clock CpGs, we restricted the DNAm matrix to the 335 of 353 Horvath pan-tissue clock CpGs available in the data. Using the published clock weights, we computed a linear clock score as a weighted sum of those 335 CpGs. We standardized the clock score to have zero mean and unit variance (*C*_*std*_); and then rescaled the clock weights accordingly for use in SARDINE. In the main simulation results, the DNAm variance-covariance matrix was estimated from all the simulation samples (in-sample reference). We additionally assessed the impact of reference matrices estimated from the holdout TruDiagnostic participants not included in the simulation sample.

Continuous phenotypes were simulated as *Y* = *rC*_*std*_ + *ϵ*, where *ϵ* is an independent random noise with mean 0 and variance 1 – *r*^2^, and *r* is the target correlation between the simulated phenotype *Y* and the standardized clock *C*_*std*_. This data-generating process ensures that the true linear effect size between *Y* and *C*_*std*_ equals *r*, allowing us to establish true effect sizes of 0, 0.1, 0.2, …, 0.9. Under the null hypothesis of no association between the phenotype and the clock, the phenotype was drawn from a standard normal distribution.

Binary phenotypes were simulated using a continuous liability threshold model, where an underlying liability was first generated using the aforementioned formula and subsequently was dichotomized at a threshold corresponding to a prevalence of 30%. To achieve predetermined effect sizes on the observed binary scale, we derived the relationship between *r* (which is the correlation between the epigenetic clock and the unobserved continuous liability) and *λ* and *γ* (**Supplementary Note**). Specifically, to achieve a predetermined case-control mean difference (*λ*) in clock score, we set the liability correlation to *r* = *λK*(1 − *K*)/*ϕ*(*τ*), where *K* is the disease prevalence and *ϕ*(*τ*) is the standard normal density at the liability threshold. For a predetermined logistic log-odds ratio (*γ*), no closed-form formula exists because the liability threshold model implies a probit rather than a logistic relationship between the phenotype and the clock. The logistic regression is thus misspecified. We therefore solved numerically for the value of *r* at which the population logistic regression slope of the phenotype on the standardized clock score equals the target *γ*.

In each replicate, EWAS summary statistics were computed for each CpG by marginal regression without covariates. We ran EWAS in two directions: linear regression of the CpG beta-value on the phenotype, and linear (logistic) regression of the continuous (binary) phenotype on the CpG beta-value. SARDINE was applied to each set of summary statistics. As a benchmark, *λ* and *γ* were also estimated from individual-level data.

We performed 2,000 replicates for each setting. The type-I error rate was evaluated by calculating the proportion of false positives at a 5% nominal significance level under the null, and estimation bias was assessed by comparing the distribution of the estimated effect sizes against their true parameter values.

### EWAS details

We conducted EWAS using the EPICv1 array data from the TruDiagnostic cohort. We used all the samples whose self-reported sex is either male or female (N = 15,677). We extracted 26 phenotypes with sufficient effective sample sizes, comprising 16 binary and 10 continuous traits. For weight and body mass index (BMI), phenotypic outliers were assigned as missing values based on the interquartile range (IQR). Categorical survey responses were numerically recoded to approximate continuous or ordinal variables (**Supplementary Table 13**). For example, alcohol consumption and caffeine use were recoded into approximate weekly frequencies (ranging from 0 for “Never” to 7 for “Regularly”). Binary traits (e.g., self-reported medical conditions and supplement use) were coded as 1 for presence/affirmative and 0 for absence. Following recoding, all continuous and ordinal categorical variables (including age, weight, height, BMI, stress level, sleep hours, and substance use frequencies) were standardized prior to analysis. In the EWAS, DNAm Beta-values were regressed on each standardized phenotype while adjusting for varying sets of covariates. In simulation studies, and when examining the impact of DNAm references, we randomly selected 8,880 European samples for EWAS.

### Public EWAS of age and smoking

Public EWAS summary statistics were obtained from the EWAS Catalog^34^. For each dataset, we reviewed the corresponding publication and study design to verify phenotype coding and harmonize effect directions according to the contrasts defined in the original study. For the age EWAS, the reported effect directions were retained for continuous-age analyses conducted in whole blood, breast tissue, and colorectal tissue. For the extreme-age EWAS comparing nonagenarians with younger adults, we reversed the sign of the EWAS coefficients so that the meaning of the coefficient was consistent with the direction used in the other three age EWAS.

The same harmonization procedure was applied to the smoking EWAS. Effect directions were reversed for the former-versus-never, current-versus-never, and current-versus-former smoking contrasts to ensure consistent directionality across studies, whereas the smokers-versus-nonsmokers contrast was retained as originally reported. These public EWAS only adjusted for cell-type compositions. The whole-blood age EWAS (GSE40279) was used in developing the Hannum, Horvath, EpiTOC1, EpiTOC2, and HepatoXu clocks, for either CpG selection or model training. Associations of these clocks with age in this dataset may therefore be inflated.

### Analysis of EWAS data from Generation Scotland

We downloaded publicly available EWAS summary statistics for cross-sectional associations between baseline whole-blood CpG methylation and 14 prevalent disease states, and for longitudinal associations between baseline CpG methylation and 19 incident disease states from Generation Scotland cohort with N = 18,413 Scottish participants. DNA methylation was measured at 752,722 CpG sites using the Illumina MethylationEPIC array. Prevalent disease status was assessed primarily using self-reported information collected at baseline, whereas incident disease cases were identified through linkage to electronic health records during follow-up. We excluded disease outcomes with fewer than 100 cases, resulting in 10 prevalent and 14 incident diseases for downstream analyses. We used the EWAS summary statistics where age, sex, experimental batch, and cell-type compositions were adjusted. Logistic regressions were used for prevalent disease and Cox proportional hazard models were used for incident outcomes. DNAm was quantified on M-value scale. We then applied SARDINE to estimate associations between these disease outcomes and 29 epigenetic clocks, excluding clocks that had been trained using Generation Scotland data to minimize potential bias due to sample overlap. For SARDINE, we used reference DNAm covariance matrices residualized for age, age^2^, sex, and 12 estimated cell-type compositions.

## Data and code availability

EWAS summary statistics and DNAm covariance matrices derived from the TruDiagnostic cohort are available at https://qlu-lab.org/data.html. SARDINE software is freely available at https://github.com/qlu-lab/SARDINE. TruDiagnostic data is available upon request to TruDiagnostic Inc.

## Author contributions

Y.Wu and Q.L. conceived and designed the study.

Y.Wu and Y.Wang performed data analysis.

Y.Wu developed the method.

R.M. implemented the software.

V.B.D. and K.B.S. processed TruDiagnostic data.

R.S. advised on TruDiagnostic study design and result interpretation.

S.M.M. and L.L.S. advised on epigenetic clock models and results interpretation.

Q.L. advised on statistical and genetic issues.

Y.Wu, Y.Wang, and Q.L. wrote the manuscript.

All authors contributed to manuscript editing and approved the manuscript.

## Acknowledgments

We thank Shan Leng and the members of the UW-Madison social genomics group for helpful discussion and feedback.

