## Supplementary Note for "SARDINE unlocks EWAS summary statistics for large-scale epigenetic biomarker analysis"

Yuchang Wu

Department of Biostatistics and Medical Informatics
University of Wisconsin–Madison


Nov. 2026

### 1 Statistical Framework

#### 1.1 EWAS model and notation: $M = Y\beta^\top + \epsilon$

We consider the following data-generating model, which is widely used in epigenome-wide association studies (EWAS):

$$M = Y\beta^\top + \epsilon \tag{1}$$

where

- $M = (M_1 \quad M_2 \quad \cdots \quad M_J)$ is an $N \times J$ matrix containing DNA methylation measurements on the beta-value scale for $N$ individuals across $J$ CpG sites
- $Y$ is an $N \times 1$ column vector containing each individual's exposure
- $\beta = (\beta_1 \quad \beta_2 \quad \cdots \quad \beta_J)^\top$ is a $J \times 1$ column vector containing CpG effect sizes
- $\epsilon = (\epsilon_1 \quad \epsilon_2 \quad \cdots \quad \epsilon_J)$ is an $N \times J$ matrix containing random noise, which are uncorrelated across individuals

For simplicity, we first illustrate the setting in which the EWAS is designed to study the effect of an exposure, such as smoking, on DNAm. Generalizing this framework to EWAS in which $Y$ is the dependent variable is straightforward (see section 3.2).

Without loss of generality, we *assume* that $Y$ and DNAm $M_j$ at any CpG site $j$ are centered at 0. For individual $i$ and CpG $j$, the model can be written as

$$m_{ij} = y_i\beta_j + \epsilon_{ij} \tag{2}$$
$$\text{Var}(\epsilon_{ij}) = \sigma^2_{\epsilon j} \tag{3}$$

EWAS model written for the $j$-th CpG:

$$M_j = Y\beta_j + \epsilon_j$$

The ordinary least-squares (OLS) estimate is

$$\begin{aligned}\hat{\beta}_j &= (Y^\top Y)^{-1}Y^\top M_j \qquad (4)\\ &= \frac{\frac{1}{N}\sum y_i m_{ij}}{\frac{1}{N}\sum y_i^2} \\ &\xrightarrow{p} \frac{\text{Cov}(Y, M_j)}{\text{Var}(Y)} \qquad \text{as } N \to \infty\end{aligned}$$

Applying the variance operator rule $\text{Var}(AX) = A\text{Var}(X)A^\top$, the sampling variance is:

$$\begin{aligned}
\text{Var}(\hat{\beta}_j \mid Y) &= (Y^\top Y)^{-1} Y^\top \text{Var}(\boldsymbol{\epsilon}_j) Y (Y^\top Y)^{-1} \\
&= (Y^\top Y)^{-1} Y^\top (\sigma_{\epsilon j}^2 I_N) Y (Y^\top Y)^{-1} \\
&= \sigma_{\epsilon j}^2 (Y^\top Y)^{-1} \\
&= \frac{\sigma_{\epsilon j}^2}{N\left(\frac{1}{N}\sum y_i^2\right)} \\
&\approx \frac{\sigma_{\epsilon j}^2}{N\text{Var}(Y)} \qquad (5)
\end{aligned}$$

Define the $J \times J$ covariance matrix of DNAm as (recall that we assumed $M$ is centered at 0)

$$\Sigma := \frac{1}{N} M^\top M \qquad (6)$$

Ideally, this matrix should be estimated using the EWAS samples. However, in practice, it may be necessary to use an external reference dataset to approximate this matrix. We will assess the robustness of our framework to this reference approximation in section 2.1.

#### 1.2 Epigenetic clock as a weighted DNAm score

A linear methylation clock $C$ is defined as

$$C = Mw \tag{7}$$

Explicitly,

$$\begin{pmatrix} c_1 \\ c_2 \\ \vdots \\ c_N \end{pmatrix}_{N\times 1} = \begin{pmatrix} m_{11} & m_{12} & \cdots & m_{1J} \\ m_{21} & m_{22} & \cdots & m_{2J} \\ \vdots & \vdots & \ddots & \vdots \\ m_{N1} & m_{N2} & \cdots & m_{NJ} \end{pmatrix}_{N\times J} \begin{pmatrix} w_1 \\ w_2 \\ \vdots \\ w_J \end{pmatrix}_{J\times 1}$$

where $w = (w_1 \;\; w_2 \;\; \cdots \;\; w_J)^\top$ is a $J \times 1$ vector containing clock weights for each CpG. Typically, a clock uses only a small subset of CpGs, and we assign $w_j = 0$ to CpGs that are not used in the clock. Only CpGs with nonzero clock weights contribute to subsequent calculations. Without loss of generality,

1. we ignore the intercept term
2. we assumed $M$ is centered at 0 and thus $C$ is also centered at zero

Neither the intercept nor centering affects slopes, variances, or covariances. The in-sample variance of $C$ among these $N$ samples is

$$\begin{aligned} \mathrm{Var}(C) &= \frac{1}{N} C^\top C \\ &= \frac{1}{N} w^\top M^\top M w \\ &= w^\top \Sigma w \end{aligned} \tag{8}$$

Sections 2.2 and 2.3 cover the cases where $C$ is constructed in more complicated ways.

##### 1.3 Regressing the clock on the phenotype

Consider the individual-level regression

$$C = Y\lambda + u \tag{9}$$

where the variance of the noise term is

$$\mathrm{Var}(u) = \sigma_u^2 \tag{10}$$

The ordinary least-squares (OLS) estimate is given by

$$\begin{aligned}
\hat{\lambda} &= (Y^\top Y)^{-1} Y^\top C \\
&= (Y^\top Y)^{-1} Y^\top M w \\
&= (Y^\top Y)^{-1} Y^\top (M_1 \; M_2 \; \cdots \; M_J) w \\
&= (Y^\top Y)^{-1} Y^\top Y \hat{\beta}^\top w \\
&= \hat{\beta}^\top w = w^\top \hat{\beta} = \sum_{j=1}^{J} \hat{\beta}_j w_j
\end{aligned} \tag{11}$$

Note, the point estimate does not depend on DNAm variance-covariance matrix $\Sigma$. It also does not depend explicitly on $\mathrm{Var}(Y)$ because the scale of $Y$ is already encoded in $\beta_j$. The sampling variance is

$$\begin{aligned}
\mathrm{Var}(\hat{\lambda} \mid Y) &= (Y^\top Y)^{-1} \mathrm{Var}\,(u \mid Y)\, Y (Y^\top Y)^{-1} \\
&= \sigma_u^2 (Y^\top Y)^{-1} \approx \frac{\sigma_u^2}{N \mathrm{Var}(Y)}
\end{aligned} \tag{12}$$

where the residual variance is estimated by

$$\widehat{\sigma}_u^2 = \mathrm{Var}(u) = \mathrm{Var}(C) - \widehat{\lambda}^2 \mathrm{Var}(Y) = w^\top \Sigma w - \widehat{\lambda}^2 \mathrm{Var}(Y) \tag{13}$$

In section 1.5, we'll show how to estimate $\mathrm{Var}(Y)$ using only EWAS summary statistics. We then can compute $Z_\lambda = \frac{\hat{\lambda}}{\mathrm{SE}(\hat{\lambda})}$ and the corresponding two-sided $P$-value for testing $H_0 : \lambda = 0$.

$$Z_\lambda = \frac{\hat{\lambda}}{\mathrm{SE}(\hat{\lambda})} = \frac{w^\top \hat{\beta}}{\sqrt{\frac{w^\top \Sigma w}{N\,\mathrm{Var}(Y)} - \frac{\hat{\lambda}^2}{N}}} \tag{14}$$

The important insight is that the effect of an exposure on an epigenetic clock can be estimated without individual-level methylation data. The point estimate $\hat{\lambda} = w^\top \beta$ is the inner product of the pre-trained clock weights and the EWAS effect-size estimates across the CpGs included in the clock, and it does not depend on the methylation covariance matrix $\Sigma$.

##### 1.4 Regressing the phenotype on the clock

Consider the individual-level regression

$$Y = C\gamma + e \tag{15}$$

where

$$\text{Var}(e) = \sigma_e^2 \tag{16}$$

The OLS estimate is

$$\begin{aligned}
\hat{\gamma} &= (C^\top C)^{-1} C^\top Y \\
&= (w^\top M^\top M w)^{-1} w^\top M^\top Y \\
&= (w^\top M^\top M w)^{-1} w^\top \begin{pmatrix} M_1^\top \\ M_2^\top \\ \vdots \\ M_J^\top \end{pmatrix} Y \\
&= (w^\top M^\top M w)^{-1} w^\top \begin{pmatrix} Y^\top M_1 \\ Y^\top M_2 \\ \vdots \\ Y^\top M_J \end{pmatrix} \\
&= (w^\top M^\top M w)^{-1} w^\top (Y^\top Y) \begin{pmatrix} \hat{\beta}_1 \\ \hat{\beta}_2 \\ \vdots \\ \hat{\beta}_J \end{pmatrix} \\
&= \frac{1}{N w^\top \Sigma w} w^\top (Y^\top Y \hat{\beta}) \\
&= \frac{w^\top \hat{\beta}}{w^\top \Sigma w} \text{Var}(Y)
\end{aligned} \tag{17}$$

Thus, the summary-based method can recover the regression coefficient that would otherwise be estimated using individual-level data. The sampling variance

$$\begin{aligned}
\text{Var}(\hat{\gamma} \mid C) &= (C^\top C)^{-1} \sigma_e^2 C (C^\top C)^{-1} \\
&= \sigma_e^2 (C^\top C)^{-1} \\
&= \frac{1}{N} \frac{1}{w^\top \Sigma w} \sigma_e^2
\end{aligned} \tag{18}$$

We can estimate $\sigma_e^2$ as

$$\begin{aligned}\widehat{\sigma}_e^2 &= \mathrm{Var}(Y) - \hat{\gamma}^2 \mathrm{Var}(C) \\ &= \mathrm{Var}(Y) - \hat{\gamma}^2 w^\top \Sigma w \\ &= \mathrm{Var}(Y)(1 - R^2) \end{aligned} \tag{19}$$

where

$$\begin{aligned} R^2 &= \frac{\hat{\gamma}^2 w^\top \Sigma w}{\mathrm{Var}(Y)} \\ &= \left[\frac{w^\top \hat{\beta}}{w^\top \Sigma w} \mathrm{Var}(Y)\right]^2 \frac{w^\top \Sigma w}{\mathrm{Var}(Y)} \\ &= \frac{(w^\top \hat{\beta})^2}{w^\top \Sigma w} \mathrm{Var}(Y) = \frac{\hat{\lambda}^2}{w^\top \Sigma w} \mathrm{Var}(Y) & (20) \\ &= \hat{\lambda}\hat{\gamma} & (21) \end{aligned}$$

is the phenotypic variance captured by the clock.

$$Z_\gamma = \frac{\widehat{\gamma}}{\mathrm{SE}(\widehat{\gamma})} = \frac{w^\top \widehat{\beta}}{\sqrt{\frac{w^\top \Sigma w}{N\,\mathrm{Var}(Y)} - \frac{\widehat{\lambda}^2}{N}}} = \frac{\widehat{\lambda}}{\mathrm{SE}\left(\widehat{\lambda}\right)} = Z_\lambda \tag{22}$$

which is, as expected, identical to the $Z$ score for $\lambda$ in eq. (14).

One technical point: from eq. (13) in section 1.3, a non-negative value for $\sigma_u^2$ requires

$$\begin{aligned} w^\top \Sigma w - \hat{\lambda}^2\, \mathrm{Var}(Y) &\geq 0 \\ w^\top \Sigma w - (w^\top \hat{\beta})^2\, \mathrm{Var}(Y) &\geq 0 \\ \mathrm{Var}(Y) &\leq \frac{w^\top \Sigma w}{(w^\top \hat{\beta})^2} \end{aligned} \tag{23}$$

Denote

$$\mathrm{Var}(Y)_{\max} = \frac{w^\top \Sigma w}{(w^\top \hat{\beta})^2} = \frac{w^\top \Sigma w}{\hat{\lambda}^2} \tag{24}$$

and then we find that

$$\frac{\mathrm{Var}(Y)}{\mathrm{Var}(Y)_{\max}} = \frac{(w^\top \hat{\beta})^2}{w^\top \Sigma w} \mathrm{Var}(Y) = R^2 \tag{25}$$

Therefore, there's an upper bound for the value of $\mathrm{Var}(Y)$ (equivalently, $R^2$ should not be larger than 1).

##### 1.5 Estimating $\mathrm{Var}(Y)$ from summary statistics alone

If the variance of the phenotype used in the EWAS is unknown, it can be estimated using the EWAS summary statistics and a DNAm reference covariance matrix. From the EWAS model $M_j = Y\beta_j + \epsilon_j$, we have

$$\mathrm{Var}(M_j) = \hat{\beta}_j^2 \mathrm{Var}(Y) + \sigma_{\epsilon_j}^2 \tag{26}$$

$$\mathrm{Var}(\hat{\beta}_j) = \frac{\sigma_{\epsilon_j}^2}{N\mathrm{Var}(Y)} \tag{27}$$

Therefore,

$$\begin{aligned}\mathrm{Var}(M_j) &= \hat{\beta}_j^2 \mathrm{Var}(Y) + N\mathrm{Var}(\hat{\beta}_j)\mathrm{Var}(Y) \\ &= \left[\hat{\beta}_j^2 + N\mathrm{Var}(\hat{\beta}_j)\right] \mathrm{Var}(Y)\end{aligned}$$

Thus,

$$\mathrm{Var}(Y) = \frac{\mathrm{Var}(M_j)}{\hat{\beta}_j^2 + N\mathrm{Var}(\hat{\beta}_j)} = \frac{\Sigma_{jj}}{\hat{\beta}_j^2 + N\mathrm{Var}(\hat{\beta}_j)} \tag{28}$$

In practice, to improve numerical stability, we can use the median across CpGs:

$$\widehat{\mathrm{Var}(Y)} = \mathrm{Median}_j \left( \frac{\Sigma_{jj}}{\hat{\beta}_j^2 + N\mathrm{Var}(\hat{\beta}_j)} \right) \tag{29}$$

Here, $\Sigma_{jj}$ is the $j$-th diagonal element of the DNAm covariance matrix computed using the EWAS samples, and all other quantities are available in the EWAS summary statistics. Later in section 2.1, we will discuss the use of DNAm matrix from an external reference panel.

In section 1.4, we found that there's an upper bound for the value of $\mathrm{Var}(Y)$ (eq. (23)). When the clock is highly predictive of the phenotype (e.g., Horvath clock on chronological age), $\widehat{\mathrm{Var}(Y)}$ will be very close to $\mathrm{Var}(Y)_{\mathrm{max}}$. Due to numerical rounding inaccuracies, the estimated $\mathrm{Var}(Y)$ can become larger than its upper bound. In this case, we'll cap the $\widehat{\mathrm{Var}(Y)}$ to be slightly smaller than its upper bound.

##### 1.6 Covariate adjustments

In EWAS, it is common to adjust for covariates such as sex, age, cell type compositions, and other potential confounders. In this section we discuss the inclusion of covariates in linear regression EWAS. Later in section 3.7 we'll discuss the inclusion of covariates in logistic or Cox regressions.

The EWAS model with covariates can be written as

$$M_j = Y\beta_j + X\theta^\top + \epsilon_j \tag{30}$$

where $Y$ is the main exposure, $\beta_j$ is the effect size, and $X$ is an $N \times K$ covariate matrix. Define the projection, or residual-maker, matrix that regresses out the covariates:

$$P_{N\times N} = I - X(X^\top X)^{-1}X^\top \tag{31}$$

where $X$ contains a column of ones such that $P\mathbf{1} = 0$. Let

$$\widetilde{M_j} = PM_j \quad \text{(residualized DNAm)} \tag{32}$$

$$\widetilde{Y} = PY \quad \text{(residualized phenotype)} \tag{33}$$

According to the Frisch-Waugh-Lovell theorem [1, 2, 3],

$$\beta_j = \frac{\text{Cov}(\widetilde{Y}, \widetilde{M_j})}{\text{Var}(\widetilde{Y})} \tag{34}$$

The covariate-adjusted clock is

$$\widetilde{C} = \widetilde{M}w \tag{35}$$

The Frisch-Waugh-Lovell theorem allows the model with covariates to be rewritten as a model without covariates after both the exposure and outcome are residualized by the covariates. Using $\widetilde{C}$ and $\widetilde{Y}$, everything we derived previously go through. Just to be more specific, here we'll briefly show the derivations for $\lambda$ and $\gamma$ when there are covariates (in individual-level model).

**Covariate-adjusted Goal 1:** $C \sim Y$. Consider the individual-level clock-phenotype regression with covariates

$$C = Y\lambda + X\theta^\top + u \tag{36}$$

By the Frisch-Waugh-Lovell theorem, this is equivalent to

$$\widetilde{C} = \widetilde{Y}\lambda + \tilde{u} \tag{37}$$

Then

$$\begin{aligned}\hat{\lambda} &= \frac{\text{Cov}(\tilde{Y}, \tilde{C})}{\text{Var}(\tilde{Y})} = \frac{\text{Cov}(\tilde{Y}, \tilde{M}w)}{\text{Var}(\tilde{Y})} \\ &= w^\top \frac{\text{Cov}(\tilde{Y}, \tilde{M})}{\text{Var}(\tilde{Y})} \\ &= w^\top \hat{\beta} = \hat{\beta}^\top w = \sum_{j=1}^{J} \hat{\beta}_j w_j \end{aligned} \tag{38}$$

which has a same form with the case without any covariates (eq. (11)). Note here the $\hat{\beta}$ comes from the EWAS that adjusted for the same set of covariates. The sampling variance is

$$\text{Var}(\hat{\lambda}) = (\widetilde{Y}^\top \widetilde{Y})^{-1}\text{Var}(\tilde{u}) = \frac{1}{N\text{Var}(\widetilde{Y})}\sigma_u^2 \tag{39}$$

The residual variance is

$$\begin{aligned}\sigma_u^2 &= \text{Var}(\tilde{u}) \\ &\approx \text{Var}(\widetilde{C}) - \hat{\lambda}^2\text{Var}(\widetilde{Y}) \\ &= \frac{\widetilde{C}^\top \widetilde{C}}{N} - \hat{\lambda}^2\text{Var}(\widetilde{Y}) \\ &= w^\top \Sigma_{\text{adj}} w - \hat{\lambda}^2\text{Var}(\widetilde{Y}) \end{aligned} \tag{40}$$

Here the DNAm $\Sigma_{\text{adj}}$ was computed using residulized DNAm (after adjusting for the same set of covariates). Thus $\text{Var}(\hat{\lambda})$ also has the same form with the case without any covariates (eq. (12)).

**Covariate-adjusted Goal 2:** $Y \sim C$. Consider the individual-level clock-phenotype regression with covariates

$$Y = C\gamma + X\theta^\top + e \tag{41}$$

By the Frisch-Waugh-Lovell theorem,

$$\widetilde{Y} = \widetilde{C}\gamma + \tilde{e} \tag{42}$$

Therefore,

$$\hat{\gamma} = \frac{\mathrm{Cov}(\widetilde{Y}, \widetilde{C})}{\mathrm{Var}(\widetilde{C})} \tag{43}$$
$$= \frac{w^\top \mathrm{Cov}(\widetilde{Y}, \widetilde{M})}{w^\top \mathrm{Var}(\widetilde{M}) w}$$
$$= \frac{w^\top \beta}{w^\top \Sigma_{\mathrm{adj}} w} \mathrm{Var}(\widetilde{Y}) \tag{44}$$

which has the same form with the case without any covariates (eq. (17)). Its variance is

$$\mathrm{Var}(\hat{\gamma}) = (\widetilde{C}^\top \widetilde{C})^{-1} \mathrm{Var}(\tilde{e})$$
$$= \frac{1}{N} \frac{1}{w^\top \Sigma_{\mathrm{adj}} w} \sigma_e^2 \tag{45}$$

where residual variance is

$$\mathrm{Var}(\tilde{e}) = \sigma_e^2 = \mathrm{Var}(\widetilde{Y}) - \mathrm{Var}(\widetilde{C}) \hat{\gamma}$$
$$\approx \mathrm{Var}(\widetilde{Y}) - \gamma^2 \frac{1}{N} \widetilde{C}^\top \widetilde{C}$$
$$= \mathrm{Var}(\widetilde{Y}) - \gamma^2 w^\top \Sigma_{\mathrm{adj}} w \tag{46}$$

Thus $\mathrm{Var}\,\hat{\gamma}$ also has a same form with the case without any covariates (eq. (18)).

#### 2 Method inputs

In addition to EWAS summary statistics, SARDINE also requires a DNAm variance–covariance matrix $\Sigma$ and an epigenetic clock expressed as a weighted sum of CpG methylation levels, $C = Mw$. The derivations in the preceding section assume that $\Sigma$ is computed from the EWAS samples and that the published clock takes exactly this form.

Neither assumption holds in all applications. Individual-level DNAm data from the EWAS samples are often unavailable, so $\Sigma$ must be estimated from an external reference panel. In addition, several widely used clocks are not published in the form $C = Mw$: GrimAge includes chronological age and sex as predictors; the PC clocks are defined on principal components of CpG methylation; and DunedinPACE applies quantile normalization before computing the weighted sum.

This section examines the consequences of each departure and, where possible, how to correct for it.

#### 2.1 Reference DNAm variance-covariance matrix

Ideally, SARDINE needs a DNAm variance-covariance matrix derived from the EWAS samples, which is often not available in practice. Alternatively we can use a matrix from an external reference panel. In this section, we'll discuss the impact of the external DNAm variance-covariance matrix. Let superscript $^E$ denote the quantity estimated using EWAS samples, and superscript $^R$ denote the counterparts estimated using the reference panel.

**$\widehat{\boldsymbol{\lambda}}$ does not depend on $\boldsymbol{\Sigma}$ at all.** Check eqs. (11) and (38) for $\widehat{\lambda}$: they don't depend on $\Sigma$.

**Var($\boldsymbol{Y}$) uses only the diagonal terms in $\boldsymbol{\Sigma}$.** Recall in eq. (29), to estimate $\mathrm{Var}(Y)$, we need $\mathrm{Var}(M_j)$ which can be found the diagonal terms in $\Sigma^E$ derived from the EWAS samples. Using an external reference:

$$
\begin{aligned}
\widehat{\mathrm{Var}(Y)}^R &= \underset{j}{\mathrm{median}}\left(\frac{\Sigma^R_{jj}}{\hat{\beta}_j^2 + N\mathrm{Var}(\hat{\beta}_j)}\right) \\
&= \widehat{\mathrm{Var}(Y)}^E \underset{j}{\mathrm{median}}\left(\frac{\Sigma^R_{jj}}{\Sigma^E_{jj}}\right) \\
&= \kappa_1 \widehat{\mathrm{Var}(Y)}^E
\end{aligned} \tag{47}
$$

where

$$
\kappa_1 = \underset{j}{\mathrm{median}}\left(\frac{\Sigma^R_{jj}}{\Sigma^E_{jj}}\right) \tag{48}
$$

**$\widehat{\boldsymbol{\gamma}}$ uses the quadratic form.** Recall eqs. (17) and (44) for $\hat{\gamma}$; when we use external reference, it becomes

$$
\begin{aligned}
\widehat{\gamma}^R &= \frac{w^\top \hat{\beta}}{w^\top \Sigma^R w}\widehat{\mathrm{Var}(Y)}^R \\
&= \widehat{\gamma}^E \frac{w^\top \Sigma^E w}{w^\top \Sigma^R w}\underset{j}{\mathrm{median}}\left(\frac{\Sigma^R_{jj}}{\Sigma^E_{jj}}\right) \\
&= \tau\, \widehat{\gamma}^E
\end{aligned} \tag{49}
$$

where

$$
\kappa_2 = \frac{w^\top \Sigma^R w}{w^\top \Sigma^E w} \tag{50}
$$

$$
\tau = \frac{\kappa_1}{\kappa_2} \tag{51}
$$

So $\kappa_1$ tracks the ratio of diagonal/variance elements while $\kappa_2$ tracks the ratio of the quadratic forms (which are heavily dominated by the off-diagonal covariance structure, especially as the number of CpGs grows). And

$$R^{2,R} = \frac{\hat{\lambda}^2}{w^\top \Sigma^R w} \widehat{\mathrm{Var}(Y)}^R = \tau\, R^{2,E} \tag{52}$$

$$\begin{aligned} Z^R &= \frac{w^\top \hat{\beta}}{\sqrt{\frac{w^\top \Sigma^R w}{N \widehat{\mathrm{Var}(Y)}^R} - \frac{\hat{\lambda}^2}{N}}} \\ &= \frac{w^\top \hat{\beta}}{\sqrt{\frac{w^\top \Sigma^E w}{N \widehat{\mathrm{Var}(Y)}^E} - \frac{\hat{\lambda}^2}{N}}} \frac{\sqrt{\frac{w^\top \Sigma^E w}{N \widehat{\mathrm{Var}(Y)}^E} - \frac{\hat{\lambda}^2}{N}}}{\sqrt{\frac{w^\top \Sigma^R w}{N \widehat{\mathrm{Var}(Y)}^R} - \frac{\hat{\lambda}^2}{N}}} \\ &= Z^E \, \frac{\sqrt{\frac{w^\top \Sigma^E w}{N \widehat{\mathrm{Var}(Y)}^E} - \frac{\hat{\lambda}^2}{N}}}{\sqrt{\frac{w^\top \Sigma^R w}{N \widehat{\mathrm{Var}(Y)}^R} - \frac{\hat{\lambda}^2}{N}}} \end{aligned} \tag{53}$$

Another quantity that often appears in derivations

$$\begin{aligned} \frac{w^\top \Sigma^R w}{\widehat{\mathrm{Var}(Y)}^R} &= \frac{w^\top \Sigma^E w}{\widehat{\mathrm{Var}(Y)}^E} \frac{w^\top \Sigma^R w}{\widehat{\mathrm{Var}(Y)}^R} \frac{\widehat{\mathrm{Var}(Y)}^E}{w^\top \Sigma^E w} \\ &= \frac{1}{\tau} \, \frac{w^\top \Sigma^E w}{\widehat{\mathrm{Var}(Y)}^E} \end{aligned} \tag{54}$$

EWAS usually adjusts for several common confounding factors (e.g., sex, age, and cell type compositions) in epigenetic data which typically induce widespread positive cross-correlations among clock CpGs.

**Case 1: Under-adjusted External Reference** This case happens when the EWAS adjusted for major covariates (e.g., sex, age, and cell type compositions) while the external reference was computed from "raw" or less-adjusted data. The consequences are:

- $\kappa_1 > 1$, since the unadjusted confounders inflate the individual CpG variances
- $\kappa_2 \gg \kappa_1$, since the quadratic form $w^\top \Sigma w$ sums $J$ variance terms and $J(J-1)$ covariance terms. Because the off-diagonal terms in $\Sigma^R$ are

typically much larger and more positive than those in $\Sigma^E$, the quadratic term should have large inflation, leading to

- $\tau < 1$
- deflated $\hat{\gamma}$
- deflated $Z$ score and loss of statistical power

**Case 2: Over-adjusted External Reference** This case happens when the EWAS did not adjust for certain covariates (e.g., cell type compositions) while the external reference was computed from "over"-adjusted DNAm data (e.g., cell type adjusted). Because the covariates "explain" meaningful variances in DNAm, the consequences are

- $\kappa_1 < 1$ since the some of the variances in the external data was regressed out by the additional covariate adjustments
- $\kappa_2 \ll \kappa_1$ since regressing out the confounders/covariates reduces the correlation structure and thus decreases the quadratic form. Because the covariate adjustment impacts $J + J(J-1)$ terms in $\kappa_2$, it has a much larger impact on $\kappa_2$ than $\kappa_1$, leading to
- $\tau > 1$
- inflated $\hat{\gamma}$
- inflated $Z$ score and type-I error (false positives)

**Case 3: $\Sigma^R = c\Sigma^E$ for any $c > 0$** In this case, the two matrices are proportional to each other. Then we get $\tau = 1$ and all the point estimates, their variances and $P$-values remain the same.

**Practical Recommendations** Given the above results, we can have the following practical guidelines:

1. Users should use a external reference panel with matched covariate adjustment as used in the EWAS.
2. If a such reference panel cannot be found, it is better to use an under-adjusted external reference knowing that the $\hat{\gamma}$ is underestimated and statistical power is reduce, but we won't have inflated type-I error.

#### 2.2 GrimAge clocks

The GrimAge clocks [4, 5] were constructed using age, sex, and a surrogate of DNAm smoking pack years, and other surrogates of DNAm biomarkers. V1 used a total of 10 features whereas V2 used a total of 12 features. They used an identical set of 1,030 CpGs. Two stages (elastic net and Cox regression) were used to construct the clocks. In stage 1, surrogate $k$ is an elastic net predictor

$$\hat{S}_k = \alpha_{0k}\mathbf{1}_N + \alpha_{Ak}A + \alpha_{Sk}S + M\theta_k$$

In stage 2, an elastic net Cox model for time-to-death yields a linear predictor

$$x^\top\beta = \beta_A A + \beta_S S + \sum_{k=1}^{K} \beta_k \widehat{S}_k$$

which is then calibrated to units of years to get the final clock

$$\text{GrimAge} = t_0 + t_1 x^\top\beta$$

Therefore, the GrimAge clock can actually be written as

$$\text{GrimAge} = c_0 + aA + sS + Mw \tag{55}$$

where $c_0$ is intercept, $a$ is the coefficient for Age $A$, and $s$ is the coefficient for Sex $S$, $M$ is an $N \times J$ matrix of CpG values for $N$ individuals across $J$ CpGs (for GrimAge, $J = 1,030$), and $w$ is the CpG weights. The GrimAge is quantified on chronological age scale. The clock file for GrimAge contains intercept, age, female indicator, CpGs, and their corresponding weights.

In Sardine, the clock was assumed to be a simple $Mw$ without considering intercept, or sex and age or other features that may contribute to the clock construction. Here we'll show that when age and sex are included in covariates, the using of a simple $Mw$ for clock will not be problematic for the inferences of $\lambda$ and $\gamma$. When age cannot be adjusted (for example, when age is the phenotype), we'll show that a simple $Mw$ will lead to biased results and we can correct it using $a$ which is readily available. Denote the GrimAge clock solely based on CpGs

$$\text{GrimAge}^* = Mw \tag{56}$$

For simplicity, in this section we'll use $C$ to represent GrimAge clock and $C^*$ for GrimAge*. Recall in the previous section we defined residual maker $P_{N\times N} = I - X(X^\top X)^{-1}X^\top$ (eq. (31) in section 1.6).

**Both age and sex are in covariates.** Here the covariates matrix $X$ contains both age and sex. Then we have

$$\widetilde{C} = PC = aPA + sPS + PMw = PMw = PC^* = \widetilde{C}^* \tag{57}$$

Let $\hat{\beta}_j$ denote the CpG $j$'s effect on $Y$ in the covariate-adjusted EWAS, $\hat{\lambda}_{\text{full}}$ denote the coefficient of the "raw" clock $C$ on $Y$, and $\hat{\lambda}_{\text{SARDINE}}$ denote the estimate given by SARDINE using only $C^*$ and $C = c_0 + aA + sS + C^*$. Then,

$$\begin{aligned}\hat{\beta}_j &= \frac{\text{Cov}(\widetilde{M_j}, \widetilde{Y})}{\text{Var}(\widetilde{Y})} = \frac{\text{Cov}(PM_j, PY)}{\text{Var}(PY)} \\ &= \frac{M_j^\top P^\top PY}{Y^\top P^\top PY} = \frac{M_j^\top PY}{Y^\top PY}\end{aligned} \tag{58}$$

$$\hat{\lambda}_{\text{full}} = \frac{\text{Cov}(\widetilde{C}, \widetilde{Y})}{\text{Var}(\widetilde{Y})} = \frac{C^\top P^\top Y}{Y^\top P^\top Y} \tag{59}$$

$$\begin{aligned}\hat{\lambda}_{\text{SARDINE}} &= w^\top \hat{\beta} = \frac{w^\top M^\top PY}{Y^\top PY} \\ &= \frac{(PMw)^\top PY}{Y^\top PY} = \frac{(PC)^\top PY}{Y^\top PY} \\ &= \frac{C^\top PY}{Y^\top PY} = \hat{\lambda}_{\text{full}}\end{aligned} \tag{60}$$

And the clock variance

$$\begin{aligned}w^\top \Sigma_{\text{adj}} w &= \frac{w^\top M^\top PMw}{N} \\ &= \frac{(PMw)^\top (PMw)}{N} = \frac{(PC)^\top PC}{N} = \text{Var}(\widetilde{C})\end{aligned} \tag{61}$$

Therefore, when age and sex are adjusted for, $\hat{\lambda}, \hat{\gamma}$, their variances, Wald statistics $Z$, and $P-$values from SARDINE based on $Mw$ alone are numerically identical to those that would be obtained from the "raw" GrimAge clock given that $\Sigma_{\text{adj}}$ is computed using EWAS samples with sex and age adjusted.

For logistic and Cox regressions, we can consider a linear predictor through which the likelihood or partial likelihood is linked with data

$$\begin{aligned}\eta &= \vartheta_0 + \gamma C + \psi_A A + \psi_S S + Z\phi \\ &= \vartheta_0 + \gamma(c_0 + aA + sS + Mw) + \psi_A A + \psi_S S + Z\phi \\ &= \vartheta_0 + \gamma c_0 + \gamma Mw + (\gamma a + \psi_A)A + (\gamma s + \psi_S)S + Z\phi\end{aligned} \tag{62}$$

So clearly at the individual-level, the two designs (one included age and sex in clock vs. the other does not) index the same set of linear predictors and the inferences on $\gamma$ will be identical. Therefore, when age and sex are included in covariates, it does not matter whether they are included in the clock construction or not. The SARDINE approximation for logistic and Cox regression will be discussed in later Sections.

*Age Acceleration Clock.* The age acceleration version of a clock is defined as the residual from the linear regression of Clock $\sim$ Chronological age:

$$\begin{aligned}\text{AA} &= P_{\{1,A\}}C = P_{\{1,A\}}(c_0 + aA + sS + Mw) \\ &= sP_{\{1,A\}}S + P_{\{1,A\}}Mw \end{aligned} \tag{63}$$

When AgeAccelGrim is used in the regression *and* age and sex are included in the regression covariates, we have

$$P\text{AA} = P(sP_{\{1,A\}}S + P_{\{1,A\}}C^*) = PC^* = PC \tag{64}$$

Therefore, when age and sex are adjusted for, SARDINE estimates the association of AgeAccelGrim with the phenotype.

These conclusions also apply to sex-specific phenotypes such as breast cancer, ovarian cancer, prostate cancer ect where sex cannot be included in covariates (but age is included in covariates). Because for these phenotypes, all the EWAS samples are of a same sex, the sex contribution in GrimAge is essentially an intercept.

**Sex is in covariates, but age is not.** Now the covariates matrix only contains sex (and an intercept). The residualized full GrimAge clock becomes

$$\begin{aligned}\widetilde{C} &= P_{\{1,S\}}C = P_{\{1,S\}}(c_0 + aA + sS + Mw) \\ &= aP_{\{1,S\}}A + P_{\{1,S\}}Mw \\ &= aP_{\{1,S\}}A + P_{\{1,S\}}C^* \\ &= aP_{\{1,S\}}A + \widetilde{C}^* \end{aligned} \tag{65}$$

The estimate given by SARDINE solely based on $C^*$ is

$$\widehat{\lambda}_{\text{SARDINE}} = w^\top \hat{\beta} = \frac{(C^*)^\top P_{\{1,S\}}Y}{Y^\top P_{\{1,S\}}Y} \tag{66}$$

and the estimates given by the "raw" (or "full") GrimAge clock is

$$
\begin{aligned}
\widehat{\lambda}_{\text{full}} &= \frac{C^\top P_{\{1,S\}} Y}{Y^\top P_{\{1,S\}} Y} = \frac{\left(aP_{\{1,S\}}A + \widetilde{C}^*\right)^\top P_{\{1,S\}} Y}{Y^\top P_{\{1,S\}} Y} \\
&= a\frac{A^\top P_{\{1,S\}} Y}{Y^\top P_{\{1,S\}} Y} + \frac{(C^*)^\top P_{\{1,S\}} Y}{Y^\top P_{\{1,S\}} Y} \\
&= \widehat{\lambda}_{\textsc{Sardine}} + a\widehat{\delta}_{A\sim Y+S}
\end{aligned} \tag{67}
$$

where $\hat{\delta}_{A\sim Y+S}$ is the coefficient from linear regression of Age $\sim Y +$ Sex. To perform inferences, we also need the variance of residualized full clock

$$
\begin{aligned}
\operatorname{Var}(\widetilde{C}) &= \frac{C^\top P_{\{1,S\}} C}{N} \\
&= \frac{(aA + C^*)^\top P_{\{1,S\}}(aA + C^*)}{N} \\
&= a^2\frac{A^\top P_{\{1,S\}} A}{N} + 2a\frac{A^\top P_{\{1,S\}} C^*}{N} + \frac{(C^*)^\top P_{\{1,S\}} C^*}{N} \\
&= a^2 \operatorname{Var}(\widetilde{A}) + 2a \operatorname{Cov}(\widetilde{A}, \widetilde{C}^*) + \operatorname{Var}(\widetilde{C}^*) \\
&= a^2 \operatorname{Var}(\widetilde{A}) + 2a \operatorname{Cov}(\widetilde{A}, \widetilde{C}^*) + w^\top \Sigma_{\text{adj}} w
\end{aligned} \tag{68}
$$

where $\widetilde{A} = P_{\{1,S\}}A$.

$$
\begin{aligned}
\widehat{\gamma}_{\text{full}} &= \frac{C^\top P_{\{1,S\}} Y}{C^\top P_{\{1,S\}} C} = \frac{(aA + C^*)^\top P_{\{1,S\}} Y}{C^\top P_{\{1,S\}} C} \\
&= \frac{aA^\top P_{\{1,S\}} Y + (C^*)^\top P_{\{1,S\}} Y}{C^\top P_{\{1,S\}} C} \\
&= \frac{aN \operatorname{Cov}(\widetilde{A}, \widetilde{Y}) + N \operatorname{Cov}(\widetilde{C}^*, \widetilde{Y})}{N \operatorname{Var}(\widetilde{C})} \\
&= \frac{a \operatorname{Cov}(\widetilde{A}, \widetilde{Y}) + \hat{\gamma}_{\textsc{Sardine}} \operatorname{Var}(\widetilde{C}^*)}{\operatorname{Var}(\widetilde{C})} \\
&= \widehat{\gamma}_{\textsc{Sardine}} \frac{\operatorname{Var}(\widetilde{C}^*)}{\operatorname{Var}(\widetilde{C})} + a\frac{\operatorname{Cov}(\widetilde{A}, \widetilde{Y})}{\operatorname{Var}(\widetilde{C})} \\
&= \widehat{\gamma}_{\textsc{Sardine}} \frac{w^\top \Sigma_{\text{adj}} w}{\operatorname{Var}(\widetilde{C})} + a\frac{\operatorname{Cov}(\widetilde{A}, \widetilde{Y})}{\operatorname{Var}(\widetilde{C})}
\end{aligned} \tag{69}
$$

In general, we may not have enough information to recover $\widehat{\lambda}_{\text{full}}$ or $\widehat{\gamma}_{\text{full}}$ since we don't know $\hat{\delta}_{A\sim Y+S}$, $\operatorname{Var}(\widetilde{A})$, $\operatorname{Cov}(\widetilde{A}, \widetilde{C}^*)$, and $\operatorname{Cov}(\widetilde{A}, \widetilde{Y})$.

*Special case: Phenotype is Age.* When age is the phenotype $Y$, $\hat{\delta}_{A\sim Y+S} = 1$, and the formulas can be simplified:

$$\widehat{\lambda}_{\text{full}} = a + \widehat{\lambda}_{\text{SARDINE}} \tag{70}$$

$$\text{Var}(\hat{\lambda}_{\text{full}}) = \text{Var}(\hat{\lambda}_{\text{SARDINE}}) \tag{71}$$

$$Z = (a + \widehat{\lambda}_{\text{SARDINE}})/(\text{SE}(\widehat{\lambda}_{\text{SARDINE}})) \tag{72}$$

$$\begin{aligned} \text{Var}(\widetilde{C}) &= a^2\,\text{Var}(\widetilde{A}) + 2a\widehat{\lambda}_{\text{SARDINE}}\,\text{Var}(\widetilde{A}) + w^\top \Sigma_{\text{adj}} w \\ &= (a^2 + 2a\widehat{\lambda}_{\text{SARDINE}})\,\text{Var}(\widetilde{A}) + w^\top \Sigma_{\text{adj}} w \end{aligned} \tag{73}$$

Thus under this special case we can recover point estimate and $P$-value for $\lambda_{\text{full}}$ using SARDINE's output since the value of $a$ is publicly available.

One thing to note here, if age is standardized in EWAS, SARDINE's $\hat{\lambda}$ represent the effect size per unit standardized clock, thus, we should scale $a$ accordingly to also be in a per-1-SD unit for eqs. (70) and (72): $a_{\text{std}} = a \times \text{sd}_{\text{age}}$.

**Age is in covariates, but sex is not.** For a special phenotype such as "Is_female", sex cannot be included in covariates. And if age is adjusted, we'll have similar results as those in previous subsection:

$$\widehat{\lambda}_{\text{full}} = b + \widehat{\lambda}_{\text{SARDINE}} \tag{74}$$

$$Z = (b + \widehat{\lambda}_{\text{SARDINE}})/(\text{SE}(\widehat{\lambda}_{\text{SARDINE}})) \tag{75}$$

**Neither age nor sex is in covariates.** Here the design matrix $X = [\mathbf{1}, \boldsymbol{Z}]$ where $\boldsymbol{Z}$ represents other possible covariates (such as cell type compositions), but age and sex are not adjusted. Then following similar derivations, we have

$$\widetilde{C} = PC = a\widetilde{A} + s\widetilde{S} + \widetilde{C}^* \tag{76}$$

and

$$\begin{aligned} \widehat{\lambda}_{\text{full}} &= \frac{C^\top PY}{Y^\top PY} = \frac{(a\widetilde{A} + s\widetilde{S} + \widetilde{C}^*)^\top PY}{Y^\top PY} \\ &= \widehat{\lambda}_{\text{SARDINE}} + a\hat{\delta}_A + s\hat{\delta}_S \end{aligned} \tag{77}$$

where

$$\hat{\delta}_A = \frac{A^\top PY}{Y^\top PY} \tag{78}$$

$$\hat{\delta}_S = \frac{S^\top PY}{Y^\top PY} \tag{79}$$

are the coefficients of Y in the linear regression of $A \sim Y + Z$ and $S \sim Y + Z$. Check: when sex is adjusted, $\hat{\delta}_S = 0$ and when age is adjusted $\hat{\delta}_A = 0$, consistent with our previous results.

#### 2.3 PC clocks

The principal component (PC) clock [6] was developed to improve clock "reliability". It first conduct PCA analyses for 78,464 CpGs, and then used PCs with elastic net to train clock models. To compute the PC version of a clock, users only need to project their DNAm data into their PC space and then multiply by the elastic net (EN) coefficients. They shared data for computing six PC clocks that are measured in chronological age: `PCHorvath1, PCHorvath2, PCHannum, PCPhenoAge, PCDNAmTL, and PCGrimAgeV1`.

In this Section, we'll show that PC clocks can be effectively written as a weighted sum across the 78,464 CpGs plus an intercept, and thus can be directly used by SARDINE. Let's first briefly show how PC clocks were constructed. Let $X_t = X - m_c$ denote a centered DNAm $N \times p$ matrix (where $p = 78,464$) and $m_c$ is a matrix of DNAm mean values. Then conduct SVD

$$X_t = U\Sigma V^\top = AV^\top \tag{80}$$

where $A = U\Sigma$. And then running EN to choose PCs and estimate their coefficients

$$\min_{\beta_0,\beta} \frac{1}{N}\sum_{i=1}^{N} \frac{1}{2}(y_i - \beta_0 - A_i^\top \beta)^2 + \lambda_{\min}\left[\frac{(1-\alpha)||\beta||_2^2}{2} + \alpha||\beta||_1\right] \tag{81}$$

where $\alpha$ is set to be 0.5. Let $m_c \in \mathbb{R}^p$, rotation $V_{dc} \in \mathbb{R}^{p\times dc}$, EN model $\theta \in \mathbb{R}^{dc}$, and intercept $\theta_{0c}$. Here we use subscript $dc$ to denote the PCs that are selected by the EN model. $V_{dc}$ is the right singular vector matrix $V$ from the SVD of the training matrix with columns restricted to the $dc$ PCs with non-zero EN coefficients. Then the PC clock for a new DNAm data $M_{N\times p}$ is given by

$$\begin{aligned} L^{(c)} &= (M - m_c)V_{dc}\theta_c + \theta_{0c} \\ &= MV_{dc}\theta_c + \theta_{0c} - m_c V_{dc}\theta_c \\ &= Mw_c + a_c \end{aligned} \tag{82}$$

where $w_c = V_{dc}\theta_c$ are the CpG weights and $a_c$ is a constant intercept. For `PCHannum, PCPhenoAge, PCDNAmTL`, eq. (82) is the clock value; for `PCHorvath`, it's used as input for a piecewise inverse transformation function to get the final clock. For `PCGrimAge`, eq. (82) is used to first build eight surrogates, and then combine them to get the final clock value.

Since the PC clocks are linear functions of CpGs, the weight $w_c$ can be used by SARDINE to estimate the PCClock-phenotype associations. Since author released the values of $V_{dc}$ and $\theta_c$, we can easily compute $w_c$ for each PC clock.

`PCGrimAge`. As we discussed in section 2.2, computing `PCGrimAge` similarly also takes a little bit more algebra. It will first compute a PC version of the surrogates, then sum over the surrogates using weights from `GrimAge` clock.

$$S_k = (M - m_c)V_{dc}\theta_{ck} + \theta_k^F \text{Female} + \theta_k^A \text{Age} + \theta_{0ck} \tag{83}$$

$$\begin{aligned} \texttt{PCGrimAge} &= \gamma_0 + \sum_{k \in S} \gamma_k S_k + \gamma_F \text{Female} + \gamma_A \text{Age} \\ &= c_0 + M w_c + c_F \text{Female} + c_A \text{Age} \end{aligned} \tag{84}$$

where

$$w_c = V_{dc} \sum_{k \in S} \gamma_k \theta_{ck} \tag{85}$$

$$c_F = \gamma_F + \sum_{k \in S} \gamma_k \theta_k^F \tag{86}$$

$$c_A = \gamma_A + \sum_{k \in S} \gamma_k \theta_k^A \tag{87}$$

$$c_0 = \gamma_0 + \sum_{k \in S} \gamma_k \theta_{0ck} - m_c w_c \tag{88}$$

Note, the PC clock paper only released data for computing the PC version of `GrimAgeV1`.

#### 2.4 DunedinPACE clock

The `DunedinPACE` clock [7] is another epigenetic clock calculated as a weighted sum across CpGs. However, unlike its predecessor `DunedinPoAm` [8], the `DunedinPACE` algorithm first applies quantile normalization to align the input data with the DNA methylation (DNAm) distribution of the clock's training data for 20,000 "gold-standard" background CpGs (including the 173 clock CpGs) to remove the global distributional differences. The weighted sum is then computed using this transformed DNAm, ensuring the output clock values are on the same scale as those in the original Dunedin Study cohort.

During this transformation, an individual's CpG values are first sorted and replaced with the correspondingly ranked DNAm values from the training cohort. These substituted values are then restored to the individual's original CpG order. Once this process is complete for all individuals, the algorithm calculates the weighted sum across the transformed CpGs to yield the final `DunedinPACE` value. Consequently, every sample achieves the exact same cumulative distribution as the "gold standard" reference, while the relative ranking of CpGs within each individual is preserved.

Let $g_{(1)} \leq g_{(2)} \leq \cdots \leq g_{(20{,}000)}$ be the sorted "gold-standard" mean beta values from the clock training cohort, and $m_{ij}$ the $i$-th individual's DNAm beta value at the $j$-th CpG. The quantile normalization replaces it by the gold-standard value at the same rank

$$Q_i(m_{ij}) = g_{(\mathrm{rank}(m_{ij}))} = g_{ij} \tag{89}$$

where $\mathrm{rank}(m_{ij})$ is the rank of the $j$-th CpG for individual $i$, we use notation $g_{ij}$ for convenience. Then

$$\mathrm{PACE}_i^{\mathrm{official}} = c_0 + \sum_{j=1}^{173} w_j Q_i(m_{ij}) \tag{90}$$

$$\mathrm{PACE}_i^{\mathrm{simple}} = \sum_{j=1}^{173} w_j m_{ij} \tag{91}$$

where $c_0 = -1.949859$ is the intercept, $\mathrm{PACE}_i^{\mathrm{simple}}$ is the one that SARDINE utilizes. All between-sample variation in $\mathrm{PACE}_i^{\mathrm{official}}$ comes from the permutation since every sample has a same set of DNAm values and they only differ in their orders.

$$\begin{aligned}\Delta_i &= \text{PACE}_i^{\text{official}} - \text{PACE}_i^{\text{simple}} \\ &= \sum_{j=1}^{173} w_j \left[Q_i(m_{ij}) - m_{ij}\right] + c_0 \qquad (92) \\ \text{PACE}_i^{\text{official}} &= \text{PACE}_i^{\text{simple}} + \Delta_i \qquad (93)\end{aligned}$$

Apparently, this $\Delta_i$ may not be a linear function of $\text{PACE}_i^{\text{simple}}$ and the official clock is not an exact linear function of the simple weighted sum unless under some special conditions.

**Special cases**:

For concreteness, let's assume the observed/measured betas contain a technical location-scale artifact of its quantile-normalized value:

$$m_{ij} = a_i g_{ij} + b_i \qquad (94)$$

Note, since DNAm cannot be negative, $a_i$ will be positive. Then we have

$$\begin{aligned}\text{PACE}_i^{\text{official}} &= c_0 + \sum_{j=1}^{173} w_j g_{ij} \qquad (95) \\ \text{PACE}_i^{\text{simple}} &= \sum_{j=1}^{173} w_j m_{ij} = a_i \sum_{j=1}^{173} w_j g_{ij} + b_i \sum_{j=1}^{173} w_j \\ &= a_i \left(\text{PACE}_i^{\text{official}} - c_0\right) + b_i \sum_{j=1}^{173} w_j \\ &= a_i \text{PACE}_i^{\text{official}} - a_i c_0 + b_i \sum_{j=1}^{173} w_j \qquad (96)\end{aligned}$$

The covariance between the two clocks:

$$\begin{aligned}\text{Cov}(\text{PACE}^{\text{official}}, \text{PACE}^{\text{simple}}) &= \text{Cov}(\text{PACE}_i^{\text{official}}, \text{PACE}_i^{\text{simple}}) \\ &= \text{Cov}(\text{PACE}_i^{\text{official}}, a_i \text{PACE}_i^{\text{official}}) \\ &+ \text{Cov}(\text{PACE}_i^{\text{official}}, b_i \sum_{j=1}^{173} w_j - a_i c_0) \qquad (97)\end{aligned}$$

Covariance of the clock with phenotype $Y$:

$$\operatorname{Cov}(Y, \text{PACE}^{\text{simple}}) = \operatorname{Cov}(Y_i, a_i \text{PACE}_i^{\text{official}}) + \operatorname{Cov}\left(Y_i, b_i \sum_{j=1}^{173} w_j - a_i c_0\right) \tag{98}$$

1. If the artifact only creates a location shift and no scaling (i.e., $a_i = 1, b_i \neq 0$), then we have

$$\text{PACE}_i^{\text{simple}} = \text{PACE}_i^{\text{official}} - c_0 + b_i \sum_{j=1}^{173} w_j \tag{99}$$

   Since the clock intercept $c_0 = -1.949859$ and $\sum_j w_j = 3.5884 \neq 0$ for `DunedinPACE`, each sample's simple clock value is the official one plus an offset and this offset is sample-specific. If further the offset $b_i$ is same across individuals, then these two clocks will be perfectly correlated.

2. Phenotype association testing. We want to know whether the association test of phenotype $\sim$ PACE$^{\text{official}}$ is equivalent with phenotype $\sim$ PACE$^{\text{simple}}$.

   - If $(a_i, b_i) \perp (Y_i, \sum_{j=1}^{173} w_j g_{ij})$, and denote $\rho$ as the correlation between (PACE$^{\text{official}}$ and PACE$^{\text{simple}}$) then

$$\operatorname{Cov}(Y, \text{PACE}^{\text{simple}}) = \mathbb{E}(a_i) \operatorname{Cov}\left(Y, \text{PACE}^{\text{official}}\right) \tag{100}$$

$$\operatorname{Cov}(\text{PACE}^{\text{official}}, \text{PACE}^{\text{simple}}) = \mathbb{E}(a_i) \operatorname{Var}\left(\text{PACE}^{\text{official}}\right) \tag{101}$$

$$\operatorname{Cor}(Y, \text{PACE}^{\text{simple}}) = \rho \operatorname{Cor}\left(Y, \text{PACE}^{\text{official}}\right) \tag{102}$$

$$Z^{\text{simple}} = \rho Z^{\text{official}} \tag{103}$$

$$\lambda^{\text{simple}} = \mathbb{E}(a_i) \lambda^{\text{official}} \tag{104}$$

$$\gamma^{\text{simple}} = \frac{\rho^2}{\mathbb{E}(a_i)} \gamma^{\text{official}} \tag{105}$$

   Note, $\rho \leq 1$ with equality only when $a_i = a$ and $b_i = b$ are constant across individuals. Effect size from a linear regression of $Y$ on clock is attenuated and with a larger $P$-value for the simple clock. The point estimates are rescaled rather than attenuated in general. The type-I error remains valid and only power is lost.

- When the artifacts are correlated with phenotype (e.g., most cases are from batch 1 while most controls are from batch 2), the simple clock will yield a false association. In contrast, the official clock forces all $a_i \to 1$ and $b_i \to 0$ relative to the gold standard, eliminating this technical confounding. However, it also worth exploring whether the official score removes genuine biology signals, for example, when a phenotype shifts the global methylation distribution.

  If the batch effects are included in the regression covariates, the effects of $b_i$ should be removed, however, the impact of the scaling factor $a_i$ remains leading to biased results.

Thus, to conclude, in general, this quantile normalization is a non-linear transformation, SARDINE cannot recover the effect sizes for the `DunedinPACE` clock based on the transformed DNAm. Instead, SARDINE's results for `DunedinPACE` should be interpreted as effect sizes for the clock based on the "raw" DNAm data.

### 3 Extensions to other EWAS designs

The framework derived in the preceding sections assumes that the EWAS is a linear regression of DNAm beta values on the phenotype, $M = Y\beta^\top + \epsilon$. It estimates the slopes of the linear regressions of the clock on the phenotype ($\lambda$) and of the phenotype on the clock ($\gamma$).

In practice, an EWAS can depart from this "preferred" design in four ways: the methylation scale, the direction of the regression, the model for the phenotype, or the handling of covariates. To accommodate these designs, the reported summary statistics are first converted to their equivalents under the "preferred" model, and the estimators of the preceding sections are then applied.

The subsections below derive these conversions in turn for EWAS on the M-value scale, EWAS with the regression direction reversed ($Y = M\alpha + e$), logistic EWAS of binary phenotypes, and Cox proportional hazards EWAS of time-to-event phenotypes. For binary and time-to-event phenotypes, we can also recover the odds ratio or hazard ratio that an individual-level logistic or Cox regression of the phenotype on the clock would estimate. We then extend these results to two-stage, residual-based EWAS designs that researchers sometimes would use.

#### 3.1 EWAS on the M-value scale

It's not uncommon for researcher to use DNAm M-value in a linear regression. We can convert regression coefficients from the M-value scale to the Beta-value scale by utilizing Taylor expansion.

Let $M_j^*$ denote M-value while $M_j$ denote Beta-value at the $j$-th CpG site.

$$M_j^* = \log_2\left(\frac{M_j}{1 - M_j}\right) \tag{106}$$

Define

$$\begin{aligned} f(x) &= \log_2\left(\frac{x}{1-x}\right) = \log_2 x - \log_2(1-x) \\ &= \frac{1}{\ln(2)}\left[\ln(x) - \ln(1-x)\right] \\ f'(x) &= \frac{1}{\ln(2)}\frac{1}{x(1-x)} \end{aligned}$$

where $\ln(\cdot)$ is the natural logarithm (i.e., base of $e$). Then eq. (106) can be approximated by Taylor expansion around the mean

$$M_j^* \approx f(\mu_j) + f'(\mu_j)(M_j - \mu_j) \tag{107}$$

where $\mu_j = \mathbb{E}(M_j)$ is the $j$-th CpG site's mean methylation Beta-value, which can be approximated by an external reference panel. Note the linear relationship between the DNAm Beta-value and M-value: later we'll find that this is the key that we can convert the regression coefficients for many regression models. Then, we can have

$$\begin{aligned} \mathrm{Cov}(M_j^*,\ Y) &\approx \mathrm{Cov}\left(f(\mu_j) + f'(\mu_j)(M_j - \mu),\ Y\right) \\ &= f'(\mu_j)\mathrm{Cov}(M_j,\ Y) \\ &= \frac{1}{\ln(2)}\frac{1}{\mu_j(1-\mu_j)}\mathrm{Cov}(M_j,\ Y) \end{aligned} \tag{108}$$

$$\mathrm{Var}(M_j^*) \approx \left[\frac{1}{\ln(2)}\frac{1}{\mu_j(1-\mu_j)}\right]^2 \mathrm{Var}(M_j) \tag{109}$$

Thus, using the scaling factor $\ln(2)\mu_j(1-\mu_j)$, we can convert EWAS summary statistics from M-value scale to the Beta-value scale.

##### 3.2 Reversed EWAS direction: $Y = M\alpha + e$

If the EWAS model is

$$Y = \alpha_j M_j + e_j \tag{110}$$

could we still use its summary statistics to estimate $\lambda$ and $\gamma$? The answer is yes because we can convert this summary statistics to the ones produced by $M_j \sim Y$ regressions.

$$\begin{aligned}
\alpha_j &= (M_j^\top M_j)^{-1} M_j^\top Y \qquad (111)\\
&= \frac{\mathrm{Cov}(Y, M_j)}{\mathrm{Var}(M_j)}\\
&= \frac{\mathrm{Cov}(Y, M_j)}{\mathrm{Var}(Y)} \frac{\mathrm{Var}(Y)}{\mathrm{Var}(M_j)}\\
&= \frac{\mathrm{Var}(Y)}{\mathrm{Var}(M_j)} \beta_j\\
\mathrm{Var}(\widehat{\alpha}_j) &= \frac{\mathrm{Var}(e_j)}{N\,\mathrm{Var}(M_j)} \qquad (112)\\
&\approx \frac{\mathrm{Var}(Y) - \widehat{\alpha}_j^2\,\mathrm{Var}(M_j)}{N\,\mathrm{Var}(M_j)}\\
&= \frac{1}{N} \frac{\mathrm{Var}(Y)}{\mathrm{Var}(M_j)} - \frac{1}{N} \widehat{\alpha}_j^2 \qquad (113)\\
\frac{\mathrm{Var}(Y)}{\mathrm{Var}(M_j)} &= \widehat{\alpha}_j^2 + N\,\mathrm{Var}(\widehat{\alpha}_j) \qquad (114)
\end{aligned}$$

And now we can convert $\widehat{\alpha}_j$ to $\widehat{\beta}_j$:

$$\widehat{\beta}_j \approx \frac{\widehat{\alpha}_j}{\widehat{\alpha}_j^2 + N\,\mathrm{Var}(\widehat{\alpha}_j)} \tag{115}$$

$$\mathrm{SE}(\widehat{\beta}_j) \approx \frac{\mathrm{SE}(\widehat{\alpha}_j)}{\widehat{\alpha}_j^2 + N\,\mathrm{Var}(\widehat{\alpha}_j)} \tag{116}$$

##### 3.3 Binary phenotypes: Logistic EWAS

When $Y$ is binary (for example, disease status), the EWAS model can be a logistic regression of $Y$ on each $M_j$. The question is how can we use the summary statistics from a logistic regression to estimate the associations between phenotype $Y$ and clock $C$. Without loss of generality, we assume $M_j$ is centered at 0 in this subsection.

$$\operatorname{logit}(\mathbb{P}(Y = 1 \mid M)) = \alpha_0 + M_j \alpha_j \tag{117}$$

Since our main method was developed based on linear regression models, the task is whether we can link $\alpha_j$ with $\beta_j$ in eq. (4) estimated from the linear regression

$$M_j = Y\beta_j + \epsilon_j \tag{118}$$

Since $Y$ is binary, it's easy to see

$$\beta_j = \mathbb{E}(M_j \mid Y = 1) - \mathbb{E}(M_j \mid Y = 0) \tag{119}$$

Let's list here the moments that will be used later:

$$\begin{aligned}
\mathbb{E}(Y \mid M_j) &= p_j && (120)\\
\mathbb{E}(Y) &= K && (121)\\
\mathbb{E}(M_j) &= \mathbb{E}[\mathbb{E}(M_j \mid Y)] \\
&= K\mathbb{E}(M_j \mid Y = 1) + (1-K)\mathbb{E}(M_j \mid Y = 0) \\
&= K\mu_{1j} + (1-K)\mu_{0j} && (122)\\
\mu_1 &= \mathbb{E}(M \mid Y = 1) && (123)\\
\mu_0 &= \mathbb{E}(M \mid Y = 0) && (124)\\
\operatorname{Var}(Y \mid M_j) &= p_j(1-p_j) && (125)\\
\operatorname{Var}(M_j) &= \beta_j^2 \operatorname{Var}(Y) + \sigma^2_{\epsilon_j} && (126)\\
\operatorname{Var}(M_j \mid Y) &= \operatorname{Var}(Y\beta_j + \epsilon_j \mid Y) = \sigma^2_{\epsilon_j} && (127)\\
\mathbb{E}(YM_j) &= p_j\mathbb{E}(M_j \mid Y = 1) = p_j\mu_1 && (128)\\
\operatorname{Cov}(Y, M_j) &= \mathbb{E}(YM_j) - \mathbb{E}(Y)\mathbb{E}(M_j) \\
&= K\mu_{1j} - K(K\mu_{1j} + (1-K)\mu_{0j}) \\
&= K(1-K)(\mu_{1j} - \mu_{0j}) && (129)
\end{aligned}$$

We can double-check that $\beta = \operatorname{Cov}(Y, M)/\operatorname{Var}(Y) = \mu_1 - \mu_0$ is consistent. We next calculate the log-odds ratio from the logistic regression model eq.(117) (we omitted subscript $j$):

$$
\begin{aligned}
\operatorname{logit}(\mathbb{P}(Y=1 \mid M)) &= \ln\left[\frac{\mathbb{P}(Y=1 \mid M)}{\mathbb{P}(Y=0 \mid M)}\right] \\
&= \ln\left[\frac{\mathbb{P}(Y=1, M)}{\mathbb{P}(M)} \frac{\mathbb{P}(M)}{\mathbb{P}(Y=0, M)}\right] \\
&= \ln\left[\frac{\mathbb{P}(M \mid Y=1)\mathbb{P}(Y=1)}{\mathbb{P}(M \mid Y=0)\mathbb{P}(Y=0)}\right] \\
&= \ln\left[\frac{\mathbb{P}(M \mid Y=1)}{\mathbb{P}(M \mid Y=0)}\right] + \ln\left(\frac{K}{1-K}\right)
\end{aligned} \tag{130}
$$

Now we need to know the conditional density function of $M$. If we are willing to *assume* normality with equal variance:

$$M \mid Y=0 \sim N(\mu_0, \sigma_\epsilon^2) \tag{131}$$

$$M \mid Y=1 \sim N(\mu_1, \sigma_\epsilon^2) \tag{132}$$

Then we have

$$
\begin{aligned}
\operatorname{logit}(\mathbb{P}(Y=1 \mid M)) &= -\frac{(M-\mu_1)^2}{2\sigma_\epsilon^2} + \frac{(M-\mu_0)^2}{2\sigma_\epsilon^2} + \ln\left(\frac{K}{1-K}\right) \\
&= \frac{\mu_1-\mu_0}{\sigma_\epsilon^2} M + \frac{\mu_0^2-\mu_1^2}{2\sigma_\epsilon^2} + \ln\left(\frac{K}{1-K}\right) \\
&= \alpha_0 + M\alpha
\end{aligned} \tag{133}
$$

Now we can clearly see that

$$\beta_j = \alpha_j \sigma_{\epsilon_j}^2 \tag{134}$$

Thus now the task is to estimate the DNA methylation residual variance $\sigma_\epsilon^2$ within each group. Denote $\operatorname{Var}(M_j) = \sigma_{M_j}^2 = \Sigma_{jj}$

$$
\begin{aligned}
\Sigma_{jj} &= \beta_j^2 \operatorname{Var}(Y) + \sigma_{\epsilon_j}^2 \\
&= \sigma_{\epsilon_j}^4 \alpha_j^2 \operatorname{Var}(Y) + \sigma_{\epsilon_j}^2
\end{aligned} \tag{135}
$$

We can solve this quadratic equation to get $\sigma_\epsilon^2$

$$\sigma_{\epsilon_j}^2 = \frac{-1 + \sqrt{1 + 4\alpha_j^2 \Sigma_{jj} \operatorname{Var}(Y)}}{2\alpha_j^2 \operatorname{Var}(Y)} \tag{136}$$

and

$$\beta_j = \alpha_j \sigma^2_{\epsilon_j} = \frac{-1 + \sqrt{1 + 4\alpha_j^2 \Sigma_{jj} \operatorname{Var}(Y)}}{2\alpha_j \operatorname{Var}(Y)}$$
$$= \frac{2\alpha_j \Sigma_{jj}}{1 + \sqrt{1 + 4\alpha_j^2 \Sigma_{jj} \operatorname{Var}(Y)}} \tag{137}$$

When $\alpha_j$ is small, we can utilize the approximation that $\sqrt{1+x} = 1 + x/2 + O(x^2)$ to get

$$\beta_j \approx \frac{\alpha_j \Sigma_{jj}}{1 + \alpha_j^2 \Sigma_{jj} \operatorname{Var}(Y) + O(\alpha_j^4)}$$
$$\approx \alpha_j \Sigma_{jj} - \alpha_j^3 \Sigma_{jj}^2 \operatorname{Var}(Y) + O(\alpha_j^5)$$
$$\approx \alpha_j \Sigma_{jj} \tag{138}$$

Note converting $\alpha_j$ to $\beta_j$ (eq. (137)) requires the value of $\operatorname{Var}(Y)$, which is not directly provided in the EWAS summary statistics. Quick check: when $\alpha_j = 0$, $\beta_j = 0$, so same hypothesis testing. In a previous section we estimated $\operatorname{Var}(Y)$, however that one requires the input EWAS model to be a linear regression of DNA methylation Beta-value $M \sim Y$. That is, converting $\alpha_j$ to $\beta_j$ requires knowing $\operatorname{Var}(Y)$, but estimating $\operatorname{Var}(Y)$ requires knowing $\beta_j$. This creates a chicken-and-egg paradox!

However, we can use $\operatorname{Var}(\hat{\alpha_j})$ reported in the (logistic regression) EWAS summary statistics to estimate $\operatorname{Var}(Y)$. Let's define $\theta = (\alpha_{0j}, \alpha_j)^\top$, $x_i = (1, M_{ij})^\top$, we can get Fisher information matrix

$$\boldsymbol{I}(\alpha_j) = -\mathbb{E}\left[\frac{\partial^2 \ell}{\partial\theta\partial\theta^\top}\right]$$
$$= \mathbb{E}\left[\sum_{i=1}^N w_i x_i x_i^\top\right]$$
$$= \mathbb{E}\left[\begin{pmatrix} \sum_i w_i & \sum_i w_i M_{ij} \\ \sum_i w_i M_{ij} & \sum_i w_i M_{ij}^2 \end{pmatrix}\right]$$
$$\approx N \begin{pmatrix} \mathbb{E}(w) & \mathbb{E}(wM) \\ \mathbb{E}(wM) & \mathbb{E}(wM^2) \end{pmatrix} \tag{139}$$

and

$$
\begin{aligned}
\operatorname{Var}(\hat{\alpha}_j) &= [\boldsymbol{I(\alpha_j)}^{-1}]_{22} && (140)\\
&= \frac{1}{N(\mathbb{E}(w_j M_j^2) - \mathbb{E}(w_j M_j)^2/\mathbb{E}(w_j))} \\
&= \frac{1}{N\mathbb{E}(w_j)(\mathbb{E}(w_j M_j^2)/\mathbb{E}(w_j) - \mathbb{E}(w_j M_j/\mathbb{E}(w_j))^2)} \\
&= \frac{1}{N\mathbb{E}(w_j)\operatorname{Var}_Q(M_j)} && (141)
\end{aligned}
$$

where we denote

$$
\operatorname{Var}_Q(M_j) = \mathbb{E}(w_j M_j^2)/\mathbb{E}(w_j) - [\mathbb{E}(w_j M_j)/\mathbb{E}(w_j)]^2 \qquad (142)
$$

Recall for any random variable $X$, $\operatorname{Var}(X) = \mathbb{E}(X^2) - (\mathbb{E}(X))^2$, here the $\operatorname{Var}_Q(M_j)$ is called "weighted" (or "tilted") variance. $N$ is the total sample size. Now we need to calculate $\mathbb{E}(w_j) = \mathbb{E}(p_j(1-p_j))$ and $\operatorname{Var}_Q(M_j)$.

Since

$$
\begin{aligned}
\operatorname{Var}(Y) &= \mathbb{E}[\operatorname{Var}(Y \mid M_j)]) + \operatorname{Var}(\mathbb{E}(Y \mid M_j)) && (143)\\
&= \mathbb{E}(w_j) + \operatorname{Var}(p_j) && (144)
\end{aligned}
$$

where

$$
\begin{aligned}
p_j(M_j) = \mathbb{E}(Y \mid M_j) &= \frac{1}{1+e^{-(\alpha_0+\alpha_j M_j)}} && (145)\\
&\approx p_j(\mathbb{E}(M_j)) + p'(\mathbb{E}(M_j))(M_j - \mathbb{E}(M_j)) \quad \text{Taylor expansion} && (146)\\
&= p_j(\mathbb{E}(M_j)) + \alpha_j p_j(\mathbb{E}(M_j))[1 - p_j(\mathbb{E}(M_j))](M_j - \mathbb{E}(M_j)) && (147)
\end{aligned}
$$

The coefficient before $M_j$ can be estimated by $\operatorname{Cov}(p_j, M_j)/\operatorname{Var}(M_j)$, thus,

$$
\begin{aligned}
\operatorname{Var}(p_j) &\approx \left[\frac{\operatorname{Cov}(p_j, M_j)}{\operatorname{Var}(M_j)}\right]^2 \operatorname{Var}(M_j) && (148)\\
&= \frac{\operatorname{Cov}(p_j, M_j)^2}{\operatorname{Var}(M_j)} && (149)\\
&= \frac{\operatorname{Cov}(Y, M_j)^2}{\operatorname{Var}(M_j)} && (150)\\
&= \frac{\beta_j^2(\operatorname{Var}(Y))^2}{\Sigma_{jj}} && (151)
\end{aligned}
$$

where we used tower rule to get $\text{Cov}(Y, M_j) = \text{Cov}(p_j, M_j)$. And we have

$$\mathbb{E}(w_j) = \text{Var}(Y) - \text{Var}(p_j) \tag{152}$$

$$= \text{Var}(Y)\left[1 - \frac{\beta_j^2 \,\text{Var}(Y)}{\Sigma_{jj}}\right] \tag{153}$$

To compute $\text{Var}_Q(M_j)$, we need to utilize **Stein's Lemma**, which essentially states that for a normally distributed variable $M_j$ with mean 0 and variance $\Sigma_{jj}$, and a differentiable function $f(M_j)$,

$$\mathbb{E}(M_j f(M_j)) = \Sigma_{jj}\mathbb{E}[f'(M_j)] \tag{154}$$

After extensively applying Stein's Lemma (details omitted to save space), we can have

$$\mathbb{E}(w_j M_j) = \Sigma_{jj}\mathbb{E}(w_j'') \tag{155}$$

$$\mathbb{E}(w_j M_j^2) = \Sigma_{jj}\mathbb{E}(w_j) + \Sigma_{jj}^2\mathbb{E}(w_j'') \tag{156}$$

where $w_j' = d(p_j - p_j^2)/dM_j$ is the derivative of $w_j$ with respect to $M_j$. Then,

$$\begin{aligned}
\text{Var}_Q(M_j) &= \Sigma_{jj} + \Sigma_{jj}^2\left[\frac{\mathbb{E}(w_j'')}{\mathbb{E}(w_j)} - \left(\frac{\mathbb{E}(w_j')}{\mathbb{E}(w_j)}\right)^2\right] \\
&= \Sigma_{jj} + \Sigma_{jj}^2\left[\frac{d^2}{dM_j^2}\log(w_j)\right] \\
&\approx \Sigma_{jj} - 2\alpha_j^2\Sigma_{jj}^2\mathbb{E}(w_j) \\
&= \Sigma_{jj}\left[1 - 2\alpha_j^2\Sigma_{jj}\mathbb{E}(w_j)\right]
\end{aligned} \tag{157}$$

Since the formulas will become very messy, we'll use some approximations to make the derivations clearer: $\beta_j \approx \alpha_j\Sigma_{jj}$, $\mathbb{E}(w_j) \approx \text{Var}(Y)$, and let's denote $R_j^2 = \beta_j^2\,\text{Var}(Y)/\Sigma_{jj}$, which quantifies the proportion of DNA methylation variance at site $j$ that's "explained" by phenotype $Y$. Then we have

$$\mathrm{Var}_Q(M_j) \approx \Sigma_{jj}(1 - 2R_j^2) \tag{158}$$

$$\mathbb{E}(w_j) \approx \mathrm{Var}(Y)(1 - R_j^2) \tag{159}$$

$$\mathrm{Var}(\hat{\alpha}_j) \approx \frac{1}{N\Sigma_{jj}\,\mathrm{Var}(Y)(1 - 3R_j^2 + 2R_j^4)} \tag{160}$$

$$\begin{aligned}
\mathrm{Var}(Y) &\approx \frac{1}{N\Sigma_{jj}\,\mathrm{Var}(\hat{\alpha}_j)(1 - 3R_j^2)} \\
&\approx \frac{1}{N\Sigma_{jj}\,\mathrm{Var}(\hat{\alpha}_j)}(1 + 3R_j^2) \\
&\approx \frac{1}{N\Sigma_{jj}\,\mathrm{Var}(\hat{\alpha}_j)}
\end{aligned} \tag{161}$$

Recall that $R_j^2$ is a function of $\mathrm{Var}(Y)$ but under small effect (i.e., small $\beta_j$ in EWAS which should be reasonable at an individual CpG level), $R_j^2$ is relatively small and can be omitted. To improve numerical stability, we can use median (and can also use an external reference for $\Sigma_{jj}$)

$$\widehat{\mathrm{Var}(Y)} \approx \mathrm{median}_j \frac{1}{N\Sigma_{jj}\,\mathrm{Var}(\hat{\alpha}_j)} \tag{162}$$

Plugging into eq. (137), we can convert $\widehat{\alpha}_j$ to $\widehat{\beta}_j$. Now we also need to use delta method to compute $\mathrm{Var}(\widehat{\beta}_j)$:

$$\frac{d\beta_j}{d\alpha_j} = \frac{2\Sigma_{jj}}{\sqrt{1 + 4\alpha_j^2\Sigma_{jj}\,\mathrm{Var}(Y)}\left[1 + \sqrt{1 + 4\alpha_j^2\Sigma_{jj}\,\mathrm{Var}(Y)}\right]} \tag{163}$$

$$= \frac{2\Sigma_{jj}}{A(1 + A)} \tag{164}$$

$$\mathrm{Var}(\widehat{\beta}_j) = \left[g'(\alpha_j)\right]^2 \mathrm{Var}(\widehat{\alpha}_j) \tag{165}$$

$$= \frac{4\Sigma_{jj}^2}{A^2(1 + A)^2}\,\mathrm{Var}(\widehat{\alpha}_j) \tag{166}$$

where we denote $A = \sqrt{1 + 4\alpha_j^2\Sigma_{jj}\,\mathrm{Var}(Y)}$. Finally, we can convert an EWAS summary statistics from a logistic model to a linear model:

$$\widehat{\beta}_j = \frac{2\Sigma_{jj}}{1 + A}\widehat{\alpha}_j \tag{167}$$

$$\mathrm{SE}(\widehat{\beta}_j) = \frac{2\Sigma_{jj}}{A(1 + A)}\mathrm{SE}(\widehat{\alpha}_j) \tag{168}$$

Note here we'll still use the logistic EWAS model's P value.

##### 3.4 Time-to-event phenotypes: Cox proportional hazards EWAS

Our method can take the summary statistics from a Cox regression as input. The reason is that we can approximate the estimates from the Cox regression by a linear regression coefficient. Here we show the detailed derivations.

We can write the Cox Proportional Hazards (PH) model as

$$h(t|X_i) = h_0(t)\exp(X_i\beta_{\text{Cox}}) \tag{169}$$

where $h_0(t)$ is an unspecified baseline hazard (non-negative) function, and $h(t|X_i = 0) = h_0(t)$. The coefficients $\beta_{\text{Cox}}$ are assumed to be independent of time $t$. Let

- $i = 1, 2, \ldots, N$ individuals
- $X_i$: a vector of $J$ predictors (including intercept). In EWAS context, it represents DNA methylation in Beta-value
- $y_{i0} \in \{0, 1\}$: case-control status at baseline (”prevalence”)

For those $y_{i0} = 0$ samples (i.e., ”controls” at baseline when $t = 0$):

$$\delta_i = \begin{cases} 0 & \text{if censored} \\ 1 & \text{if an event was observed} \end{cases}$$
$$T_i : \text{time from baseline to incident disease OR censoring}$$
$$C_i : \text{end-of-study time (administrative censoring)}$$

Define the binary end-of-study incident disease status:

$$D_i = \delta_i = I(T_i \leq C_i) \tag{170}$$

Note, here $D_i$ is the number of disease cases among those who were controls at the beginning of the study (that is, the number of new cases during study time period.)

The cumulative hazard function up to time $t$ is:

$$
\begin{aligned}
H(t|X_i) &= \int_0^t h(u|X_i)du \qquad (171)\\
&= \int_0^t h_0(u)\exp(X_i\beta_{\text{Cox}})du \\
&= \exp(X_i\beta_{\text{Cox}})\int_0^t h_0(u)du \\
&= H_0(t)\exp(X_i\beta_{\text{Cox}}) \qquad (172)
\end{aligned}
$$

where $H_0(t) := \int_0^t h_0(u)du$ is defined as the baseline cumulative hazard. Survival probability (i.e., prob. of still being a control) by time $t$ is:

$$
S(t|X_i) = \mathbb{P}(T_i > t|X_i) \qquad (173)
$$

Since the hazard function is defined as

$$
h(t) = \lim_{\Delta t \to 0} \frac{\mathbb{P}(t \leq T \leq t + \Delta t \mid T \geq t)}{\Delta t} \qquad (174)
$$

which is the instantaneous rate of failure at time $t$, given survival up to $t$:

$$
\begin{aligned}
\mathbb{P}(t \leq T \leq t + \Delta t \mid T \geq t) &= \frac{\mathbb{P}(t \leq T \leq t + \Delta t)}{\mathbb{P}(T \geq t)} \\
&= \frac{\mathbb{P}(t \leq T \leq t + \Delta t)}{S(t)} \\
&\approx \frac{f(t)\Delta t}{S(t)} \qquad (175)
\end{aligned}
$$

where $f(t)$ is the PDF of $T$. Thus,

$$
h(t) \approx \lim_{\Delta t \to 0} \frac{f(t)}{S(t)} = \frac{f(t)}{S(t)} \qquad (176)
$$

Now let's find the relationship between $S(t)$ & $H(t)$.

$$S(t) = P(T > t) = 1 - F(t) \tag{177}$$

$$\frac{dS(t)}{dt} = S'(t) = -f(t) \tag{178}$$

$$h(t) = \frac{f(t)}{S(t)} = -\frac{S'(t)}{S(t)} = -\frac{d}{dt}(\ln S(t)) \tag{179}$$

$$\begin{aligned} H(t) &= \int_0^t h(u)du = -\int_0^t \frac{d}{du}[\ln S(u)]du \\ &= \ln S(0) - \ln S(t) \\ &= -\ln S(t) \end{aligned} \tag{180}$$

Recall every subject is alive at time 0 (we're conditioning on a subset of $y_{i0} = 0$), $S(0) = \mathbb{P}(T > 0) = 1 \Rightarrow \ln S(0) = 0$.

$$S(t) = \exp(-H(t)) \tag{181}$$

$$\begin{aligned} &= \exp\left(-H_0(t)\exp(X\beta_{\text{Cox}})\right) \\ &= e^{-H_0(t)e^{X\beta_{\text{Cox}}}} \\ &= S_0(t)^{\exp(X\beta_{\text{Cox}})} \end{aligned} \tag{182}$$

where $S_0(t) = e^{-H_0(t)}$.

Now let's consider a ***linear*** regression of case-control status (defined as $D$) on $X$ at the ***end*** of study:

$$D = \alpha X + e \tag{183}$$

$$\alpha = \frac{\text{Cov}(X, D)}{\text{Var}(X)} = \frac{\mathbb{E}(XD) - \mathbb{E}(X)\mathbb{E}(D)}{\text{Var}(X)} \tag{184}$$

Under the Cox model, the probability of being a case at the end of study is

$$\mathbb{P}(D_i = 1|X_i) = p(X_i) \tag{185}$$

$$\begin{aligned} &= \mathbb{P}(T_i \le C_i|X_i) \\ &= 1 - S(C_i) \\ &= 1 - \exp(-H(C_i)) \\ &= 1 - \exp\left[-H_0(C_i)\exp(X_i\beta_{\text{Cox}})\right] \end{aligned} \tag{186}$$

Rearrange to take natural log (will use these later):

$$\log(1 - p(X_i)) = -H_0(C_i)\exp(X_i\beta_{\text{Cox}}) \tag{187}$$

$$\log\left[-\log(1 - p(X_i))\right] = \log(H_0(C_i)) + X_i\beta_{\text{Cox}} \tag{188}$$

Since $D_i$ is binary,

$$
\begin{aligned}
\mathbb{E}(D_i|X_i) &= \mathbb{P}(D_i = 1|X_i) = p(X_i) && (189)\\
\mathbb{E}(XD) &= \mathbb{E}\left[\mathbb{E}(XD|X)\right] = \mathbb{E}\left[X\mathbb{E}(D|X)\right] \\
&= \mathbb{E}(Xp(X)) && (190)\\
\alpha &= \frac{\mathrm{Cov}(X,D)}{\mathrm{Var}(X)} = \frac{\mathbb{E}(Xp(X)) - \mathbb{E}(X)\mathbb{E}(p(X))}{\mathrm{Var}(X)} \\
&= \frac{\mathrm{Cov}(p(X),X)}{\mathrm{Var}(X)} && (191)
\end{aligned}
$$

**Taylor Expansion Approximation.** Note, we don't know the distribution of $X$, but we can use Taylor expansion to approximate:

$$
\begin{aligned}
p(X) &\approx p(\mu_x) + p'(\mu_x)(X - \mu_x) && (192)\\
\alpha &\approx \frac{\mathrm{Cov}(X, p(\mu_x) + p'(\mu_x)(X - \mu_x))}{\mathrm{Var}(X)} \\
&= p'(\mu_x) && (193)
\end{aligned}
$$

Taking the derivative of eq. (186):

$$
\begin{aligned}
p'(X) &= -\exp\left[-H_0(C)\exp(X\beta_{\mathrm{Cox}})\right](-H_0(C))\exp(X\beta_{\mathrm{Cox}})\beta_{\mathrm{Cox}} \\
&= \beta_{\mathrm{Cox}} H_0(C)\exp\left[-H_0(C)\exp(X\beta_{\mathrm{Cox}})\right]\exp(X\beta_{\mathrm{Cox}}) \\
&= \beta_{\mathrm{Cox}}\left[-\ln(1 - p(X))\right](1 - p(X)) && (194)
\end{aligned}
$$

Let

$$
\begin{aligned}
\mu_x &= \mathbb{E}X && (195)\\
p(\mu_x) &= \pi = \frac{E}{N} && (196)
\end{aligned}
$$

where $p(\mu_x) = \pi$ is the final proportion of cases, $E$ is the number of incident cases, and $N$ is the number of controls at the beginning ($t = 0$) of the study. Then we have

$$
\alpha \approx (1 - \pi)[-\ln(1 - \pi)]\beta_{\mathrm{Cox}} \quad (197)
$$

When the disease is rare (i.e., $\pi$ is small), $-\ln(1 - \pi) \approx \pi$, and

$$
\alpha \approx \pi(1 - \pi)\beta_{\mathrm{Cox}}
$$

Therefore, we can convert EWAS summary statistics from a Cox PH model to the linear regression (of $Y \sim M$) scale by

$$\widehat{\alpha}_j \approx (1-\pi)[-\ln(1-\pi)]\beta_{\text{Cox},j} \tag{198}$$

$$\text{SE}(\widehat{\alpha}_j) \approx (1-\pi)[-\ln(1-\pi)]\text{SE}\left(\widehat{\beta}_{\text{Cox},j}\right) \tag{199}$$

where $\pi$ or both $E$ and $N$ are usually reported by the study. Then we can continue converting these into a linear regression of $M \sim Y$ scale as the final input.

##### DNAm in M-value

If the M-value ($M_j^*$) DNAm is used in the Cox PH model (eq. 169), we can first convert the effect size from the M-value scale to Beta-value scale by taking advantage of the approximately linear relationship between M- and Beta-value (eq.(107)). Let's use $\beta^*_{\text{Cox},j}$ and $\beta_{\text{Cox},j}$ to denote the effect sizes under M- and Beta-value, respectively. Then the Cox PH model using M-value can be written as

$$\begin{aligned} h(t; M_j^*) &= h_0^*(t)\text{exp}(M_j^*\beta^*_{\text{Cox},j}) \qquad (200) \\ &= h_0^*(t)\text{exp}\left\{[f(\mu_j) + f'(\mu_j)(M_j - \mu_j)]\beta^*_{\text{Cox},j}\right\} \\ &= h_0^*(t)\text{exp}((f(\mu_j) - \mu_j f'(\mu_j))\beta_j^*)\ \text{exp}(M_j f'(\mu_j)\beta_j^*) \\ &= \tilde{h}_0^*(t)\text{exp}\left[M_j f'(\mu_j)\beta_j^*\right] \qquad (201) \end{aligned}$$

Comparing with the Cox PH model under Beta-value (eq.(169)), we can easily find that

$$\hat{\beta}_{\text{Cox},j} = \frac{1}{\ln(2)}\frac{1}{\mu_j(1-\mu_j)}\hat{\beta}^*_{\text{Cox},j} \tag{202}$$

$$\text{SE}(\hat{\beta}_{\text{Cox},j}) = \frac{1}{\ln(2)}\frac{1}{\mu_j(1-\mu_j)}\text{SE}(\hat{\beta}^*_{\text{Cox},j}) \tag{203}$$

where $\mu_j = \mathbb{E}(M_j)$ is the mean DNA methylation Beta-value. Also note that when there are covariates, we still need the raw mean Beta-value (not the "residualized" since its mean is always 0).

##### 3.5 Recovering individual-level odds ratio

When the phenotype $Y \in \{0, 1\}$ is binary, we can run a logistic regression to estimate the effects of clock on the log of odds

$$\text{logit}(P(Y = 1)) = \gamma_0 + \gamma_{\text{logistic}} C \tag{204}$$

Since our main framework was derived based on linear regressions, by default we can estimate $\gamma_{\text{linear}}$ from a linear regression of $Y \sim C$. So the questions is can we convert $\gamma_{\text{linear}}$ to $\gamma_{\text{logistic}}$?

The answer is yes; and we can use two different approaches: Taylor expansion is more straightforward but can have large bias, and another approach utilizes Stein's lemma with a little more complicated derivations but much less bias.

**Taylor expansion to link between $\gamma_{\text{lin}}$ and $\gamma_{\text{logistic}}$**

$$\mathbb{P}(Y = 1 \mid C) = \frac{1}{1 + e^{-(\gamma_0 + \gamma_{\text{logistic}} C)}} = p(C) = f(C) \tag{205}$$

Note here $p(C)$ is a random variable and $\mathbb{E}(p(C)) = \bar{p} = \mathbb{E}(Y) = K$ is the prevalence. Also note that since $p(C)$ is a nonlinear function of $C$, $\mathbb{E}[\, g(p)\,] \neq g(\mathbb{E}[p])$ for a nonlinear function $g$. We can use Taylor expansion around mean $\mathbb{E}(C) = \mu_C$

$$f(C) = f(\mu_C) + f'(\mu_C)(C - \mu_C) \tag{206}$$

$$f'(C) = \gamma_{\text{logistic}} f(C)(1 - f(C)) \tag{207}$$

$$\gamma_{\text{linear}} = \frac{\text{Cov}(Y, C)}{\text{Var}(C)} \tag{208}$$

$$= \frac{\mathbb{E}(YC) - \mathbb{E}(Y)\mathbb{E}(C)}{\text{Var}(C)}$$

$$= \frac{\mathbb{E}\left[\mathbb{E}(YC \mid C)\right] - \mathbb{E}\left[\mathbb{E}(Y \mid C)\right]\mathbb{E}(C)}{\text{Var}(C)}$$

$$= \frac{\mathbb{E}(Cf(C)) - \mathbb{E}(f(C))\mathbb{E}(C)}{\text{Var}(C)}$$

$$= \frac{\text{Cov}(C, f(C))}{\text{Var}(C)} \tag{209}$$

$$\approx f'(\mu_C)$$

$$= \gamma_{\text{logistic}} p(\mu_C)(1 - p(\mu_C)) \tag{210}$$

Therefore, we got the relationship between $\gamma_{\text{linear}}$ and $\gamma_{\text{logistic}}$. However, we still need to know $\text{Var}(Y \mid C = \mu_C) = p(\mu_C)(1 - p(\mu_C))$ and the best we can do is to use $\text{Var}(Y)$ leading to

$$\gamma_{\text{logistic}} \approx \frac{\gamma_{\text{linear}}}{\text{Var}(Y)} \tag{211}$$

However, empirically we found this approximation can have large biases and thus not recommended. In the following section we will show the source of the biases.

##### Stein's Lemma to link $\gamma_{\text{lin}}$ and $\gamma_{\text{logistic}}$

Since later we'll use it later, re-call that **Stein's Lemma** states that for a normal random variable $X$,

$$\mathbb{E}[g(X)(X - \mathbb{E}(X)] = \mathbb{E}[Xg(X)] - \mathbb{E}(X)\mathbb{E}[g(X)] \tag{212}$$
$$= \text{Var}(X)\mathbb{E}[g'(X)] \tag{213}$$

Comparing with our previous Taylor expansion, eq. (213) replaces the derivative *at a point* — which is all a first-order Taylor expansion supplies — with the *expectation of the derivative* over the whole distribution of $X$. The two coincide only when $g'$ is constant. Also note here we **assumed** clock follows a normal distribution. This assumption may not hold in practice.

We have derived $\text{Cov}(Y, C)$ and instead of Taylor expansion, we can apply Stein's lemma

$$\begin{aligned}\text{Cov}(Y, C) &= \text{Cov}\big(f(C),\, C\big) \\ &= \text{Var}(C)\mathbb{E}[f'(C)] \qquad \text{(by Stein's Lemma)} \\ &= \gamma_{\text{logistic}}\,\text{Var}(C)\mathbb{E}[f(C)(1 - f(C))]\end{aligned} \tag{214}$$

Therefore, we have

$$\gamma_{\text{linear}} = \frac{\operatorname{Cov}(Y,C)}{\operatorname{Var}(C)} = \gamma_{\text{logistic}}\mathbb{E}[f(C)(1-f(C))]$$

$$\gamma_{\text{logistic}} = \frac{\gamma_{\text{linear}}}{\mathbb{E}[f(C)(1-f(C))]} \tag{215}$$

$$\begin{aligned}
\mathbb{E}[f(C)(1-f(C))] &= \mathbb{E}[p(C)(1-p(C))] \\
&= \mathbb{E}[p(C)] - \mathbb{E}[p(C)^2] \\
&= \mathbb{E}[\mathbb{E}(Y \mid C)] - \operatorname{Var}(p(C)) - [\mathbb{E}[p(C)]]^2 \\
&= \mathbb{E}(Y) - \operatorname{Var}(p(C)) - [\mathbb{E}(Y)]^2 \\
&= K(1-K) - \operatorname{Var}(\mathbb{E}(Y \mid C)) && (216) \\
&= \operatorname{Var}(Y) - \operatorname{Var}(\mathbb{E}(Y \mid C)) && (217)
\end{aligned}$$

And now we have an *exact* relationship between $\gamma_{\text{linear}}$ and $\gamma_{\text{logistic}}$:

$$\gamma_{\text{logistic}} = \frac{\gamma_{\text{linear}}}{\operatorname{Var}(Y) - \operatorname{Var}(\mathbb{E}(Y \mid C))} \tag{218}$$

Now it's difficult to estimate $\operatorname{Var}(p(C)) = \operatorname{Var}(\mathbb{E}(Y \mid C))$ using only EWAS summary statistics. But we can find its bound to reduce the bias. From eq. (214), we find $\operatorname{Cov}(Y,C) = \operatorname{Cov}(p(C),C)$, that is,

$$\gamma_{\text{linear}} = \frac{\operatorname{Cov}(Y,C)}{\operatorname{Var}(C)} = \frac{\operatorname{Cov}(p(C),C)}{\operatorname{Var}(C)}$$

Therefore, we can write $p(C) = \gamma_{\text{linear}}C + \text{noise}$ and have

$$\operatorname{Var}(p(C)) = \operatorname{Var}[\mathbb{E}(Y \mid C)] \geq \gamma_{\text{linear}}^2 \operatorname{Var}(C) \tag{219}$$

where the equality holds when $p(C) = \mathbb{E}(Y = 1 \mid C)$ is proportional to $C$ (i.e., noise is 0). Then,

$$\begin{aligned}
\gamma_{\text{logistic}} &\geq \frac{\gamma_{\text{linear}}}{\operatorname{Var}(Y) - \gamma_{\text{linear}}^2 \operatorname{Var}(C)} = \frac{\gamma_{\text{linear}}}{\sigma_e^2} && (220) \\
&= \frac{\gamma_{\text{linear}}}{\operatorname{Var}(Y) - \gamma_{\text{linear}}^2 w^\top \Sigma w} \\
&= \frac{1}{\operatorname{Var}(Y)[1 - \gamma_{\text{linear}}^2 \operatorname{Var}(C)/\operatorname{Var}(Y)]} \frac{w^\top \beta}{w^\top \Sigma w} \operatorname{Var}(Y) \\
&= \frac{w^\top \beta}{w^\top \Sigma w} \frac{1}{1-R^2} && (221)
\end{aligned}$$

where

$$\sigma_e^2 = \text{Var}(Y) - \hat{\gamma}_{\text{linear}}^2 w^\top \Sigma w = \text{Var}(Y)(1 - R^2) \tag{222}$$

$$R^2 = \frac{\hat{\gamma}_{\text{linear}}^2 w^\top \Sigma w}{\text{Var}(Y)} \tag{223}$$

$\sigma_e^2$ is the error variance in the linear regression of $Y = C\gamma + e$, and $R^2$ is the phenotype variance that can be explained by the clock. Practically, we can use

$$\gamma_{\text{logistic}} \approx \frac{\gamma_{\text{linear}}}{\sigma_e^2} \tag{224}$$

to convert $\gamma_{\text{linear}}$ to $\gamma_{\text{logistic}}$ knowing that we may have some downward bias when $\gamma_{\text{linear}} \neq 0$. And the bias becomes stronger when the true $\gamma_{\text{logistic}}$ is large. Now we need to also get its variance. Note that $\hat{\gamma}_{\text{linear}}$ appears in both the numerator and denominator in the formula for $\hat{\gamma}_{\text{logistic}}$, so we use delta method to estimate $\text{Var}(\hat{\gamma}_{\text{logistic}})$.

$$\begin{aligned}
\text{Var}[\hat{\gamma}_{\text{logistic}}] &= \text{Var}[g(\hat{\gamma}_{\text{linear}})] \\
&= \text{Var}[g(\gamma_{\text{linear}}) + g'(\gamma_{\text{linear}})(\hat{\gamma}_{\text{linear}} - \gamma_{\text{linear}})] \\
&= [g'(\gamma_{\text{linear}})]^2 \, \text{Var}(\hat{\gamma}_{\text{linear}}) \\
&= \frac{[\text{Var}(Y) + w^\top \Sigma w \gamma_{\text{linear}}^2]^2}{[\text{Var}(Y) - w^\top \Sigma w \gamma_{\text{linear}}^2]^4} \text{Var}(\hat{\gamma}_{\text{linear}}) \qquad (225) \\
&= \left[ \frac{\text{Var}(Y) + w^\top \Sigma w \gamma_{\text{linear}}^2}{(\sigma_e^2)^2} \right]^2 \text{Var}(\hat{\gamma}_{\text{linear}}) \qquad (226)
\end{aligned}$$

Practically we can plug-in the $\hat{\gamma}_{\text{linear}}$ to equations (224) and (225) to estimate $\hat{\gamma}_{\text{logistic}}$ and its variance.

**SE and P value.** Worth noting that although $\gamma_{\text{logistic}}$ (eq. (224)) is biased towards 0, when $\gamma_{\text{linear}} = 0$, $\gamma_{\text{logistic}} = 0$, thus $H_0 : \gamma_{\text{linear}} = 0$ and $H_0 : \gamma_{\text{logistic}} = 0$ are the same hypothesis. However,

$$\frac{\widehat{\gamma}_{\text{logistic}}}{\text{SE}(\widehat{\gamma}_{\text{logistic}})} = \frac{\widehat{\gamma}_{\text{linear}}}{\text{SE}(\widehat{\gamma}_{\text{linear}})} \frac{\sigma_e^2}{\text{Var}(Y) + w^\top \Sigma w \widehat{\gamma}_{\text{linear}}^2} \tag{227}$$

meaning that the two Z scores are equal only when $\gamma = 0$, and the Z score computed using delta-method SE is attenuated when the association is strong and tends to 0 as $\sigma_e^2 \to 0$ which is not what we want. Therefore, in our method's output, we report the Z score and P value from $\gamma_{\text{lin}}$ for $\gamma_{\text{logistic}}$. The delta-method standard error is still reported, as it remains the first-order measure of the uncertainty in $\hat{\gamma}^{\text{logit}}$; but the reported $Z$ for this parameter will not be equal to $\widehat{\gamma}_{\text{logistic}}/\text{SE}(\widehat{\gamma}_{\text{logistic}})$. Confidence intervals on the log-odds scale should be obtained by mapping the endpoints of the interval for $\gamma_{\text{linear}}$ (eq. (224)) rather than as symmetric Wald intervals.

**Comparing with Taylor expansion.** Comparing eqs. (210) and (218), the Taylor expansion approach makes *two* substitutions, each replacing the correct denominator by a larger one:

$$\mathbb{E}\big[p(C)(1-p(C))\big] \;<\; f(\mu_C)(1-f(\mu_C)) \;<\; K(1-K) = \operatorname{Var} Y \tag{228}$$

Therefore, Taylor expansion *inflate* the denominator leading to *attenuated* estimates for $\gamma_{\text{logistic}}$ toward the null.

##### 3.6 Recovering the individual-level hazard ratio

When a research has individual-level longitudinal data, he/she can compute each sample's epigenetic clock $\boldsymbol{C} = \boldsymbol{M}\boldsymbol{w}$ and conduct Cox regression using individual-level data

$$h(t \mid C_i) = h_0(t) \exp(C_i \Theta_{\text{Cox}}) \,, \tag{229}$$

The question is, whether SARDINE can give an estimator for $\Theta_{\text{Cox}}$.

In section 3.4 we showed that SARDINE can handle summary statistics from a Cox PH model. There, we found that the coefficient of a linear regression of end-of-study case–control status on a single CpG is approximately $\kappa(\pi) := (1-\pi)\,[-\ln(1-\pi)]$ times the log hazard ratio at that CpG. In section 1.4, we have shown that SARDINE can estimate $\gamma$ in a linear regression of $Y = C\gamma + u$. Applying the same arguments and following eqs. (198) and (199), we have

$$\widehat{\Theta}_{\text{Cox}} \approx \frac{\widehat{\gamma}}{(1-\pi)\,[-\ln(1-\pi)]} \tag{230}$$

$$\text{SE}\left(\widehat{\Theta}_{\text{Cox}}\right) \approx \frac{\text{SE}(\widehat{\gamma})}{(1-\pi)\,[-\ln(1-\pi)]} \tag{231}$$

then we can calculate the hazard ratio and confidence interval

$$\widehat{\text{HR}} = \exp\left(\widehat{\Theta}_{\text{Cox}}\right) \tag{232}$$

$$95\%\ \text{CI} = \exp\left(\widehat{\Theta}_{\text{Cox}} \pm 1.96\ \text{SE}\left(\widehat{\Theta}_{\text{Cox}}\right)\right) \tag{233}$$

with $Z$ and the $p$-value inherited unchanged from the linear-scale $\gamma$.

Here $\Theta_{\text{Cox}}$ is the coefficient of $C$ in a Cox model adjusted for the same covariates as the EWAS. Note that $\kappa(\pi)$ is applied to the input summary statistics and then divided out again here, so $\widehat{\Theta}_{\text{Cox}}$ is invariant to the value of $\pi$ supplied by the user; $\widehat{\lambda}$ and $\widehat{\gamma}_{\text{logistic}}$ are not, and remain sensitive to its accuracy. Also worth reminding, $\widehat{\Theta}_{\text{Cox}}$ is a log hazard ratio whereas $\widehat{\gamma}_{\text{logistic}}$ is a log odds ratio for cumulative incidence; the two agree only as $\pi \to 0$.

#### 3.7 Two-stage (residual-based) EWAS design

In a perfect world, for a binary disease outcome or in a longitudinal study on incident disease, we would estimate the clock's effect by fitting a full model (e.g., logistic or Cox PH) with both covariates ($X$) and the clock ($C$) simultaneously to find the true parameter $\Theta$. However, some studies utilized a two-stage design (probably due to computational reasons), where the OLS slope ($b_j$) derived from regressing stage-1 residuals $\boldsymbol{\zeta}$ onto covariate-adjusted methylation. In this two-stage design:

1. Stage 1: residualize both the phenotype and DNAm
   (a) Phenotype is regressed on the covariates alone under its natural link (i.e., logistic for a binary trait, Cox for a time-to-event trait) and a *residual* is extracted
   (b) Similarly, DNAm is regressed on the exactly same set of covariates and a *residual* is extracted
2. Stage 2: for each CpG, these two residuals are used in an ordinary least-squares (OLS) regression to get the final EWAS summary statistics.

The summary statistics released by such a study are therefore neither logistic nor Cox coefficients: they are OLS slopes on a residual scale. This section derives what those slopes estimate and shows that using these summary statistics from the two-stage design, Sardine can recover the clock-level log hazard ratio or log odds ratio from individual-level phenotype-clock regressions. Recall, in section 1.6 we discussed this similar study design for linear regressions. In this section we focus on logistic and Cox regressions since the conclusions are not that obvious.

Throughout this section we assume the *same* covariate set $X$ is used to adjust the phenotype in stage 1 and the methylation in stage 2. The consequences of relaxing this will be discussed later.

**Setup and notations.** Let $X$ be the $N \times K$ covariate matrix (including an intercept) and let $P = \boldsymbol{I}_N - X(X^\top X)^{-1}X^\top$ be the residual-maker of section 1.3, so that $\widetilde{\boldsymbol{M}} = P\boldsymbol{M}$ and $\widetilde{\boldsymbol{C}} = \widetilde{\boldsymbol{M}}\boldsymbol{w}$ are the covariate-adjusted methylation matrix and clock, respectively. Write

$$\sigma^2_{\widetilde{C}} := \operatorname{Var}(\widetilde{C}) = \boldsymbol{w}^\top \Sigma_{\text{adj}} \boldsymbol{w}. \tag{234}$$

**Stage 1: Residualization.** Fit the *null* model containing covariates only, and extract the residuals $\boldsymbol{\zeta} = (\zeta_1, \ldots, \zeta_N)^\top$. For logistic regression:

$$\operatorname{logit} \mathbb{P}(Y_i = 1 \mid X_i) = X_i \eta \tag{235}$$

$$\zeta_i = Y_i - \hat{p}_i = Y_i - \operatorname{logit}^{-1}(X_i \hat{\eta}) \tag{236}$$

For Cox PH:

$$h(t \mid X_i) = h_0(t) \exp(X_i \eta) \tag{237}$$

$$\zeta_i = \delta_i - \hat{H}_0(T_i)\, e^{X_i \hat{\eta}} \tag{238}$$

where $\hat{H}_0$ is the estimated baseline cumulative hazard; $\delta_i$ and $T_i$ are as defined in section 3.4. In the Cox case $\boldsymbol{\zeta}$ is the vector of *martingale residuals*. In the logistic case $\boldsymbol{\zeta}$ must be the *response* residual $\boldsymbol{Y} - \hat{\boldsymbol{p}}$; deviance or Pearson residuals do *not* satisfy the identities below.

**Stage 2: EWAS.** For each CpG $j$, obtain by OLS from linear regression $\boldsymbol{\zeta} \sim \widetilde{\boldsymbol{M}}_j$ of the two residuals

$$b_j = \frac{\operatorname{Cov}(\boldsymbol{\zeta}, \widetilde{\boldsymbol{M}}_j)}{\operatorname{Var}(\widetilde{\boldsymbol{M}}_j)} \tag{239}$$

$$\operatorname{SE}(b_j) \approx \sqrt{\frac{\operatorname{Var}(\boldsymbol{\zeta})}{N \operatorname{Var}(\widetilde{\boldsymbol{M}}_j)}} \tag{240}$$

which are what the study reports in the summary statistics.

**Target parameters.** We wish to recover the coefficient of the clock in the model that an analyst with individual-level data would have fitted,

$$\operatorname{logit} \mathbb{P}(Y = 1 \mid X, C) = X \eta^* + C\, \Theta_{\text{logit}}, \tag{241}$$

$$h(t \mid X, C) = h_0(t) \exp(X \eta^* + C\, \Theta_{\text{Cox}})\,. \tag{242}$$

We write $\Theta$ for either target parameter on the clock scale, and $\vartheta_j$ for the corresponding CpG-level parameter (i.e. $\vartheta_j = \beta_{\text{Cox},j}$ in the Cox case and $\beta_{\text{logistic},j}$ in the logistic regression case). The key insight is that an OLS slope on a score residual *is* a rescaled score statistic.

The derivations for both models involve three CpG-level quantities:

1. $\hat{\vartheta}_j$, the *full-model* estimate: the maximum (partial) likelihood estimate of $\vartheta_j$ in the model that contains $X$ and $\boldsymbol{M}_j$ jointly (eqs. (243) and (295)), i.e., the log odds ratio or log hazard ratio that standard regression software reports from individual-level data. It is the estimate of interest for an EWAS, but the two-stage design never computes it.

2. $b_j$, the *two-stage* slope (eq. (239)), which is what the study reports.

3. $\hat{\vartheta}_j^{(1)}$, the *one-step* estimator: the result of a single Newton–Raphson iteration on the full-model (partial) likelihood, started from the stage-1 fit. Here "step" refers to one iteration of the optimization algorithm, not the stage of the study design. The one-step estimator is not computed in the two-stage analysis; it is an analytical approximation of $\hat{\vartheta}_j$ in closed form.

Later we'll show that $b_j \approx \bar{\nu}\, \hat{\vartheta}_j^{(1)}$ for the logistic model (eq. (279)), where $\bar{\nu}$ is the average fitted Bernoulli variance under the null model; and $b_j \approx \pi\, \hat{\vartheta}_j^{(1)}$ for the Cox model (eq. (336)), where $\pi$ is the event fraction. Thus the slope $b_j$ reported by such a two-stage design is *not* log odds ratio or log hazard ratio. However, the two-stage Wald statistic $b_j/\mathrm{SE}(b_j)$ is asymptotically equivalent to the full-model score test.

##### 3.7.1 Logistic model.

For a binary phenotype, the full model for CpG $j$ contains the covariates and the CpG jointly,

$$\operatorname{logit} \mathbb{P}(Y_i = 1 \mid X_i, M_{ij}) = X_i\eta + M_{ij}\vartheta_j \tag{243}$$

and its log-likelihood is

$$\ell(\eta, \vartheta_j) = \sum_{i=1}^{N} \left[ Y_i(X_i\eta + M_{ij}\vartheta_j) - \log\left(1 + e^{X_i\eta + M_{ij}\vartheta_j}\right)\right] \tag{244}$$

Note, if we define

$$\boldsymbol{\varphi}_j = \begin{pmatrix} \eta \\ \vartheta_j \end{pmatrix} \in \mathbb{R}^{K+1} \tag{245}$$

$$\boldsymbol{D}_j = [X, \boldsymbol{M}_j] \in \mathbb{R}^{N\times(K+1)} \tag{246}$$

Then the linear predictor can be written as $X_i\eta + M_{ij}\vartheta_j = \boldsymbol{D}_{ij}\boldsymbol{\varphi}_j$ and the log-likelihood can be written in the "usual" form

$$\ell(\boldsymbol{\varphi}_j) = \sum_{i=1}^{N} \left[ Y_i(\boldsymbol{D}_{ij}\boldsymbol{\varphi}_j) - \log\left(1 + e^{\boldsymbol{D}_{ij}\boldsymbol{\varphi}_j}\right)\right] \tag{247}$$

The score function is the derivative of the log-likelihood with respect to the coefficients

$$U_{\vartheta_j}(\eta, \vartheta_j) = \frac{\partial \ell}{\partial \vartheta_j} = \sum_{i=1}^{N} M_{ij} \left[Y_i - \text{logit}^{-1}(X_i\eta + M_{ij}\vartheta_j)\right] \tag{248}$$

$$U_{\eta}(\eta, \vartheta_j) = \frac{\partial \ell}{\partial \eta} = \sum_{i=1}^{N} X_i^{\top} \left[Y_i - \text{logit}^{-1}(X_i\eta + M_{ij}\vartheta_j)\right] \tag{249}$$

The full-model MLE estimates $\hat{\boldsymbol{\varphi}}_j = (\hat{\eta}^{\top}, \hat{\vartheta}_j)^{\top}$ can be obtained by setting the above two score equations to be 0. This forces the full-model residuals $\zeta_i^{\text{full}} = Y_i - \hat{p}_i^{\text{full}}$ to be orthogonal to both $\boldsymbol{M}_j$ and $X$. Because these two score functions are nonlinear functions of the parameters, $\hat{\boldsymbol{\varphi}}_j$ has no closed-form solution and must be obtained iteratively.

The two-stage design never fits eq. (243). Stage 1 fit a *null* log-likelihood *without* including CpGs in the model

$$\ell_1(\eta) = \sum_i \left[Y_i X_i \eta - \log\left(1 + e^{X_i\eta}\right)\right] \tag{250}$$

with score

$$\frac{\partial \ell_1}{\partial \eta} = \sum_{i=1}^{N} X_i^{\top} (Y_i - p_i) \tag{251}$$

$$p_i = \text{logit}^{-1}(X_i\eta). \tag{252}$$

The score is 0 at $\hat{\eta}$. Evaluating at $\hat{\eta}$ and recall $\zeta_i = Y_i - \hat{p}_i$, the stage-1 residuals have the following properties:

$$X^{\top}\boldsymbol{\zeta} = \mathbf{0} \tag{253}$$

$$\text{Var}(\boldsymbol{\zeta}) = \frac{1}{N}\boldsymbol{\zeta}^{\top}\boldsymbol{\zeta} \tag{254}$$

$$\sum_{i=1}^{N} \zeta_i = \sum_{i=1}^{N} (Y_i - \hat{p}_i) = 0 \tag{255}$$

$$\bar{\hat{p}} = \bar{Y} \tag{256}$$

$$H\boldsymbol{\zeta} = X(X^{\top}X)^{-1}X^{\top}\boldsymbol{\zeta} = 0 \tag{257}$$

$$P\boldsymbol{\zeta} = \boldsymbol{\zeta} - X(X^{\top}X)^{-1}X^{\top}\boldsymbol{\zeta} = \boldsymbol{\zeta} \tag{258}$$

$$\boldsymbol{\zeta} = P\boldsymbol{\zeta} = P(\boldsymbol{Y} - \hat{\boldsymbol{p}}) = P\boldsymbol{Y} - P\hat{\boldsymbol{p}} = \widetilde{\boldsymbol{Y}} - \widetilde{\boldsymbol{p}} \tag{259}$$

$$HY = H\hat{p} \qquad (\text{since } H\boldsymbol{\zeta} = 0) \tag{260}$$

where $H := X(X^\top X)^{-1}X^\top = \boldsymbol{I}_N - P$ is the hat matrix. At $(\hat\eta,\ \vartheta_j = 0)$, the score $U_{\vartheta_j}(\eta, \vartheta_j)$ becomes

$$U_{\vartheta_j}(\hat\eta, 0) = \sum_i M_{ij}\left(Y_i - \operatorname{logit}^{-1}(X_i\hat\eta)\right) = \sum_i M_{ij}\,\zeta_i = \boldsymbol{M}_j^\top \boldsymbol{\zeta} \tag{261}$$

and we write $U_{\vartheta_j}(0) := U_{\vartheta_j}(\hat\eta, 0)$ as shorthand below. Since $\widetilde{\boldsymbol{M}} = P\boldsymbol{M} = \boldsymbol{M} - X(X^\top X)^{-1}X^\top \boldsymbol{M}$,

$$\boldsymbol{M}_j^\top \boldsymbol{\zeta} = \widetilde{\boldsymbol{M}}_j^\top \boldsymbol{\zeta} + \boldsymbol{M}_j^\top X(X^\top X)^{-1} \underbrace{X^\top \boldsymbol{\zeta}}_{=\boldsymbol{0}} = \widetilde{\boldsymbol{M}}_j^\top \boldsymbol{\zeta} \tag{262}$$

Comparing eq.s (261) and (262) we find

$$U_{\vartheta_j}(0) = \boldsymbol{M}_j^\top \boldsymbol{\zeta} = \widetilde{\boldsymbol{M}}_j^\top \boldsymbol{\zeta} \tag{263}$$

that is, *using the residualized predictor* $\widetilde{\boldsymbol{M}}_j$ *instead of* $\boldsymbol{M}_j$ *does not change the score*. This is the reason the two-stage design works at all: the analyst may residualize methylation freely without disturbing the score, so long as the same covariates were used in stage 1.

Differentiating eq. (248) once more to get the Hessian blocks:

$$\frac{\partial^2 \ell}{\partial\eta\partial\eta^\top} = -\sum_{i=1}^N X_i^\top X_i \frac{e^{X_i\eta + M_{ij}\vartheta_j}}{(1+e^{X_i\eta + M_{ij}\vartheta_j})^2} = -X^\top W X \tag{264}$$

$$\frac{\partial^2 \ell}{\partial\eta\partial\vartheta_j} = -\sum_{i=1}^N X_i^\top M_{ij} \frac{e^{X_i\eta + M_{ij}\vartheta_j}}{(1+e^{X_i\eta + M_{ij}\vartheta_j})^2} = -X^\top W \boldsymbol{M}_j \tag{265}$$

$$\frac{\partial^2 \ell}{\partial\vartheta_j^2} = -\sum_{i=1}^N M_{ij}^2 \frac{e^{X_i\eta + M_{ij}\vartheta_j}}{(1+e^{X_i\eta + M_{ij}\vartheta_j})^2} = -\boldsymbol{M}_j^\top W \boldsymbol{M}_j \tag{266}$$

where

$$\begin{aligned} W &= \operatorname{diag}\left[p_1(1-p_1),\ p_2(1-p_2),\ \dots,\ p_N(1-p_N)\right] \\ &= \operatorname{diag}(\nu_i) \end{aligned} \tag{267}$$

In matrix format:

$$\begin{aligned} \mathcal{H}(\boldsymbol{\varphi}_j) &= -\begin{pmatrix} \sum_i \nu_i X_i^\top X_i & \sum_i \nu_i X_i^\top M_{ij} \\ \sum_i \nu_i M_{ij} X_i & \sum_i \nu_i M_{ij}^2 \end{pmatrix} \\ &= -\begin{pmatrix} X^\top W X & X^\top W \boldsymbol{M}_j \\ \boldsymbol{M}_j^\top W X & \boldsymbol{M}_j^\top W \boldsymbol{M}_j \end{pmatrix} \\ &= -\boldsymbol{D}_j^\top W \boldsymbol{D}_j \end{aligned} \tag{268}$$

and the approximate variances and covariance of $\hat{\boldsymbol{\varphi}}_j$ are obtained from

$$\operatorname{Var}(\hat{\boldsymbol{\varphi}}_j) = [-\mathcal{H}(\hat{\boldsymbol{\varphi}}_j)]^{-1} = \left[\boldsymbol{D}_j^\top W \boldsymbol{D}_j\right]^{-1} = [\mathcal{I}(\hat{\boldsymbol{\varphi}}_j)]^{-1} \tag{269}$$

Since there's no closed-form solution for a logistic regression, an iterative reweighted least squares (IRLS; or Newton-Raphson algorithm) is used to solve for $\hat{\boldsymbol{\varphi}}_j$:

$$\hat{\boldsymbol{\varphi}}_j^{(t+1)} = \hat{\boldsymbol{\varphi}}_j^{(t)} - \left[\mathcal{H}(\hat{\boldsymbol{\varphi}}_j^{(t)})\right]^{-1} U(\hat{\boldsymbol{\varphi}}_j^{(t)}) \tag{270}$$

$$= \hat{\boldsymbol{\varphi}}_j^{(t)} + \left(\boldsymbol{D}_j^\top W^{(t)} \boldsymbol{D}_j\right)^{-1} \boldsymbol{D}_j^\top \left(\boldsymbol{Y} - \boldsymbol{p}^{(t)}\right) \tag{271}$$

where $U(\hat{\boldsymbol{\varphi}}_j^{(t)}) = \boldsymbol{D}_j^\top(\boldsymbol{Y} - \boldsymbol{p}^{(t)})$ is the gradient $\frac{\partial \ell(\boldsymbol{\varphi}_j)}{\partial \boldsymbol{\varphi}_j}$ evaluated at $\hat{\boldsymbol{\varphi}}_j^{(t)}$, and $\boldsymbol{D}_j = [X, \boldsymbol{M}_j] \in \mathbb{R}^{N\times(K+1)}$ is the design matrix as defined in eq. (246). Evaluate the score at the warm start to get the "one-step" estimator:

$$\hat{\boldsymbol{\varphi}}_j^{(0)} = \begin{pmatrix} \hat{\eta} \\ 0 \end{pmatrix} \tag{272}$$

then, $\boldsymbol{Y} - \boldsymbol{p} = \boldsymbol{Y} - \hat{\boldsymbol{p}} = \boldsymbol{\zeta}$, and

$$U(\hat{\boldsymbol{\varphi}}_j^{(0)}) = \boldsymbol{D}_j^\top \boldsymbol{\zeta} = \begin{pmatrix} X^\top \boldsymbol{\zeta} \\ \boldsymbol{M}_j^\top \boldsymbol{\zeta} \end{pmatrix} = \begin{pmatrix} \boldsymbol{0} \\ U_{\vartheta_j}(0) \end{pmatrix} \tag{273}$$

$$\hat{\boldsymbol{\varphi}}_j^{(1)} = \begin{pmatrix} \hat{\eta} \\ 0 \end{pmatrix} + \left[\mathcal{I}(\hat{\boldsymbol{\varphi}}_j^{(0)})\right]^{-1} \begin{pmatrix} \boldsymbol{0} \\ U_{\vartheta_j}(0) \end{pmatrix} \tag{274}$$

$$\begin{aligned} \hat{\vartheta}_j^{(1)} &= 0 + [\mathcal{I}^{-1}]_{21} \times \boldsymbol{0} + [\mathcal{I}^{-1}]_{22} U_{\vartheta_j}(0) \\ &= [\mathcal{I}^{-1}]_{22} U_{\vartheta_j}(0) \\ &= \frac{U_{\vartheta_j}(0)}{\boldsymbol{M}_j^\top W \boldsymbol{M}_j - \boldsymbol{M}_j^\top W X (X^\top W X)^{-1} X^\top W \boldsymbol{M}_j} \\ &= \frac{U_{\vartheta_j}(0)}{\boldsymbol{M}_j^\top W \left[\boldsymbol{I}_N - X(X^\top W X)^{-1} X^\top W\right] \boldsymbol{M}_j} \end{aligned} \tag{275}$$

To simplify $\vartheta_j^{(1)}$ we will use the following assumptions:

1. $\mathbb{E}(M_{ij} \mid X_i)$ is linear in $X_i$. The $W$-weighted and unweighted projections of $\boldsymbol{M}_j$ onto $\operatorname{col}(X)$ then estimate the same conditional mean, so $X(X^\top W X)^{-1} X^\top W \boldsymbol{M}_j - X(X^\top X)^{-1} X^\top \boldsymbol{M}_j = O_p(N^{-1/2})$.
2. $\operatorname{Var}(M_{ij} \mid X_i) = \sigma_m^2$ is constant across $i$ ($M_j$ is conditionally homoscedastic given $X_i$). Since at the warm start $\nu_i = \hat{p}_i(1 - \hat{p}_i) = \nu(X_i)$

is a function of the covariates alone,

$$\frac{1}{N}\sum_{i=1}^{N}\nu_i\widetilde{M}_{ij}^2 \approx \mathbb{E}[\nu(X)\,\mathrm{Var}(M_j\mid X)]$$
$$= \sigma_m^2\,\mathbb{E}[\nu] = \bar{\nu}\,\mathrm{Var}(\widetilde{M}_j) \tag{276}$$

where

$$\bar{\nu} := \frac{1}{N}\sum_{i=1}^{N}\hat{p}_i(1-\hat{p}_i) \tag{277}$$

3. Negligible per-CpG signal. $\hat{\sigma}_{e_j}^2 = \mathrm{Var}(\boldsymbol{\zeta})(1-r_j^2) \approx \mathrm{Var}(\boldsymbol{\zeta})$, where $r_j^2$ is the stage-2 coefficient of determination.

Under assumption 1 the weighted hat matrix may be replaced by $P$, so

$$\vartheta_j^{(1)} \approx \frac{U_{\vartheta_j}(0)}{\boldsymbol{M}_j^\top WP\boldsymbol{M}_j} = \frac{U_{\vartheta_j}(0)}{\widetilde{\boldsymbol{M}}_j^\top W\widetilde{\boldsymbol{M}}_j} = \frac{U_{\vartheta_j}(0)}{\sum_i \nu_i\widetilde{M}_{ij}^2}$$
$$\approx \frac{U_{\vartheta_j}(0)}{N\bar{\nu}\,\mathrm{Var}(\widetilde{M}_j)} = \frac{\widetilde{\boldsymbol{M}}_j^\top\boldsymbol{\zeta}}{N\bar{\nu}\,\mathrm{Var}(\widetilde{M}_j)} = \frac{N\,\mathrm{Cov}(\boldsymbol{\zeta},\widetilde{\boldsymbol{M}}_j)}{N\bar{\nu}\,\mathrm{Var}(\widetilde{M}_j)} \tag{278}$$

where $\boldsymbol{M}_j^\top P\boldsymbol{M}_j = (P\boldsymbol{M}_j)^\top(P\boldsymbol{M}_j) = \widetilde{\boldsymbol{M}}_j^\top\widetilde{\boldsymbol{M}}_j$ since $P$ is symmetric and idempotent. Then we finally get the relationship

$$\boxed{b_j \approx \bar{\nu}\,\hat{\vartheta}_j^{(1)}} \tag{279}$$

where $\bar{\nu} = \sum_{i=1}^{N}\hat{p}_i(1-\hat{p}_i)/N$ (eq. (277)). Thus, the slope $b_j$ reported by a two-stage logistic design is therefore not a log odds ratio. To first order in the effect size, it approximates the full-model log odds ratio $\hat{\vartheta}_j$ multiplied by $\bar{\nu} \le 1/4$, so it is shrunk toward zero, and $b_j/\bar{\nu}$ approximately recovers $\hat{\vartheta}_j$.

From eq. (240) and assumption 3,

$$\mathrm{SE}(b_j) = \left(\hat{\sigma}_{e_j}^2/[N\,\mathrm{Var}(\widetilde{M}_j)]\right)^{1/2} \approx \left(\mathrm{Var}(\boldsymbol{\zeta})/[N\,\mathrm{Var}(\widetilde{M}_j)]\right)^{1/2} \tag{280}$$

To evaluate $\mathrm{Var}(\boldsymbol{\zeta})$, note that $Y_i \in \{0,1\}$ implies $Y_i^2 = Y_i$, so that

$$\mathrm{Var}(Y) = \mathbb{E}(Y^2) - (\mathbb{E}(Y))^2 = \mathbb{E}(Y) - (\mathbb{E}(Y))^2$$
$$= \bar{Y} - \bar{Y}^2 = \bar{Y}(1-\bar{Y}) \tag{281}$$

and since $\bar{\hat{p}} = \bar{Y}$,

$$\begin{aligned}
\bar{\nu} &= \frac{1}{N}\sum_{i=1}^{N} \hat{p}_i(1-\hat{p}_i) = \frac{1}{N}\sum_i \hat{p}_i - \frac{1}{N}\sum_i \hat{p}_i^2 \\
&= \bar{Y} - \frac{1}{N}\sum_i \hat{p}_i^2 = \bar{Y} - (\mathrm{Var}(\hat{p}) + (\bar{\hat{p}})^2) \\
&= \bar{Y}(1-\bar{Y}) - \mathrm{Var}(\hat{p}) \qquad (282)
\end{aligned}$$

The variance of the stage-1 residual is then

$$\begin{aligned}
\mathrm{Var}(\boldsymbol{\zeta}) &= \frac{1}{N}\sum_i \zeta_i^2 - 0 = \frac{1}{N}\sum_i (Y_i - \hat{p}_i)^2 \\
&= \bar{Y} - \frac{2}{N}\sum_i Y_i\hat{p}_i + \frac{1}{N}\sum_i \hat{p}_i^2 \\
&= \bar{Y} - \frac{2}{N}\sum_i (\zeta_i + \hat{p}_i)\hat{p}_i + \frac{1}{N}\sum_i \hat{p}_i^2 \\
&= \bar{Y} - \frac{1}{N}\sum_i \hat{p}_i^2 - \frac{2}{N}\hat{\boldsymbol{p}}^\top\boldsymbol{\zeta} \\
&= \bar{\nu} - \frac{2}{N}\hat{\boldsymbol{p}}^\top\boldsymbol{\zeta} = \bar{\nu} - \frac{2}{N}\hat{\boldsymbol{p}}^\top P\boldsymbol{\zeta} = \bar{\nu} - \frac{2}{N}\widetilde{\boldsymbol{p}}^\top\boldsymbol{\zeta} \\
&= \bar{\nu} - 2\,\mathrm{Cov}(\widetilde{\boldsymbol{p}}, \boldsymbol{\zeta}) \qquad (283) \\
&= \bar{\nu} + O_p(N^{-1/2}) \qquad (284)
\end{aligned}$$

where $\widetilde{\boldsymbol{p}} = P\hat{\boldsymbol{p}}$, and the last step holds because $\mathbb{E}(\zeta_i \mid X_i) = 0$ when the null logistic model is correctly specified. Therefore,

$$\mathrm{SE}\big(\hat{\vartheta}_j^{(1)}\big) = \big([\mathcal{I}^{-1}]_{22}\big)^{1/2} \approx \Big(N\bar{\nu}\,\mathrm{Var}(\widetilde{M}_j)\Big)^{-1/2} \qquad (285)$$

$$\mathrm{SE}(b_j) \approx \big(\bar{\nu}/[N\,\mathrm{Var}(\widetilde{M}_j)]\big)^{1/2} \qquad (286)$$

and

$$\begin{aligned}
\frac{\hat{\vartheta}_j^{(1)}}{\mathrm{SE}\big(\hat{\vartheta}_j^{(1)}\big)} &\approx \frac{b_j}{\bar{\nu}}\sqrt{N\bar{\nu}\,\mathrm{Var}(\widetilde{M}_j)} = b_j\sqrt{\frac{N\,\mathrm{Var}(\widetilde{M}_j)}{\bar{\nu}}} \\
&\approx \frac{b_j}{\mathrm{SE}(b_j)} \qquad (287)
\end{aligned}$$

Thus, the two-stage Wald statistic $b_j/\mathrm{SE}(b_j)$ is asymptotically equivalent to the full-model score test. The $Z$ statistics and $P$ values reported by the two-stage design thus approximate those of the full model.

**Recovering the clock-level parameter** Here we'll show that using summary statistics obtained by this two-stage study design, SARDINE existing pipeline can already recover the clock-level log odds ratio.

Since stage 2 is a linear regression of $\boldsymbol{\zeta} \sim \widetilde{M}_j$, eq.s (114) and (115) in section 3.2 can be applied to convert the effect size to the $\widetilde{M}_j \sim \boldsymbol{\zeta}$ direction:

$$b_j^2 + (N-2)\,\mathrm{SE}(b_j)^2 = \frac{\mathrm{Var}(\boldsymbol{\zeta})}{\mathrm{Var}(\widetilde{M}_j)} \tag{288}$$

$$\begin{aligned} \widehat{\beta}_j &= \frac{b_j}{b_j^2 + (N-2)\,\mathrm{SE}(b_j)^2} = \frac{b_j\,\mathrm{Var}(\widetilde{M}_j)}{\mathrm{Var}(\boldsymbol{\zeta})} \\ &= \frac{\mathrm{Cov}(\boldsymbol{\zeta}, \widetilde{\boldsymbol{M}}_j)}{\mathrm{Var}(\boldsymbol{\zeta})} \end{aligned} \tag{289}$$

where $\mathrm{Var}(\boldsymbol{\zeta})$ is estimated by eq.(28).

To compare with the individual-level analysis, write $\widetilde{\boldsymbol{Y}} = P\boldsymbol{Y}$, so that $\boldsymbol{\zeta} = \widetilde{\boldsymbol{Y}} - \widetilde{\boldsymbol{p}}$. By eq. (284), the variance of the residualized phenotype is

$$\begin{aligned} \mathrm{Var}(\widetilde{Y}) &= \mathrm{Var}(\boldsymbol{\zeta} + \widetilde{\boldsymbol{p}}) \\ &= \mathrm{Var}(\boldsymbol{\zeta}) + 2\,\mathrm{Cov}(\boldsymbol{\zeta}, \widetilde{\boldsymbol{p}}) + \mathrm{Var}(\widetilde{\boldsymbol{p}}) \\ &= \bar{\nu} - 2\,\mathrm{Cov}(\widetilde{\boldsymbol{p}}, \boldsymbol{\zeta}) + 2\,\mathrm{Cov}(\boldsymbol{\zeta}, \widetilde{\boldsymbol{p}}) + \mathrm{Var}(\widetilde{\boldsymbol{p}}) \\ &= \bar{\nu} + \mathrm{Var}(\widetilde{\boldsymbol{p}}) \end{aligned} \tag{290}$$

The effect of phenotype on clock given by SARDINE is then

$$\begin{aligned} \widehat{\lambda} = \boldsymbol{w}^\top \widehat{\boldsymbol{\beta}} &= \frac{1}{\mathrm{Var}(\boldsymbol{\zeta})} \sum_{j=1}^{J} w_j\,\mathrm{Cov}(\boldsymbol{\zeta}, \tilde{\boldsymbol{M}}_j) \\ &= \frac{\mathrm{Cov}(\boldsymbol{\zeta}, \widetilde{\boldsymbol{M}\boldsymbol{w}})}{\mathrm{Var}(\boldsymbol{\zeta})} = \frac{\mathrm{Cov}(\boldsymbol{\zeta}, \widetilde{\boldsymbol{C}})}{\mathrm{Var}(\boldsymbol{\zeta})} \\ &= \frac{\mathrm{Cov}(\boldsymbol{Y} - \hat{\boldsymbol{p}}, \widetilde{\boldsymbol{C}})}{\mathrm{Var}(\boldsymbol{\zeta})} = \frac{\mathrm{Cov}(\widetilde{\boldsymbol{Y}} - \widetilde{\boldsymbol{p}}, \widetilde{\boldsymbol{C}})}{\mathrm{Var}(\boldsymbol{\zeta})} \\ &= \frac{\mathrm{Cov}(\widetilde{\boldsymbol{Y}}, \widetilde{\boldsymbol{C}}) - \mathrm{Cov}(\widetilde{\boldsymbol{p}}, \widetilde{\boldsymbol{C}})}{\mathrm{Var}(\widetilde{\boldsymbol{\zeta}})} \\ &= \frac{\mathrm{Cov}(\widetilde{\boldsymbol{Y}}, \widetilde{\boldsymbol{C}})}{\mathrm{Var}(\widetilde{\boldsymbol{Y}})} \frac{\mathrm{Var}(\widetilde{\boldsymbol{Y}})}{\mathrm{Var}(\widetilde{\boldsymbol{\zeta}})} + \frac{O_p(N^{-1/2})}{\mathrm{Var}(\widetilde{\boldsymbol{\zeta}})} \\ &= \hat{\lambda}_{\text{ind}} \frac{\bar{\nu} + \mathrm{Var}(\widetilde{\boldsymbol{p}})}{\bar{\nu} + O_p(N^{-1/2})} + \frac{O_p(N^{-1/2})}{\bar{\nu} + O_p(N^{-1/2})} \end{aligned}$$

thus, we have

$$\boxed{\widehat{\lambda} \approx \hat{\lambda}_{\text{ind}} \left(1 + \frac{\operatorname{Var}(\widetilde{\boldsymbol{p}})}{\bar{\nu}}\right) \geq \hat{\lambda}_{\text{ind}}} \tag{291}$$

where $\hat{\lambda}_{\text{ind}}$ is the regression coefficient in individual-level linear regression $\widetilde{C} \sim \widetilde{Y}$. This shows that $\hat{\lambda}$ from a two-stage design is systematically inflated relative to the covariate-adjusted linear slope.

The log odds ratio of the clock on phenotype given by SARDINE is given by eq.s (17), (223), and (224):

$$\widehat{\gamma}_{\text{logistic}} = \frac{\widehat{\lambda}}{\boldsymbol{w}^\top \Sigma_{\text{adj}} \boldsymbol{w}} \frac{1}{1 - R^2} = \frac{\boldsymbol{w}^\top \widehat{\boldsymbol{\beta}}}{\boldsymbol{w}^\top \Sigma_{\text{adj}} \boldsymbol{w}} \frac{1}{1 - R^2} \tag{292}$$

Compare it with the log odds ratio from the individual-level logistic regression

$$\operatorname{logit}(\mathbb{P}(Y_i = 1 \mid X_i, C_i)) = X_i \eta^* + C_i \Theta \tag{293}$$

The "one-step" derivation for $\hat{\Theta}$ is very similar to those on the CpG level (eq.(278)) we just discussed (also note that the residuals under the null $\boldsymbol{\zeta}$ are identical with those in Stage 1 from CpG level).

$$\begin{aligned}
\widehat{\Theta}^{(1)} &\approx \frac{\operatorname{Cov}(\widetilde{C}, \boldsymbol{\zeta})}{\bar{\nu} \operatorname{Var}(\widetilde{C})} \approx \frac{\operatorname{Cov}(\widetilde{C}, \boldsymbol{\zeta})}{\operatorname{Var}(\boldsymbol{\zeta}) \operatorname{Var}(\widetilde{C})} \\
&= \frac{\hat{\lambda}}{\operatorname{Var}(\widetilde{C})} = \frac{\boldsymbol{w}^\top \widehat{\boldsymbol{\beta}}}{\boldsymbol{w}^\top \Sigma_{\text{adj}} \boldsymbol{w}}
\end{aligned}$$

thus we have

$$\boxed{\widehat{\Theta}^{(1)} \approx \hat{\gamma}_{\text{logistic}}(1 - R^2)} \tag{294}$$

Therefore, we find that the log odd ratios from SARDINE is an approximate of the one-step estimator for the log odds ratio in the individual-level logistic regression. That is, even though the input summary statistics come from a two-stage design, SARDINE can still recover the regression coefficient from the individual-level regressions with covariates. Worth mentioning is that, $\widehat{\Theta}^{(1)}$ is a single Newton step from the null and is itself attenuated toward zero relative to the MLE $\widehat{\Theta}_{\text{MLE}}$. Thus $\widehat{\Theta}_{\text{MLE}}$ and $\hat{\gamma}_{\text{logistic}}$ could be closer to each other than eq.(294) suggests.

##### 3.7.2 Cox model

We now apply the "two-stage" design to incident disease in a longitudinal study, where stage 1 is a Cox proportional hazards (PH) regression [9]. The three CpG-level quantities and the three-part argument stated in the beginning of this section carry over. Here $\hat{\vartheta}_j$ is the maximum partial likelihood estimate of the log hazard ratio in the full Cox model that contains $X$ and $\boldsymbol{M}_j$ jointly (eq. (295)), and the factor relating $b_j$ to $\hat{\vartheta}_j^{(1)}$ is the event fraction $\pi$, the proportion of incident cases at the end of follow-up (eq. (336)).

Write the Cox PH model for CpG $j$ as

$$\begin{aligned} h(t \mid X_i, M_{ij}) &= h_0(t)\exp(X_i\eta + M_{ij}\vartheta_j) \\ &= h_0(t)\exp(\boldsymbol{D}_{ij}\boldsymbol{\varphi}_j) \\ &= h_0(t)\exp(\psi_i) = h_i(t) \end{aligned} \tag{295}$$

with the same stacked parameter and design matrix as before,

$$\boldsymbol{\varphi}_j = \begin{pmatrix} \eta \\ \vartheta_j \end{pmatrix} \in \mathbb{R}^{K+1} \tag{296}$$

$$\boldsymbol{D}_j = [X, \boldsymbol{M}_j] \in \mathbb{R}^{N\times(K+1)} \tag{297}$$

$$\psi_i := \boldsymbol{D}_{ij}\boldsymbol{\varphi}_j \tag{298}$$

Note, a Cox model has no free intercept within the exponential term: a constant shift of the linear predictor is absorbed into $h_0(t)$. The intercept column of $X$ therefore drops out of the partial likelihood below (its coefficient can be fixed at 0), and in the information matrix $X$ refers to the covariates without the intercept column. The residual-maker $P$ is still built from $X$ *with* the intercept.

The Cox model is semi-parametric since it contains both a parametric component (coefficients $\boldsymbol{\varphi}_j$) and a non-parametric component (the baseline hazard $h_0(t)$). Let $\mathcal{R}(t) = \{k : T_k \geq t\}$ denote the risk set at time $t$, which is the set of all individuals who have not yet experienced the event or been censored just before time $t$.

In a small time interval $[t_i, t_i + \Delta t]$, the probability that individual $i$ has the event is approximately $h_i(t_i)\Delta t$. Therefore, the conditional probability that individual $i$ fails, given that one failure occurs in the risk set $\mathcal{R}(t_i)$, is the ratio of their hazard to the sum of the hazards of everyone at risk:

$$\begin{aligned}
&\mathbb{P}(\text{individual } i \text{ fails} \mid \text{one failure in } \mathcal{R}(t_i)) \\
&\qquad = \frac{h_i(t_i)}{\sum_{k\in\mathcal{R}(t_i)} h_k(t_i)} = \frac{h_0(t_i)\exp(\psi_i)}{\sum_{k\in\mathcal{R}(t_i)} h_0(t_i)\exp(\psi_k)} \\
&\qquad = \frac{\exp(\psi_i)}{\sum_{k\in\mathcal{R}(t_i)} \exp(\psi_k)} \qquad (299)
\end{aligned}$$

To find the joint probability of all observed events, we take the product of these conditional probabilities over all individuals who actually experienced the event:

$$L(\boldsymbol{\varphi}_j) = \prod_{i=1}^{N} \left[ \frac{\exp(\psi_i)}{\sum_{k\in\mathcal{R}(t_i)} \exp(\psi_k)} \right]^{\delta_i} \qquad (300)$$

which is the *partial* likelihood (since it does not involve the baseline hazard $h_0(t)$) proposed by Cox [9, 10] to estimate the parameters. (Recall eq. (170), $\delta_i$ is the binary end-of-study incident disease status indicator.) Taking log,

$$\begin{aligned}
\ell(\boldsymbol{\varphi}_j) &= \sum_{i=1}^{N} \delta_i \left[ \psi_i - \log \sum_{k\in\mathcal{R}(T_i)} e^{\psi_k} \right] \\
&= \sum_{i=1}^{N} \delta_i \left[ \boldsymbol{D}_{ij}\boldsymbol{\varphi}_j - \log \sum_{k\in\mathcal{R}(T_i)} e^{\boldsymbol{D}_{kj}\boldsymbol{\varphi}_j} \right] \qquad (301)
\end{aligned}$$

This is the objective function we maximize to estimate $\boldsymbol{\varphi}_j$. The corresponding partial score function is

$$U(\boldsymbol{\varphi}_j) = \frac{\partial \ell}{\partial \boldsymbol{\varphi}_j} = \sum_{i=1}^{N} \delta_i \left[ \boldsymbol{D}_{ij} - \bar{\boldsymbol{D}}(T_i) \right]^{\top} \qquad (302)$$

where

$$\bar{\boldsymbol{D}}(t) = \frac{\sum_{k\in\mathcal{R}(t)} \boldsymbol{D}_{kj} \exp(\boldsymbol{D}_{kj}\boldsymbol{\varphi}_j)}{\sum_{k\in\mathcal{R}(t)} \exp(\boldsymbol{D}_{kj}\boldsymbol{\varphi}_j)} \qquad (303)$$

The two blocks of the score are

$$U_{\vartheta_j}(\eta, \vartheta_j) = \sum_{i=1}^{N} \delta_i \left[ M_{ij} - \bar{M}_j(T_i) \right] \qquad (304)$$

$$U_{\eta}(\eta, \vartheta_j) = \sum_{i=1}^{N} \delta_i \left[ X_i - \bar{X}(T_i) \right]^{\top} \qquad (305)$$

The full-model maximum partial likelihood estimator $\widehat{\boldsymbol{\varphi}}_j$ solves the equation $U(\widehat{\boldsymbol{\varphi}}_j) = 0$. However, there's no closed-form solution; it must be obtained numerically. Differentiating once more to get the Hessian matrix,

$$\mathcal{H}(\boldsymbol{\varphi}_j) = \frac{\partial^2 \ell}{\partial \boldsymbol{\varphi}_j \partial \boldsymbol{\varphi}_j^\top} = -\sum_{i=1}^{N} \delta_i \left[ \frac{S^{(2)}(T_i)}{S^{(0)}(T_i)} - \bar{\boldsymbol{D}}(T_i)^\top \bar{\boldsymbol{D}}(T_i) \right] \tag{306}$$

where we denote

$$S^{(0)}(t) = \sum_{k \in \mathcal{R}(t)} e^{\psi_k} \tag{307}$$

$$S^{(2)}(t) = \sum_{k \in \mathcal{R}(t)} \boldsymbol{D}_{kj}^\top \boldsymbol{D}_{kj} e^{\psi_k} \tag{308}$$

so the observed information is

$$\mathcal{I}(\boldsymbol{\varphi}_j) = -\mathcal{H}(\boldsymbol{\varphi}_j) = \sum_{i:\, \delta_i = 1} \mathbb{V}(T_i) \tag{309}$$

$$\mathbb{V}(t) = \begin{pmatrix} \mathrm{Var}_{\mathcal{R}(t)}(X) & \mathrm{Cov}_{\mathcal{R}(t)}(X, M_j) \\ \mathrm{Cov}_{\mathcal{R}(t)}(M_j, X) & \mathrm{Var}_{\mathcal{R}(t)}(M_j) \end{pmatrix} \tag{310}$$

where $\mathrm{Var}_{\mathcal{R}(t)}$ and $\mathrm{Cov}_{\mathcal{R}(t)}$ denote variances and covariances computed over the risk set $\mathcal{R}(t)$, with individual $k$ weighted by $e^{\psi_k}$. As in the logistic case there is no closed-form solution, and $\widehat{\boldsymbol{\varphi}}_j$ is obtained by an iterative method such as Newton–Raphson.

The two-stage design of section 3.7 never fits the full model eq. (295). It first fits a Cox regression *without* CpGs and computes the martingale residuals (stage 1), and then regresses the martingale residuals on each residualized CpG by OLS (stage 2). We show below that stage 1 already yields the full-model score for $\vartheta_j$, and that stage 2 only rescales this score.

Stage 1 maximizes eq. (301) with $\vartheta_j = 0$ (covariates only), giving $\hat{\eta}$. Given $\hat{\eta}$, the baseline cumulative hazard is estimated by the Breslow estimator [11]

$$\widehat{H}_0(t) = \sum_{i:\, T_i \le t} \frac{\delta_i}{S^{(0)}(T_i; \hat{\eta})} \tag{311}$$

$$S^{(0)}(t; \hat{\eta}) = \sum_{k \in \mathcal{R}(t)} e^{X_k \hat{\eta}} \tag{312}$$

Barlow and Prentice [12] introduced **martingale residuals** for an individual $i$ as the difference between observed number of events and his/her expected number of events (or, as excess deaths)

$$\zeta_i = \delta_i - \hat{H}_0(T_i)e^{X_i\hat{\eta}} = \delta_i - \hat{\Lambda}_i \tag{313}$$

as in eq. (238). Here we denote the fitted cumulative hazard of individual $i$ at the end of his or her follow-up as

$$\hat{\Lambda}_i := \hat{H}_0(T_i)\, e^{X_i\hat{\eta}} \tag{314}$$

for the number of events individual $i$ is expected to have over his or her follow-up under the null model. We use $N_E$ to denote the number of events and $\pi$ for the event fraction, i.e., the proportion of cases at the end of follow-up (Section 3.4)

$$N_E = \sum_{i=1}^{N} \delta_i \tag{315}$$

$$\pi = N_E/N \tag{316}$$

The martingale residuals have properties similar to those of the logistic stage-1 properties we derived previously [13]. All of them follow from a single identity. Let $\boldsymbol{Z} = (Z_1, \ldots, Z_N)^\top$ be any variable measured on the $N$ individuals (a covariate, a CpG, or the clock). The partial score for adding $\boldsymbol{Z}$ to the stage-1 model, evaluated at the stage-1 fit (coefficient of $\boldsymbol{Z}$ equal to 0), is

$$U_Z(\hat{\eta}, 0) = \sum_{i=1}^{N} \delta_i \left[Z_i - \bar{Z}(T_i)\right] \tag{317}$$

$$\bar{Z}(t) = \frac{\sum_{k\in\mathcal{R}(t)} Z_k\, e^{X_k\hat{\eta}}}{S^{(0)}(t;\hat{\eta})} \tag{318}$$

and

$$\begin{aligned}\sum_{i=1}^{N} \delta_i \bar{Z}(T_i) &= \sum_{i=1}^{N} \delta_i \sum_{k:T_k\geq T_i} \frac{Z_k e^{X_k\hat{\eta}}}{S^{(0)}(T_i;\hat{\eta})} \\ &= \sum_{k=1}^{N} Z_k e^{X_k\hat{\eta}} \sum_{i:T_i\leq T_k} \frac{\delta_i}{S^{(0)}(T_i;\hat{\eta})} \\ &= \sum_{k=1}^{N} Z_k e^{X_k\hat{\eta}} \widehat{H}_0(T_k) \\ &= \sum_{k=1}^{N} Z_k \hat{\Lambda}_k \end{aligned} \tag{319}$$

where the third line uses the Breslow estimator eq. (311) and the last line uses eq. (314). We exchanged the summation because a pair $(i, k)$ enters the sum exactly when $\delta_i = 1$ and $T_i \le T_k$, that is, when $k$ is still at risk at the time $i$ fails. Substituting back into eq. (317),

$$U_Z(\hat{\eta}, 0) = \sum_{i=1}^{N} Z_i \delta_i - \sum_{i=1}^{N} Z_i \hat{\Lambda}_i = \sum_{i=1}^{N} Z_i \, \zeta_i = \boldsymbol{Z}^\top \boldsymbol{\zeta} \tag{320}$$

Applying eq. (320) to different choices of $\boldsymbol{Z}$ gives the stage-1 properties:

- $\boldsymbol{Z} = \mathbf{1}$: then $\bar{Z}(t) = 1$ for every $t$, so $\mathbf{1}^\top \boldsymbol{\zeta} = \sum_i \delta_i (1 - 1) = 0$.
- $\boldsymbol{Z}$ = a non-intercept column of $X$: then $U_Z(\hat{\eta}, 0)$ is a component of the stage-1 score, which is zero at $\hat{\eta}$. Together with the previous item, $X^\top \boldsymbol{\zeta} = \mathbf{0}$.

Hence

$$\sum_{i=1}^{N} \zeta_i = 0 \tag{321}$$

$$X^\top \boldsymbol{\zeta} = \mathbf{0} \tag{322}$$

$$\frac{1}{N} \sum_{i=1}^{N} \hat{\Lambda}_i = \frac{1}{N} \sum_{i=1}^{N} \delta_i = \pi \tag{323}$$

$$P \boldsymbol{\zeta} = \boldsymbol{\zeta} \tag{324}$$

And taking $\boldsymbol{Z} = \boldsymbol{M}_j$ gives the Cox analogue of eq. (261); because $X^\top \boldsymbol{\zeta} = \mathbf{0}$, the residualization argument of eq. (262) also applies:

$$U_{\vartheta_j}(0) := U_{\vartheta_j}(\hat{\eta}, 0) = \boldsymbol{M}_j^\top \boldsymbol{\zeta} = \widetilde{\boldsymbol{M}}_j^\top \boldsymbol{\zeta} \tag{325}$$

As in the logistic case, using the residualized $\widetilde{\boldsymbol{M}}_j$ in stage 2 does not change the score, provided the same covariates were used in stage 1.

We take one Newton–Raphson step from the warm start

$$\hat{\boldsymbol{\varphi}}_j^{(0)} = \begin{pmatrix} \hat{\eta} \\ 0 \end{pmatrix} \tag{326}$$

By eqs. (322) and (325), the score at the warm start is

$$U(\hat{\boldsymbol{\varphi}}_j^{(0)}) = \begin{pmatrix} X^\top \boldsymbol{\zeta} \\ \boldsymbol{M}_j^\top \boldsymbol{\zeta} \end{pmatrix} = \begin{pmatrix} \mathbf{0} \\ U_{\vartheta_j}(0) \end{pmatrix} \tag{327}$$

and exactly as in the logistic case,

$$\hat{\boldsymbol{\varphi}}_j^{(1)} = \begin{pmatrix} \hat{\eta} \\ 0 \end{pmatrix} + \left[\mathcal{I}(\hat{\boldsymbol{\varphi}}_j^{(0)})\right]^{-1} \begin{pmatrix} \mathbf{0} \\ U_{\vartheta_j}(0) \end{pmatrix} \tag{328}$$

$$\begin{aligned} \hat{\vartheta}_j^{(1)} &= [\mathcal{I}^{-1}]_{22}\, U_{\vartheta_j}(0) \\ &= \frac{U_{\vartheta_j}(0)}{\mathcal{I}_{MM} - \mathcal{I}_{MX}\,\mathcal{I}_{XX}^{-1}\,\mathcal{I}_{XM}} \end{aligned} \tag{329}$$

where the denominator terms are from eq. (310) evaluated at $\hat{\boldsymbol{\varphi}}_j^{(0)}$,

$$\mathcal{I}_{XX} = \sum_{i:\,\delta_i=1} \mathrm{Var}_{\mathcal{R}(T_i)}(X) \tag{330}$$

$$\mathcal{I}_{XM} = \mathcal{I}_{MX}^{\top} = \sum_{i:\,\delta_i=1} \mathrm{Cov}_{\mathcal{R}(T_i)}(X, M_j) \tag{331}$$

$$\mathcal{I}_{MM} = \sum_{i:\,\delta_i=1} \mathrm{Var}_{\mathcal{R}(T_i)}(M_j) \tag{332}$$

The denominator of eq. (329) is the information about $\vartheta_j$ that remains after accounting for $X$. It is unchanged if $\boldsymbol{M}_j$ is replaced by $\widetilde{\boldsymbol{M}}_j$:

$$\mathcal{I}_{MM} - \mathcal{I}_{MX}\,\mathcal{I}_{XX}^{-1}\,\mathcal{I}_{XM} = \sum_{i:\,\delta_i=1} \mathrm{Var}_{\mathcal{R}(T_i)}(\widetilde{M}_j) - \mathcal{I}_{\widetilde{M}X}\,\mathcal{I}_{XX}^{-1}\,\mathcal{I}_{X\widetilde{M}} \tag{333}$$

where $\mathcal{I}_{X\widetilde{M}} = \mathcal{I}_{\widetilde{M}X}^{\top} = \sum_{i:\,\delta_i=1} \mathrm{Cov}_{\mathcal{R}(T_i)}(X, \widetilde{M}_j)$.

To simplify eq. (333) we use assumptions 1–3 of the logistic case (linear $\mathbb{E}(M_{ij} \mid X_i)$, conditional homoscedasticity, negligible per-CpG signal) and *additional* assume a non-informative censoring: Given $X_i$, the censoring time of individual $i$ does not depend on $M_{ij}$. Under these assumptions, $\mathbb{E}(\widetilde{M}_{ij} \mid X_i) = 0$ and $\widetilde{M}_j$ has mean $\approx 0$ and is uncorrelated with $X$ within every risk set, so the cross term $\mathcal{I}_{\widetilde{M}X}\mathcal{I}_{XX}^{-1}\mathcal{I}_{X\widetilde{M}}$ is negligible. Also, the variance of $\widetilde{M}_j$ within every risk set equals its variance in the whole sample, $\mathrm{Var}_{\mathcal{R}(t)}(\widetilde{M}_j) \approx \sigma_m^2 = \mathrm{Var}(\widetilde{M}_j)$. Therefore

$$\begin{aligned} \mathcal{I}_{MM} - \mathcal{I}_{MX}\,\mathcal{I}_{XX}^{-1}\,\mathcal{I}_{XM} &\approx \sum_{i:\,\delta_i=1} \mathrm{Var}(\widetilde{M}_j) \\ &= N_E\,\mathrm{Var}(\widetilde{M}_j) \\ &= \pi N\,\mathrm{Var}(\widetilde{M}_j) \end{aligned} \tag{334}$$

Substituting eqs. (325) and (334) into eq. (329),

$$\hat{\vartheta}_j^{(1)} \approx \frac{U_{\vartheta_j}(0)}{N\pi \operatorname{Var}(\widetilde{M}_j)} = \frac{\widetilde{\boldsymbol{M}}_j^\top \boldsymbol{\zeta}}{N\pi \operatorname{Var}(\widetilde{M}_j)} = \frac{N \operatorname{Cov}(\boldsymbol{\zeta}, \widetilde{\boldsymbol{M}}_j)}{N\pi \operatorname{Var}(\widetilde{M}_j)} \tag{335}$$

and comparing with the effect size from the two-stage approach $b_j$ (eq. (239)),

$$\boxed{b_j \approx \pi\, \hat{\vartheta}_j^{(1)}} \tag{336}$$

Therefore, the effect size $b_j$ reported by a two-stage Cox design is *not* log hazard ratios. Under the assumptions above and to first order in the effect size, it approximates the full-model log hazard ratio $\hat{\vartheta}_j$ multiplied by $\pi = N_E/N$, the proportion of incident cases at the end of follow-up. Consequently, $b_j$ is shrunk toward zero by the factor $\pi$ (compared with $\hat{\vartheta}_j$), and $b_j/\pi$ approximately recovers $\hat{\vartheta}_j$.

From eq. (240),

$$\operatorname{SE}(b_j) = \left(\hat{\sigma}_{e_j}^2 / [N \operatorname{Var}(\widetilde{M}_j)]\right)^{1/2} \approx \left(\operatorname{Var}(\boldsymbol{\zeta}) / [N \operatorname{Var}(\widetilde{M}_j)]\right)^{1/2} \tag{337}$$

The variance of martingale residual can be approximated as [13, 14]

$$\operatorname{Var}(\boldsymbol{\zeta}) \approx \mathbb{E}(\delta_i) = \pi \tag{338}$$

which parallels $\operatorname{Var}(\boldsymbol{\zeta}) \approx \bar{\nu}$ in the logistic case (eq. (284)). Note that it is $\pi$, not $\pi(1-\pi)$: $\zeta_i$ is the residual of an event count over a person-specific follow-up time, not of a binary outcome. Hence

$$\operatorname{SE}\big(\hat{\vartheta}_j^{(1)}\big) = \left([\mathcal{I}^{-1}]_{22}\right)^{1/2} \approx \frac{1}{\sqrt{N\pi \operatorname{Var}(\widetilde{M}_j)}} \tag{339}$$

$$\operatorname{SE}(b_j) \approx \sqrt{\frac{\pi}{N \operatorname{Var}(\widetilde{M}_j)}} \tag{340}$$

and

$$\begin{aligned} \frac{\hat{\vartheta}_j^{(1)}}{\operatorname{SE}\big(\hat{\vartheta}_j^{(1)}\big)} &\approx \frac{b_j}{\pi}\sqrt{N\pi \operatorname{Var}(\widetilde{M}_j)} = b_j \sqrt{\frac{N \operatorname{Var}(\widetilde{M}_j)}{\pi}} \\ &\approx \frac{b_j}{\operatorname{SE}(b_j)} \end{aligned} \tag{341}$$

Therefore, as in the logistic case, the two-stage Wald statistic $b_j/\operatorname{SE}(b_j)$ is asymptotically equivalent to the score test for $H_0 : \vartheta_j = 0$ in the full Cox model. The $Z$ statistics and $P$ values reported by the two-stage design thus approximate those of the full model.

**Recovering the clock-level parameter.** Here we'll show that using summary statistics obtained by this two-stage study design, SARDINE existing pipeline can already recover the clock-level log hazard ratio.

Since stage 2 here is again a linear regression $\boldsymbol{\zeta} \sim \widetilde{\boldsymbol{M}}_j$, similar to the derivations for the binary outcome in this section, the conversion of eq. (289) applies unchanged,

$$\widehat{\beta}_j = \frac{b_j}{b_j^2 + (N-2)\,\mathrm{SE}(b_j)^2} = \frac{\mathrm{Cov}(\boldsymbol{\zeta}, \widetilde{\boldsymbol{M}}_j)}{\mathrm{Var}(\boldsymbol{\zeta})} \tag{342}$$

and eq. (28) estimates the variance of the phenotype, here $\mathrm{Var}(\boldsymbol{\zeta})$. With $\boldsymbol{w}^\top \Sigma_{\mathrm{adj}} \boldsymbol{w} = \mathrm{Var}(\widetilde{C})$ (eq. (234)), the two outputs are

$$\widehat{\lambda} = \boldsymbol{w}^\top \widehat{\boldsymbol{\beta}} = \frac{\mathrm{Cov}(\boldsymbol{\zeta}, \widetilde{\boldsymbol{C}})}{\mathrm{Var}(\boldsymbol{\zeta})} \tag{343}$$

$$\hat{\gamma} = \frac{\widehat{\lambda}\ \mathrm{Var}(\boldsymbol{\zeta})}{\boldsymbol{w}^\top \Sigma_{\mathrm{adj}} \boldsymbol{w}} = \frac{\mathrm{Cov}(\boldsymbol{\zeta}, \widetilde{\boldsymbol{C}})}{\mathrm{Var}(\widetilde{C})} \tag{344}$$

Thus $\hat{\gamma}$ is the least-squares slope of the stage-1 martingale residual on the covariate-adjusted clock, i.e., the phenotype-on-clock regression that SARDINE targets, and $\widehat{\lambda}$ is the slope in the reverse direction.

Replacing $\boldsymbol{M}_j$ by $\boldsymbol{C} = \boldsymbol{M}\boldsymbol{w}$ in the CpG-level derivation leaves stage 1, and hence $\boldsymbol{\zeta}$, unchanged. Because eq. (320) is linear in $\boldsymbol{Z}$, the score for adding the clock to the stage-1 model is exactly the $\boldsymbol{w}$-weighted sum of the CpG-level scores:

$$U_\Theta(\hat{\eta}, 0) = \boldsymbol{C}^\top \boldsymbol{\zeta} = \widetilde{\boldsymbol{C}}^\top \boldsymbol{\zeta} = \sum_{j=1}^{J} w_j\, U_{\vartheta_j}(0) \tag{345}$$

Applying assumptions 1–3 and non-informative censoring to the clock (with assumption 2 now meaning that $\mathrm{Var}(C_i \mid X_i)$ does not depend on $X_i$), the information is approximated by $N\pi\,\mathrm{Var}(\widetilde{C})$ as in eq. (334). The one-step estimator of $\Theta_{\mathrm{Cox}}$ in the individual-level model eq. (242) is therefore

$$\widehat{\Theta}^{(1)}_{\mathrm{Cox}} \approx \frac{\widetilde{\boldsymbol{C}}^\top \boldsymbol{\zeta}}{N\pi\,\mathrm{Var}(\widetilde{C})} = \frac{\mathrm{Cov}(\boldsymbol{\zeta}, \widetilde{\boldsymbol{C}})}{\pi\ \mathrm{Var}(\widetilde{C})} \tag{346}$$

Comparing eq. (346) with eq. (344), we have

$$\boxed{\hat{\gamma} \approx \pi\, \widehat{\Theta}^{(1)}_{\mathrm{Cox}}} \tag{347}$$

Thus, SARDINE's output `gamma` is a scaled version of the the log hazard ratio (this is the clock-level analogue of $b_j \approx \pi\, \hat{\vartheta}_j^{(1)}$ (eq. (336))). The clock-level log hazard ratio and its standard error then follow from the SARDINE output by dividing by the event fraction:

$$\widehat{\Theta}_{\text{Cox}} = \hat{\gamma}/\pi \tag{348}$$

$$\text{SE}\big(\widehat{\Theta}_{\text{Cox}}\big) = \text{SE}(\hat{\gamma})/\pi \tag{349}$$

The rescaling leaves the $Z$ statistic and $P$ value of $\hat{\gamma}$ unchanged. SARDINE's $Z$ statistic approximates the score test of $H_0 : \Theta_{\text{Cox}} = 0$ in the individual-level Cox model.

When $\pi$ is not reported, by eq. (338), it can be estimated as $\widehat{\text{Var}}(\boldsymbol{\zeta})$ which SARDINE can estimate (eq. (28)) using EWAS summary statistics and an external reference. As a diagnostic (for the mismatch between EWAS and reference panel), the ratio $\widehat{\text{Var}}(\boldsymbol{\zeta})/\pi$ should be close to one.

By eqs. (343) and (344), $\widehat{\lambda} = \hat{\gamma}\, \text{Var}(\widetilde{C})/\,\text{Var}(\boldsymbol{\zeta})$, so with $\text{Var}(\boldsymbol{\zeta}) \approx \pi$,

$$\widehat{\lambda} \approx \sigma^2_{\widetilde{C}}\, \widehat{\Theta}^{(1)}_{\text{Cox}} \tag{350}$$

Hence $\widehat{\lambda}$ is a rescaled version of $\hat{\gamma}$. It is the slope in the regression of residualized clock on the martingale residual, not the slope of the residualized clock on residualized case status. To see the difference, note that $P\boldsymbol{\zeta} = \boldsymbol{\zeta}$ gives $\boldsymbol{\zeta} = P\boldsymbol{\delta} - P\hat{\boldsymbol{\Lambda}} = \widetilde{\boldsymbol{\delta}} - \widetilde{\boldsymbol{\Lambda}}$, so that

$$\text{Cov}(\boldsymbol{\zeta}, \widetilde{\boldsymbol{C}}) = \text{Cov}(\widetilde{\boldsymbol{\delta}}, \widetilde{\boldsymbol{C}}) - \text{Cov}(\widetilde{\boldsymbol{\Lambda}}, \widetilde{\boldsymbol{C}}) \tag{351}$$

where the second term does not vanish because $\hat{\Lambda}_i = \widehat{H}_0(T_i)\, e^{X_i \hat{\eta}}$ depends on the follow-up time $T_i$, which depends on $C_i$ whenever $\Theta_{\text{Cox}} \neq 0$: for $\Theta_{\text{Cox}} > 0$, individuals with higher clock values tend to fail earlier, so their follow-up, and hence $\hat{\Lambda}_i$, is shorter on average. The martingale residual therefore records when events occurred as well as whether they occurred.

### 4 Appendix

#### 4.1 Simulation studies

In the simulation, our goal is to simulate a phenotype $Y$ to achieve a given association (i.e., $\lambda$ and $\gamma$) with the epigenetic clock $C$.

**Continuous phenotype.** To simulate a continuous trait, we first standardize $C$ to mean 0 and variance 1, denoted as $C_{\text{std}}$; and generate $Y$ as a linear combination of the clock and an independent random noise $\epsilon$:

$$Y = rC_{\text{std}} + \epsilon \tag{352}$$

where $\epsilon \sim N(0, 1 - r^2)$ is independent of $C_{\text{std}}$. Because $\text{Var}(C_{\text{std}}) = 1$, this data-generating process ensures that $Y$ has mean 0 and variance 1; and that $\text{Cov}(Y, C_{\text{std}}) = r\,\text{Var}(C_{\text{std}}) = r$. The population OLS slopes are

$$\gamma = \frac{\text{Cov}(Y, C_{\text{std}})}{\text{Var}(C_{\text{std}})} = r$$
$$\lambda = \frac{\text{Cov}(Y, C_{\text{std}})}{\text{Var}(Y)} = r$$

Therefore, simulating the continuous phenotype using eq. (352) guarantees that the phenotype–clock regression coefficients converge to $r$.

**Binary phenotype.** To simulate binary outcomes, we used a continuous liability threshold model, where an underlying liability was first generated using eq. (352) and subsequently dichotomized at a threshold corresponding to a prevalence $K$.

Let $L = rC_{\text{std}} + \epsilon$ denote the continuous liability. For the derivations below, we assume $C_{\text{std}} \sim N(0, 1)$ so that $L \sim N(0, 1)$ and $(C_{\text{std}}, L)$ follows are standard bivariate normal distribution with correlation $r$. The observed binary phenotype $Y \in \{0, 1\}$ is determined by whether $L$ exceeds the threshold $\tau$:

$$Y = \begin{cases} 1 & \text{if } L > \tau \\ 0 & \text{if } L \leq \tau \end{cases}$$

where $\tau = \Phi^{-1}(1 - K)$ and $\Phi(\cdot)$ is the cumulative distribution function of the standard normal distribution. We define $\phi(\tau)$ as the standard normal probability density function evaluated at this threshold.

Next, we establish the relationship between $r$ (the correlation between the liability and the clock) and the parameters of interests $\lambda$ and $\gamma$ (which quantify the associations between binary phenotype $Y$ and clock). In the simulation, we can easily first fix the values of $\lambda$ and $\gamma$, then compute the corresponding $r$, and then simulate $L$ and binary $Y$.

**Case-control mean difference ($\lambda$).** The parameter $\lambda$ from the regression model $C_{\text{std}} \sim Y$ represents the mean difference in the standardized clock between cases and controls:

$$\lambda = \mathbb{E}[C_{\text{std}} \mid Y = 1] - \mathbb{E}[C_{\text{std}} \mid Y = 0]$$

Because $C_{\text{std}}$ and $L$ are standard bivariate normal, $\mathbb{E}[C_{\text{std}} \mid L] = rL$. Because $Y$ is a function of $L$, the law of iterated expectations allows us to rewrite the conditional expectations of $C_{\text{std}}$ in terms of the truncated distribution of $L$:

$$\mathbb{E}[C_{\text{std}} \mid Y = 1] = r\mathbb{E}[L \mid L > \tau] = r\frac{\phi(\tau)}{K}$$
$$\mathbb{E}[C_{\text{std}} \mid Y = 0] = r\mathbb{E}[L \mid L \leq \tau] = r\frac{-\phi(\tau)}{1-K}$$

Substituting these into the equation for $\lambda$ yields:

$$\lambda = r\left(\frac{\phi(\tau)}{K} - \frac{-\phi(\tau)}{1-K}\right) = \frac{r\phi(\tau)}{K(1-K)}$$

To simulate data with a predefined mean difference $\lambda$, we reverse-engineer the required liability-scale correlation $r$:

$$r = \frac{\lambda K(1-K)}{\phi(\tau)} \tag{353}$$

**Log-odds ratio ($\gamma$).** Alternatively, one may want to run a logistic regression of disease status on the epigenetic clock ($Y \sim C_{\text{std}}$). Since the liability $L$ and clock $C_{\text{std}}$ are bivariate normal with correlation $r$, conditional on $C_{\text{std}}$ the liability follows a normal distribution:

$$L \mid C_{\text{std}} \sim N(rC_{\text{std}}, 1 - r^2)$$

The probability of being a case given the clock value is the probability that the liability $L$ exceeds the threshold $\tau$:

$$\begin{aligned}
\mathrm{P}(Y = 1 \mid C_{\text{std}}) &= \mathrm{P}(rC_{\text{std}} + \epsilon > \tau) \\
&= \Phi\left(\frac{rC_{\text{std}} - \tau}{\sqrt{1-r^2}}\right) \\
&= \Phi\left(-\frac{\tau}{\sqrt{1-r^2}} + \frac{rC_{\text{std}}}{\sqrt{1-r^2}}\right) \qquad (354)
\end{aligned}$$

By definition, a probit regression model models the conditional probability of a binary outcome $Y$ as:

$$P(Y = 1 \mid X) = \Phi(\alpha + \beta_{\text{probit}} X)$$

Comparing the two equations above, we have

$$\beta_{\text{probit}} = \frac{r}{\sqrt{1 - r^2}}$$

In empirical applications, binary traits are commonly analyzed using logistic regressions:

$$P(Y = 1 \mid C_{\text{std}}) = \frac{\exp(\alpha_{\text{logit}} + \gamma C_{\text{std}})}{1 + \exp(\alpha_{\text{logit}} + \gamma C_{\text{std}})} = \text{logit}^{-1}(\alpha_{\text{logit}} + \gamma C_{\text{std}})$$

where $\text{logit}^{-1}(x) = 1/(1 + e^{-x})$ is the CDF of the standard logistic distribution.

Since the standard logistic distribution used in logistic regression has a variance of $\pi^2/3$ whereas the standard normal distribution used in probit regression has a variance of 1, matching the two variances gives us a commonly used approximate relationship between the probit coefficient and the logistic log-odds ratio:

$$\gamma \approx \beta_{\text{probit}} \frac{\pi}{\sqrt{3}} = \frac{r}{\sqrt{1 - r^2}} \frac{\pi}{\sqrt{3}}$$

Solving for $r$ to achieve a target logistic coefficient $\gamma$:

$$r \approx \frac{\gamma\sqrt{3}/\pi}{\sqrt{1 + \left(\gamma\sqrt{3}/\pi\right)^2}} \tag{355}$$

However, despite being a closed-form solution, the approximation above can be very inaccurate, because the logistic model does not match the data-generating model.

Under the liability threshold data-generating model, the true case probability is the probit function eq. (354)

$$p(C_{\text{std}}) \equiv P(Y = 1 \mid C_{\text{std}}) = \Phi\left(\frac{rC_{\text{std}} - \tau}{\sqrt{1 - r^2}}\right)$$

which cannot be written as $\text{logit}^{-1}(\alpha_{\text{logit}} + \gamma C_{\text{std}})$ for any $(\alpha_{\text{logit}}, \gamma)$. The logistic model is therefore misspecified, and we first need to define the quantity that logistic regression estimates.

Let $\eta = \alpha_{\text{logit}} + \gamma C_{\text{std}}$. For one individual, the logistic log-likelihood is

$$\begin{aligned}\ell(\alpha_{\text{logit}}, \gamma) &= Y \log\left\{\text{logit}^{-1}(\eta)\right\} + (1-Y)\log\left\{1 - \text{logit}^{-1}(\eta)\right\} \\ &= Y(\eta - \log(1+e^{\eta})) + (1-Y)(-\log(1+e^{\eta}) \\ &= Y\eta - \log(1+e^{\eta})\end{aligned}$$

The logistic regression estimator maximizes the sample average of $\ell(\alpha_{\text{logit}}, \gamma)$. Even when the model is misspecified, this estimator converges almost surely to the parameter value that minimizes the Kullback–Leibler information criterion (KLIC) between the true distribution and the model [15, Theorem 2.2]. Here, the KLIC is

$$\text{KLIC}(\alpha_{\text{logit}}, \gamma) = \mathbb{E}\left[\log \text{P}(Y \mid C_{\text{std}})\right] - \mathbb{E}\left[\ell(\alpha_{\text{logit}}, \gamma)\right]$$

where $\text{P}(Y \mid C_{\text{std}})$ is the true conditional probability of the observed $Y$. Because the first term does not depend on $(\alpha_{\text{logit}}, \gamma)$, minimizing the KLIC is equivalent to maximizing the expected log-likelihood:

$$(\alpha^*_{\text{logit}}, \gamma^*) = \underset{\alpha_{\text{logit}}, \gamma}{\arg\max}\ \mathbb{E}\left[\ell(\alpha_{\text{logit}}, \gamma)\right]$$

where the expectation is taken under the true data-generating model. We define $\gamma^*$ as the true logistic log-odds ratio.

Because $\eta$ depends on the data only through $C_{\text{std}}$, the law of iterated expectations gives

$$\begin{aligned}\mathbb{E}\left[\ell(\alpha_{\text{logit}}, \gamma)\right] &= \mathbb{E}\left[\mathbb{E}(Y \mid C_{\text{std}})\,\eta - \log(1+e^{\eta})\right] \\ &= \mathbb{E}\left[p(C_{\text{std}})\,\eta - \log(1+e^{\eta})\right]\end{aligned}$$

where the outer expectation is taken over $C_{\text{std}} \sim N(0,1)$. Setting the derivatives with respect to $\alpha_{\text{logit}}$ and $\gamma$ to zero yields

$$\mathbb{E}\left[p(C_{\text{std}}) - \text{logit}^{-1}(\alpha_{\text{logit}} + \gamma C_{\text{std}})\right] = 0 \qquad (356)$$

$$\mathbb{E}\left[C_{\text{std}}\left\{p(C_{\text{std}}) - \text{logit}^{-1}(\alpha_{\text{logit}} + \gamma C_{\text{std}})\right\}\right] = 0 \qquad (357)$$

The solution of these two equations is $(\alpha^*_{\text{logit}}, \gamma^*)$. However, they do not have closed-form solutions. We therefore evaluated them by numerical integration and solved the two equations numerically, yielding $\gamma^*(r)$ for any given $r$. For a target log-odds ratio $\gamma$, the required $r$ is the root of

$$\gamma^*(r) = \gamma \qquad (358)$$

which we obtained by univariate root-finding over $r \in (0, 1)$.

**Linear-probability slope** ($\gamma_{\text{lin}}$). For a linear regression of the binary phenotype on the clock ($Y \sim C_{\text{std}}$), the OLS linear-probability slope is

$$\begin{aligned}\gamma_{\text{lin}} &= \frac{\text{Cov}(Y, C_{\text{std}})}{\text{Var}(C_{\text{std}})} = \mathbb{E}[C_{\text{std}} Y] \\ &= K\,\mathbb{E}[C_{\text{std}} \mid Y = 1] \\ &= r\phi(\tau)\end{aligned}$$

so that $r = \gamma_{\text{lin}}/\phi(\tau)$. Because $r \leq 1$, only $\gamma_{\text{lin}} \leq \phi(\tau)$ (`dnorm(qnorm(K, lower.tail = F))` $\approx 0.348$ for $K = 0.3$) are attainable.
