## Supplementary Figures for "SARDINE unlocks EWAS summary statistics for large-scale epigenetic biomarker analysis"

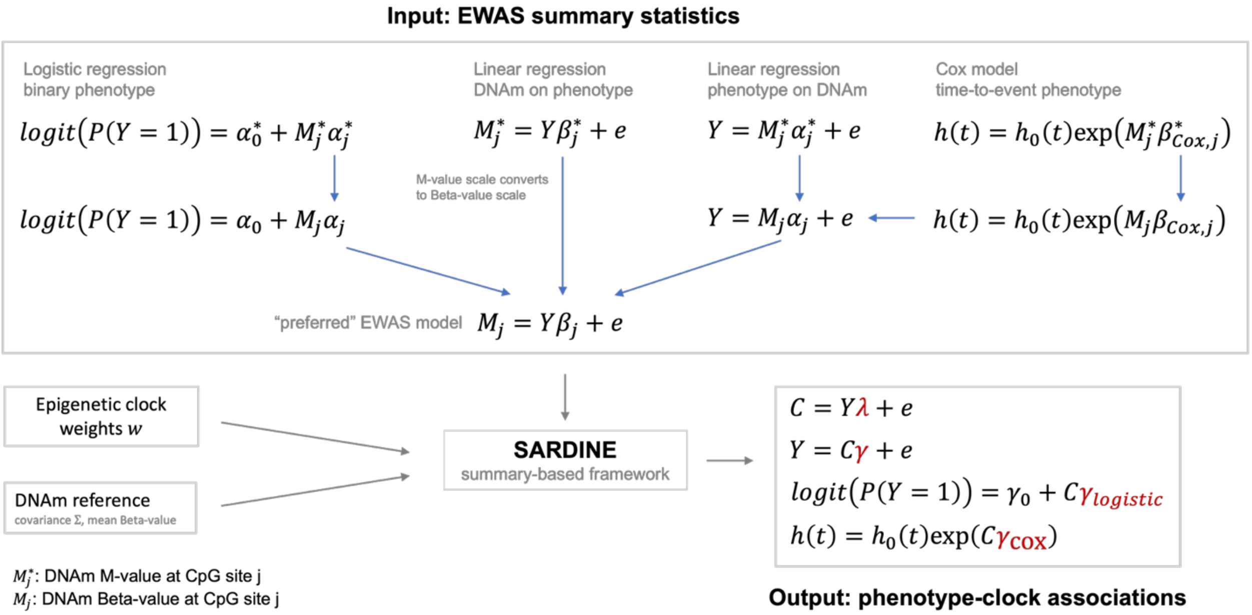


**Supplementary Figure 1. Method workflow.** SARDINE estimates the association between a phenotype Y and an epigenetic clock C without requiring individual-level data. The inputs are EWAS summary statistics, CpG weights for the epigenetic clock, and a DNAm variance–covariance matrix estimated from a reference panel. Summary statistics may be obtained from any of four study designs: linear regression of methylation M on Y, linear regression of Y on M, logistic regression of Y on M, or a Cox proportional hazards model. We allow methylation to be quantified on either the Beta-value or M-value scale. SARDINE converts each of these to a common scale internally (**Methods**) and returns the effects of the clock (phenotype) on phenotype (clock) $\gamma$ ($\lambda$) with a standard error and P value. When the phenotype is binary, SARDINE additionally estimates the log odds ratio of Y per unit increase in C. When the input summary statistics is from a Cox proportional hazards model, SARDINE additionally estimates the log hazard ratio per unit increase in C.

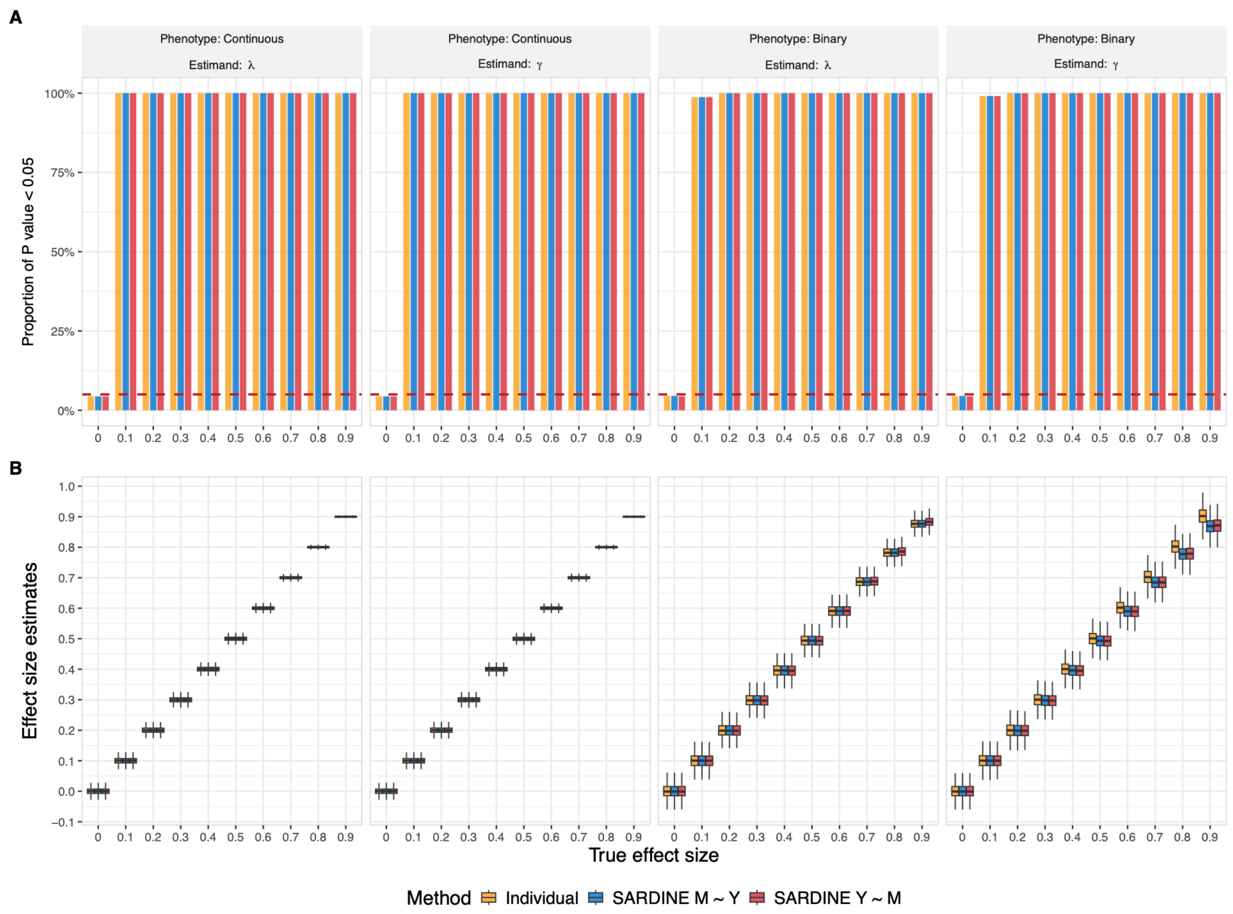


**Supplementary Figure 2. Full simulation results based on in-sample DNAm reference.** These simulations were based on identical settings presented in **Figure 2** except that here we show a broader range of parameter values. When the true effect sizes are very large (which is rare in practice), we observed a slight downward bias in log odds ratio estimates. No bias was observed for continuous traits.

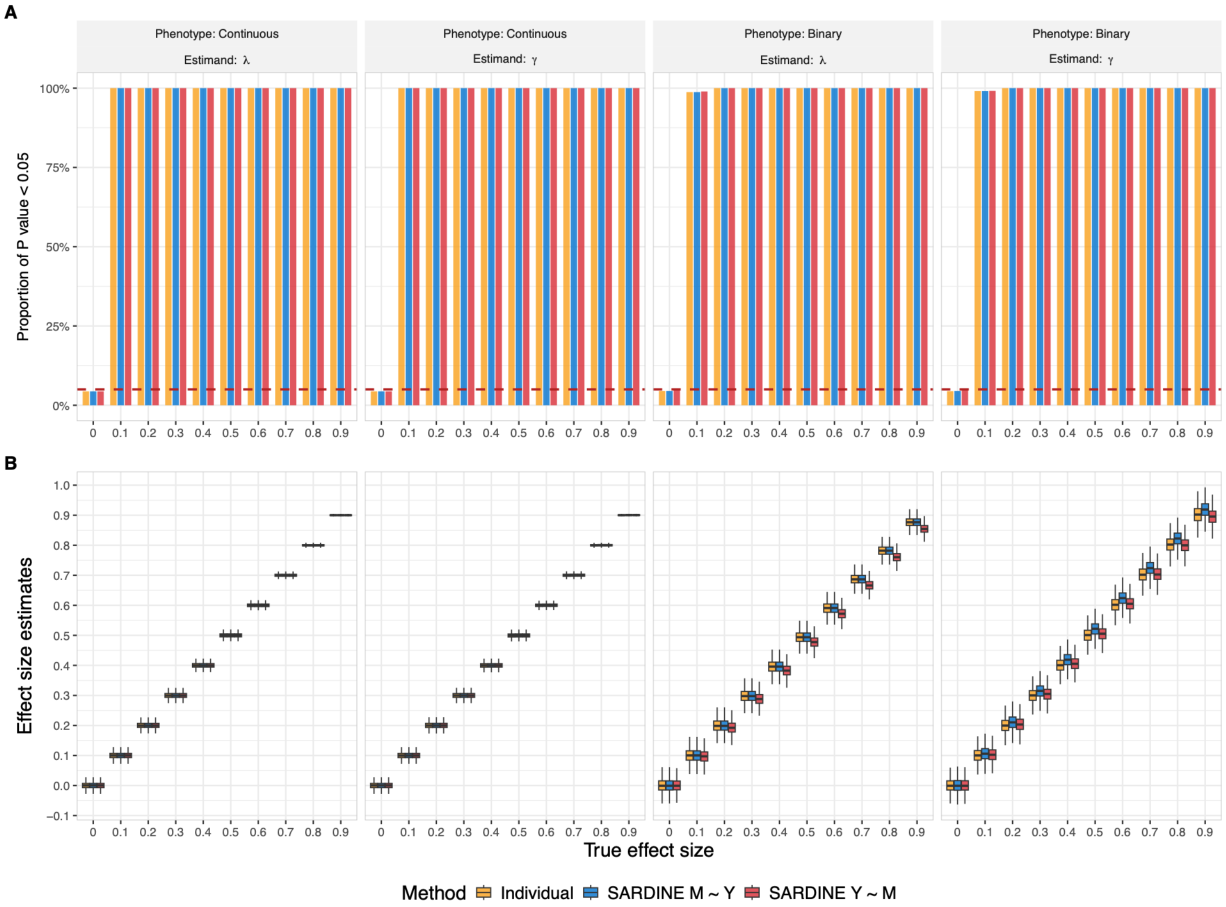


**Supplementary Figure 3. Simulation results based on an external DNAm reference derived from the holdout TruDiagnostic European ancestry samples with N = 500.** These simulations were based on identical settings presented in **Supplementary Figure 2**, except that an external reference dataset with N=500 was used to estimate DNAm covariance in SARDINE.

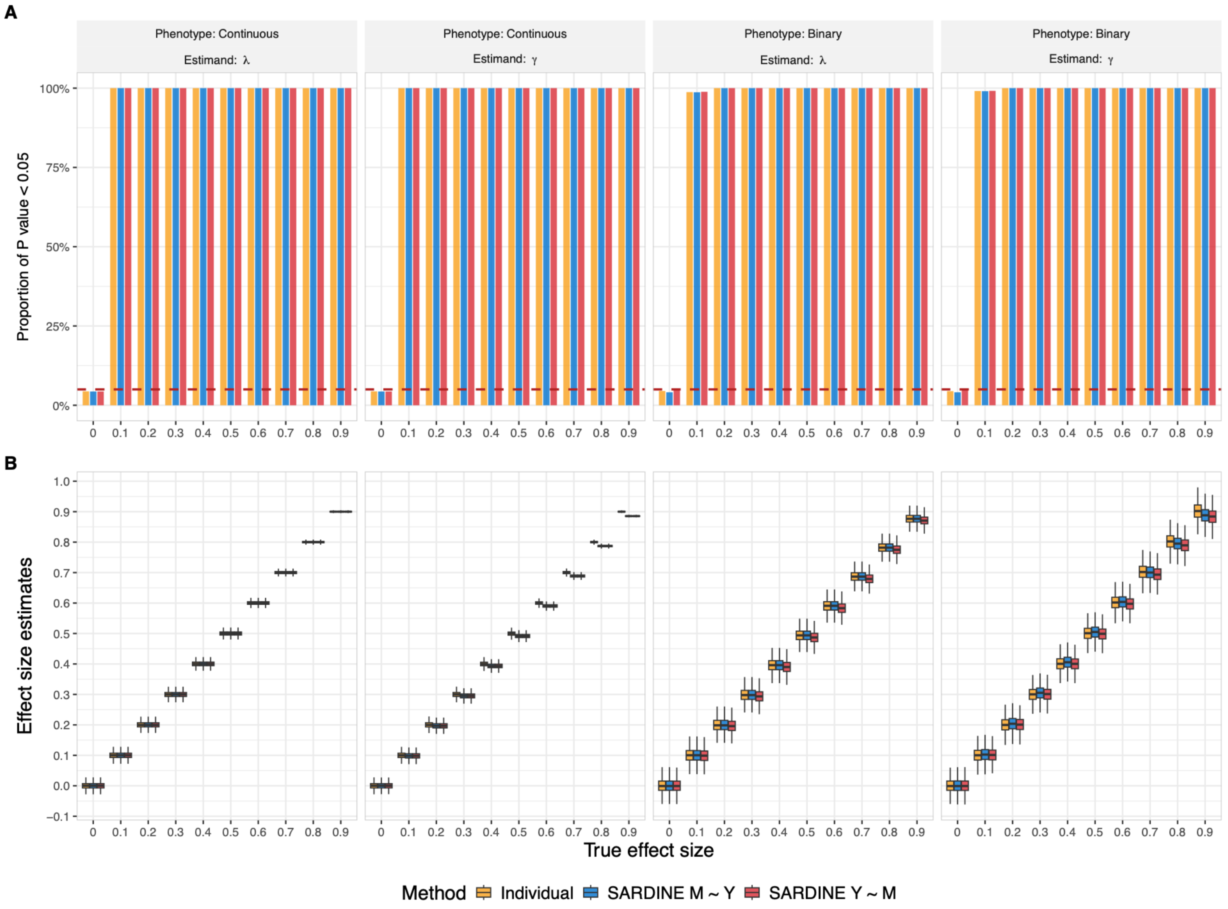


**Supplementary Figure 4. Simulation results based on an external DNAm reference derived from the holdout TruDiagnostic European ancestry samples with N = 2,000.** These simulations were based on identical settings presented in **Supplementary Figure 2**, except that an external reference dataset with N=2,000 was used to estimate DNAm covariance in SARDINE.

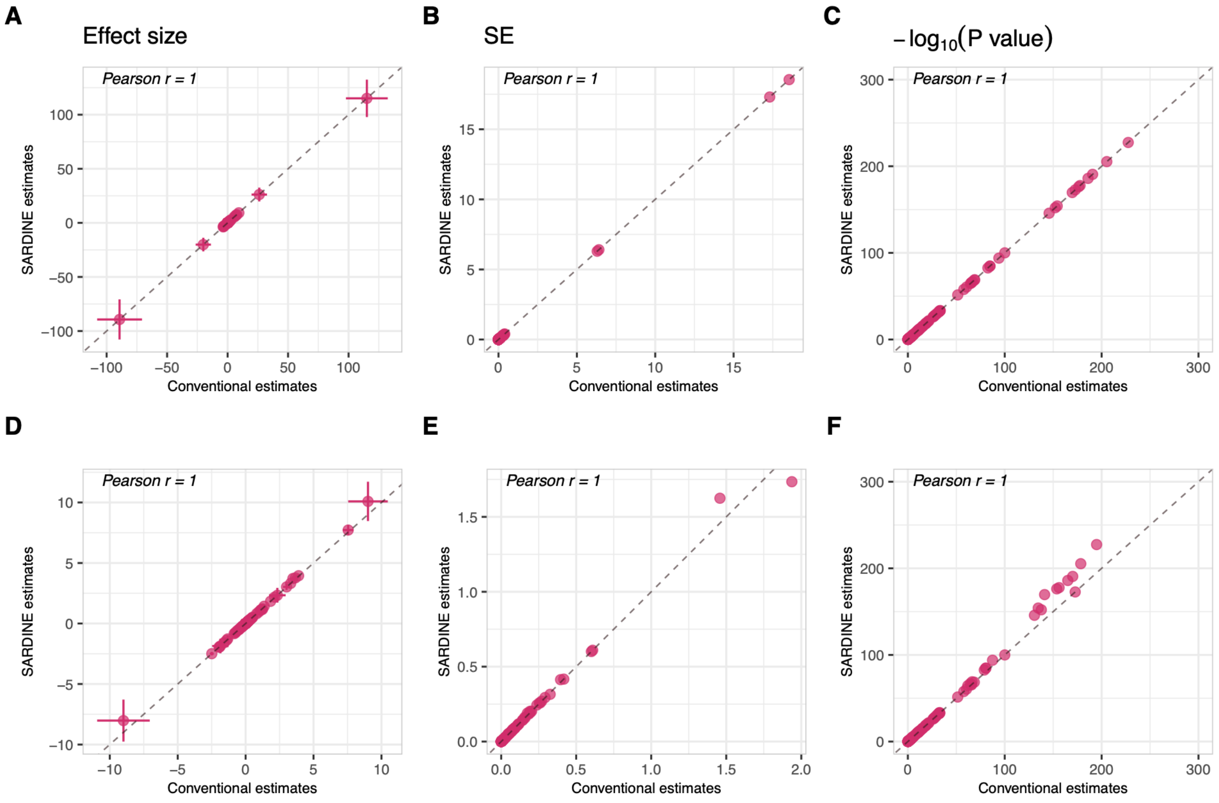


**Supplementary Figure 5. Validating SARDINE performance using four EWAS performed in TruDiagnostic samples.** (A) Estimates of $\lambda$. This is the same information presented in **Figure 3A** except that the outlier estimates with extreme values are also shown. (B) Standard errors for $\lambda$ estimates. (C) -log10(P-value) for $\lambda$ estimates. (D) Estimates of $\gamma$. This is the same information presented in **Figure 3B** except that the outlier estimates with extreme values are also shown. (E) Standard errors for $\gamma$ estimates. (F) -log10(P-value) for $\gamma$ estimates. No covariate was adjustment in these analyses. Each data point in the plot is a clock-phenotype combination, and the dashed line is y = x.

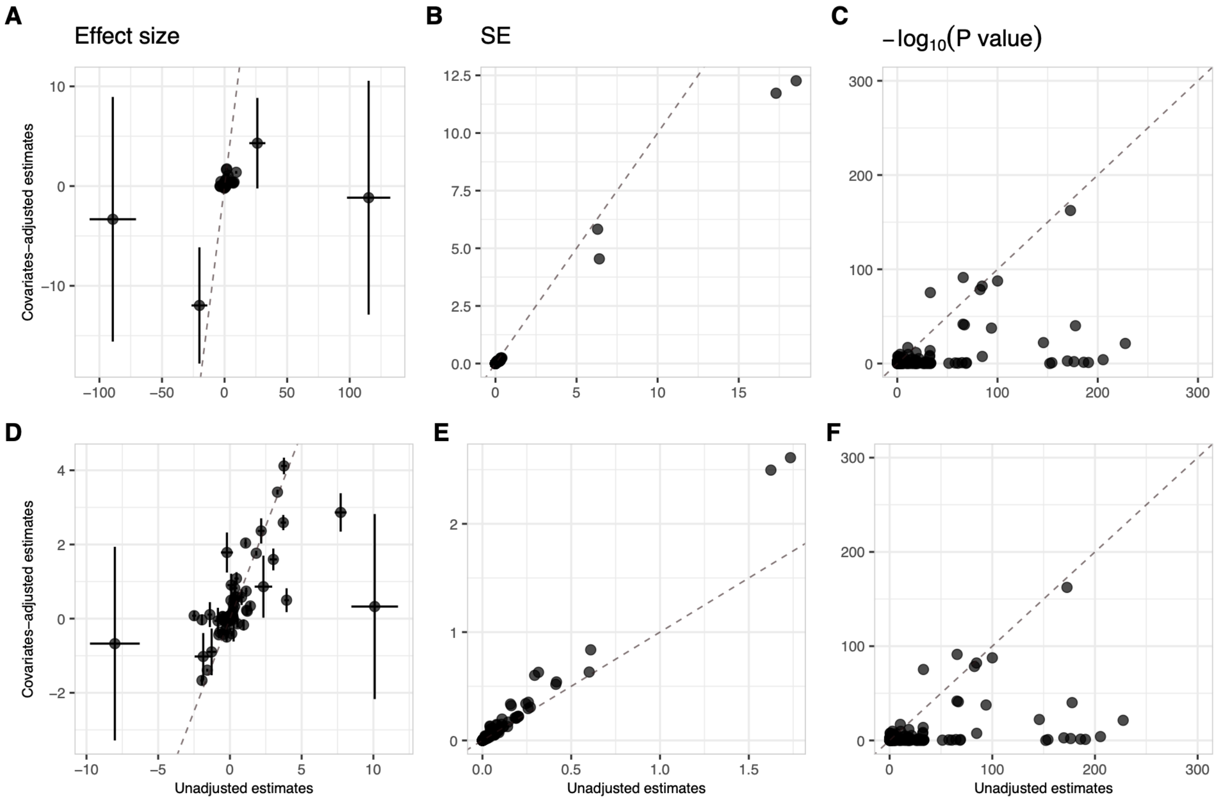


**Supplementary Figure 6. Comparing epigenetic clock-phenotype association estimates with and without covariates adjustment.** In each panel, the x-axis shows SARDINE estimates without adjusting for covariates, while the y-axis shows association estimates after adjusting for age, $age^2$, sex, and cell type composition. In all panels, each data point is a clock-phenotype pair, and the dashed line is y = x. (A) Estimates of $\lambda$. Intervals show standard errors. (B) Standard errors for $\lambda$ estimates. (C) -log10(P-value) for $\lambda$ estimates. (D) Estimates of $\gamma$. Intervals show standard errors. (E) Standard errors for $\gamma$ estimates. (F) -log10(P-value) for $\gamma$ estimates.

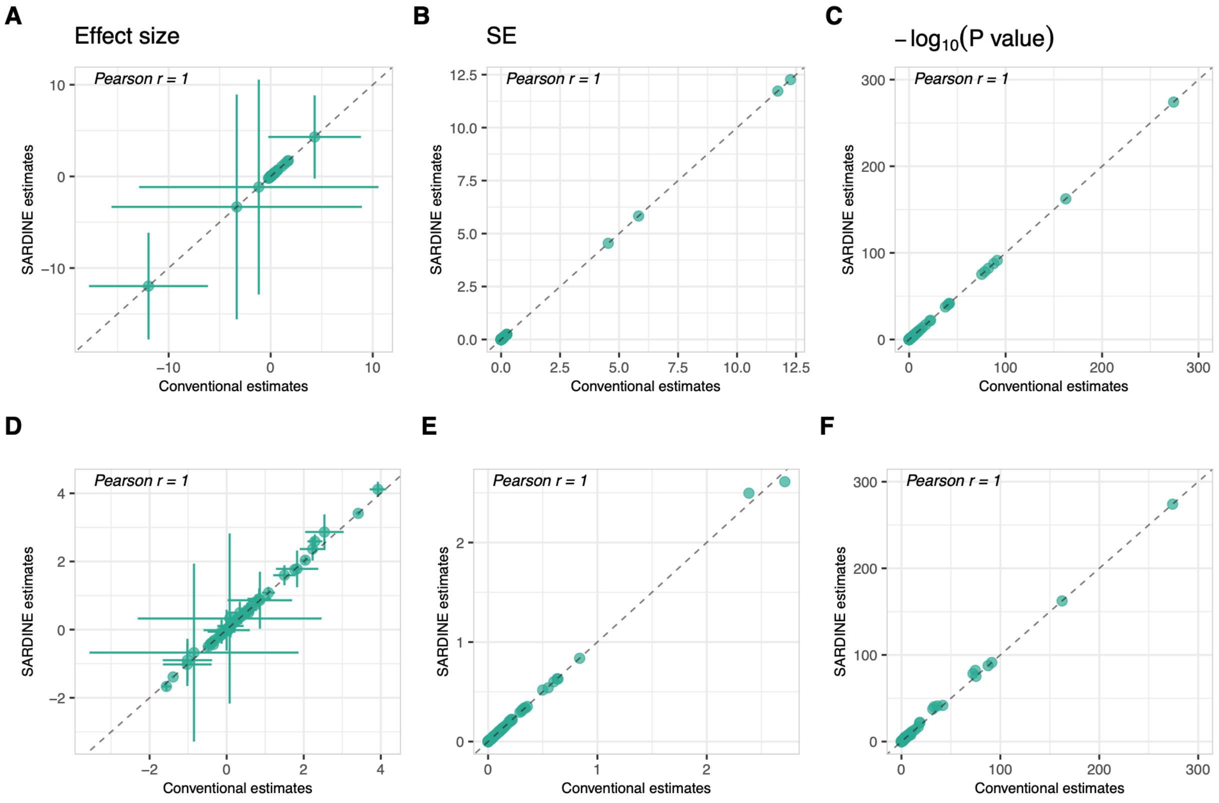


**Supplementary Figure 7. Validating covariates-adjusted SARDINE estimates using four EWAS performed in TruDiagnostic samples.** (A) Estimates of $\lambda$. This is the same information presented in **Figure 3C**. (B) Standard errors for $\lambda$ estimates. (C) -log10(P-value) for $\lambda$ estimates. (D) Estimates of $\gamma$. This is the same information presented in **Figure 3D**. (E) Standard errors for $\gamma$ estimates. (F) -log10(P-value) for $\gamma$ estimates. Sex, age, age$^2$, and cell type composition were adjusted for in these analyses. Each data point in the plot is a clock-phenotype combination, and the dashed line is y = x.

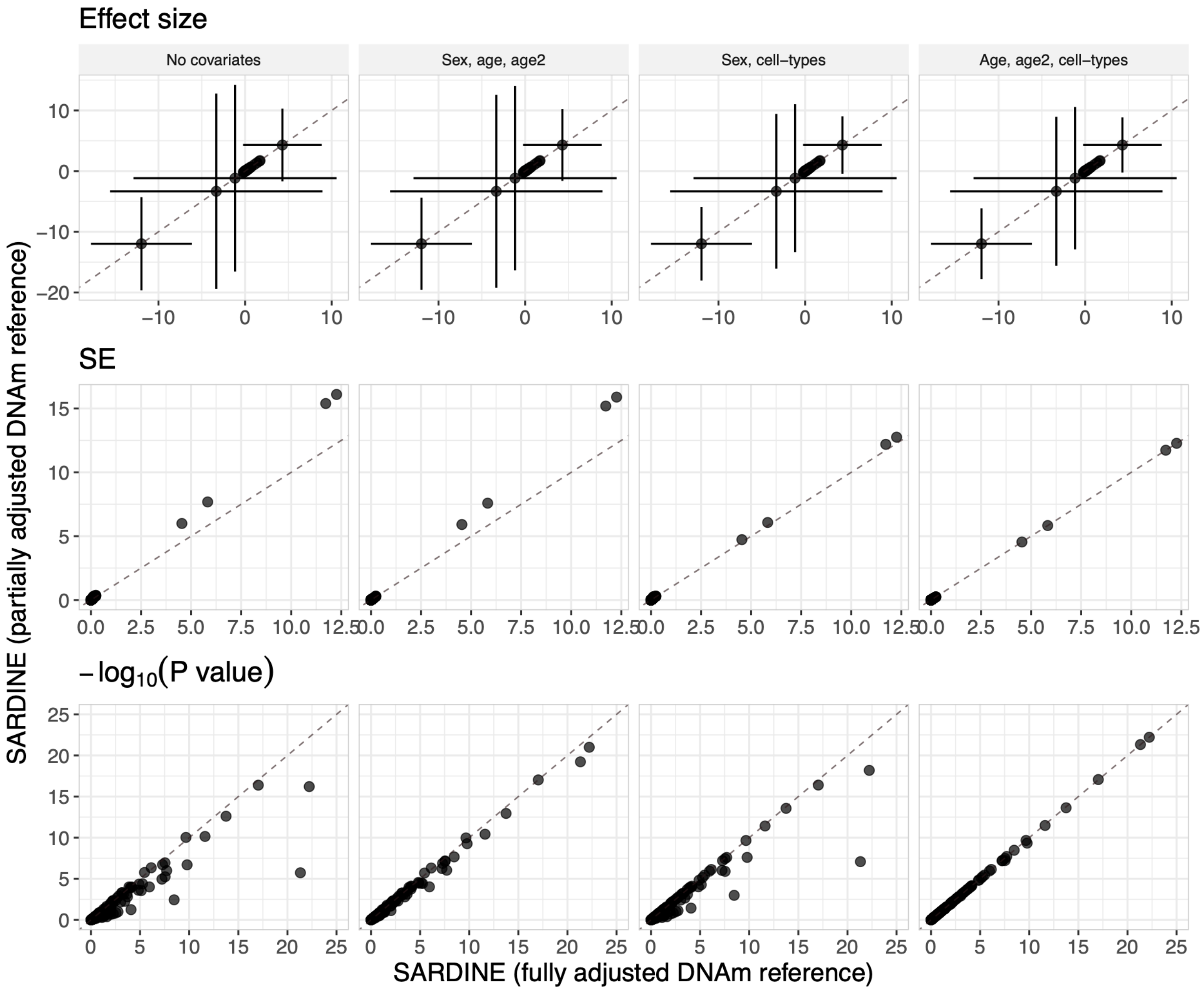


**Supplementary Figure 8. Effect of covariate adjustment in DNAm in-sample reference matrix on SARDINE's estimate for $\lambda$.** EWAS were conducted for 4 phenotypes (height, BMI, anxiety, and high blood pressure) in the full TruDiagnostic sample, adjusting for the full set of covariates (sex, age, age2, and cell-type compositions). The x-axis shows SARDINE results obtained with a matched (with EWAS) DNAm reference (i.e. adjusted for the same covariates: sex, age, $age^2$, and 12 cell type compositions), and the y-axis show SARDINE results obtained with a partially adjusted DNAm reference. Each panel is labeled by the covariates adjusted for in the DNAm reference. Points with $-log_{10}$(*P*-value) greater than 25 are omitted for better visualization. Each data point in the plot is a clock-phenotype combination, and the dashed line is y = x.

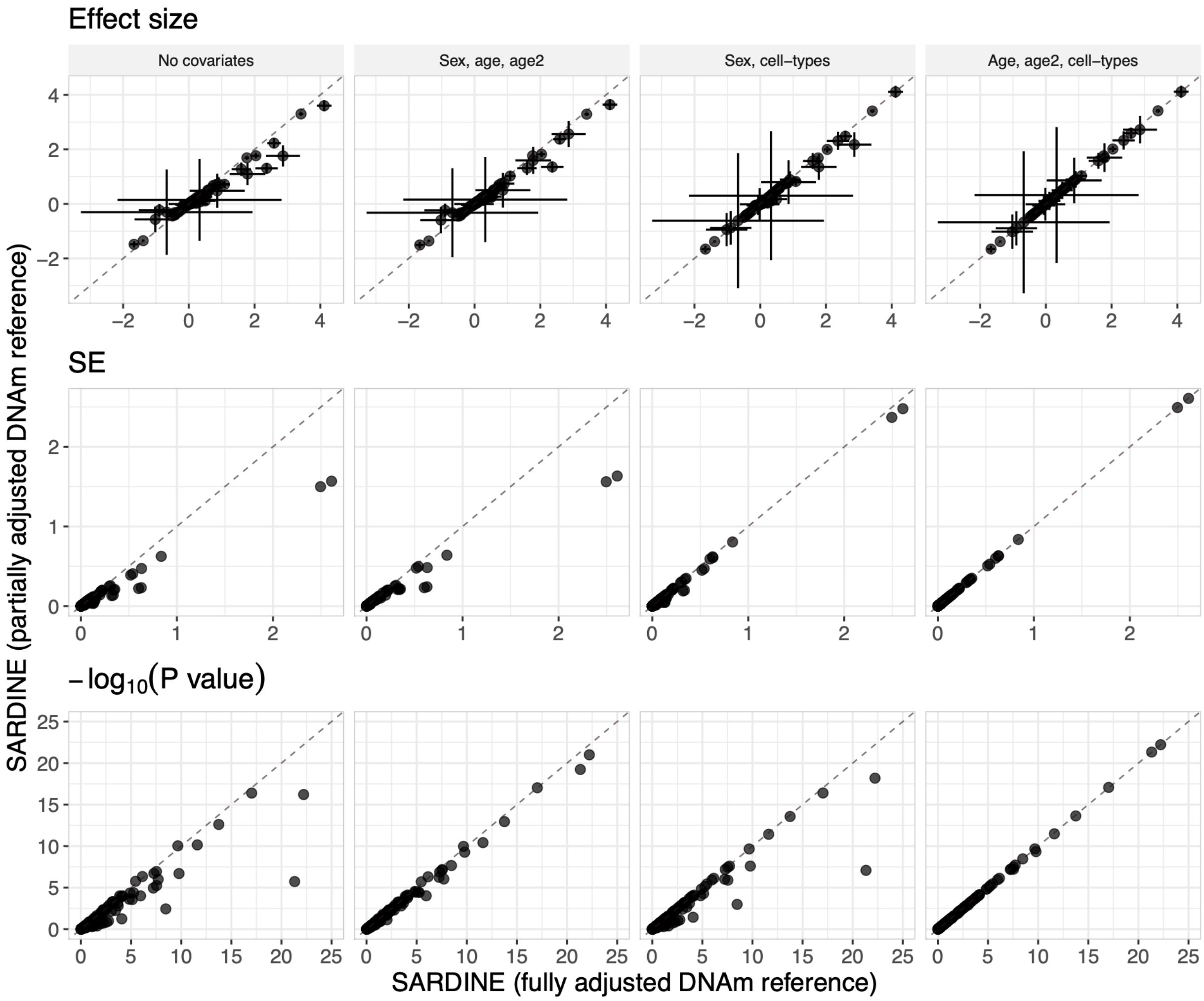


**Supplementary Figure 9. Effect of covariate adjustment in DNAm reference matrix on SARDINE's estimate for $\gamma$.** All settings are identical with those in **Supplementary Figure 8**, except in this plot we show the results for $\gamma$.

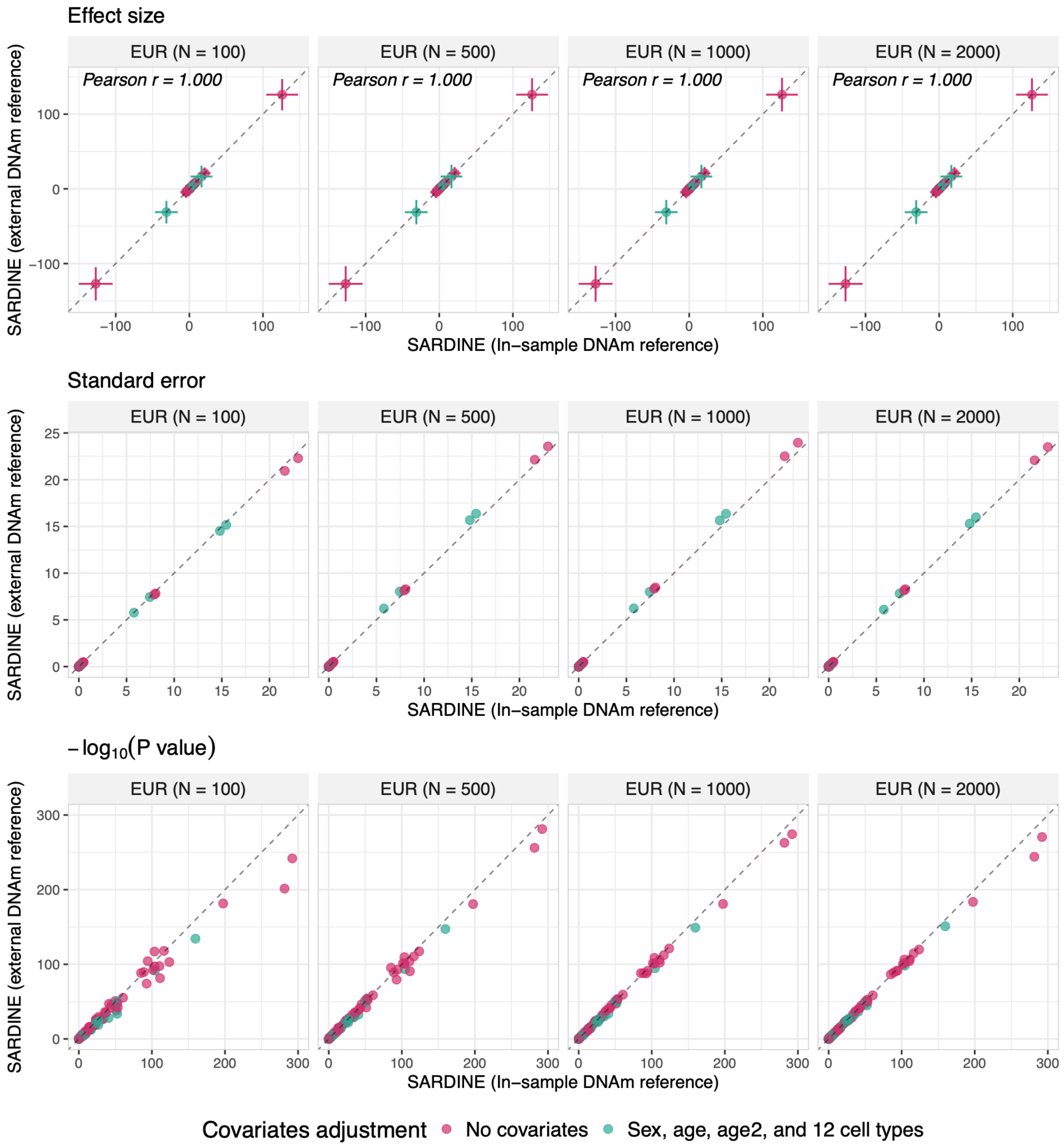


**Supplementary Figure 10. Sensitivity of SARDINE's estimates for $\lambda$ to the external DNAm reference panel.** EWAS were conducted for 4 phenotypes (height, BMI, anxiety, and high blood pressure) in a subset of TruDiagnostic cohort (N = 8,880 individuals of European ancestry). External DNAm references were computed using independent European ancestry samples from TruDiagnostic with different sample sizes as shown by each panel's label. Identical covariate adjustment was applied to the EWAS and every DNAm reference. Each data point in the plot is a clock-phenotype combination, and the dashed line is y = x.

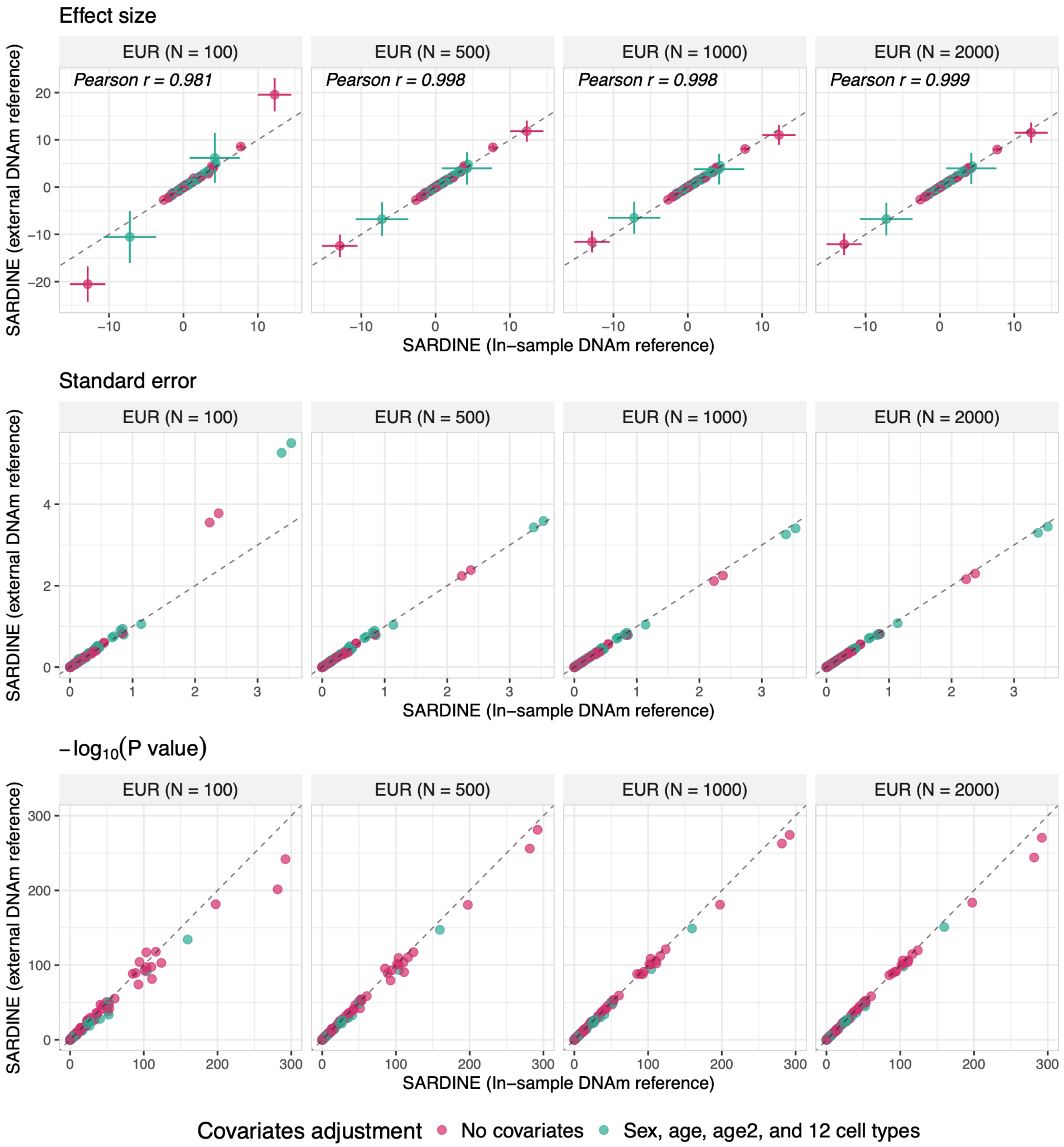


**Supplementary Figure 11. Sensitivity of SARDINE's estimates for $\gamma$ to the external DNAm reference panel.** All settings are identical with those in **Supplementary Figure 10**, except in this plot we show the results for $\gamma$.

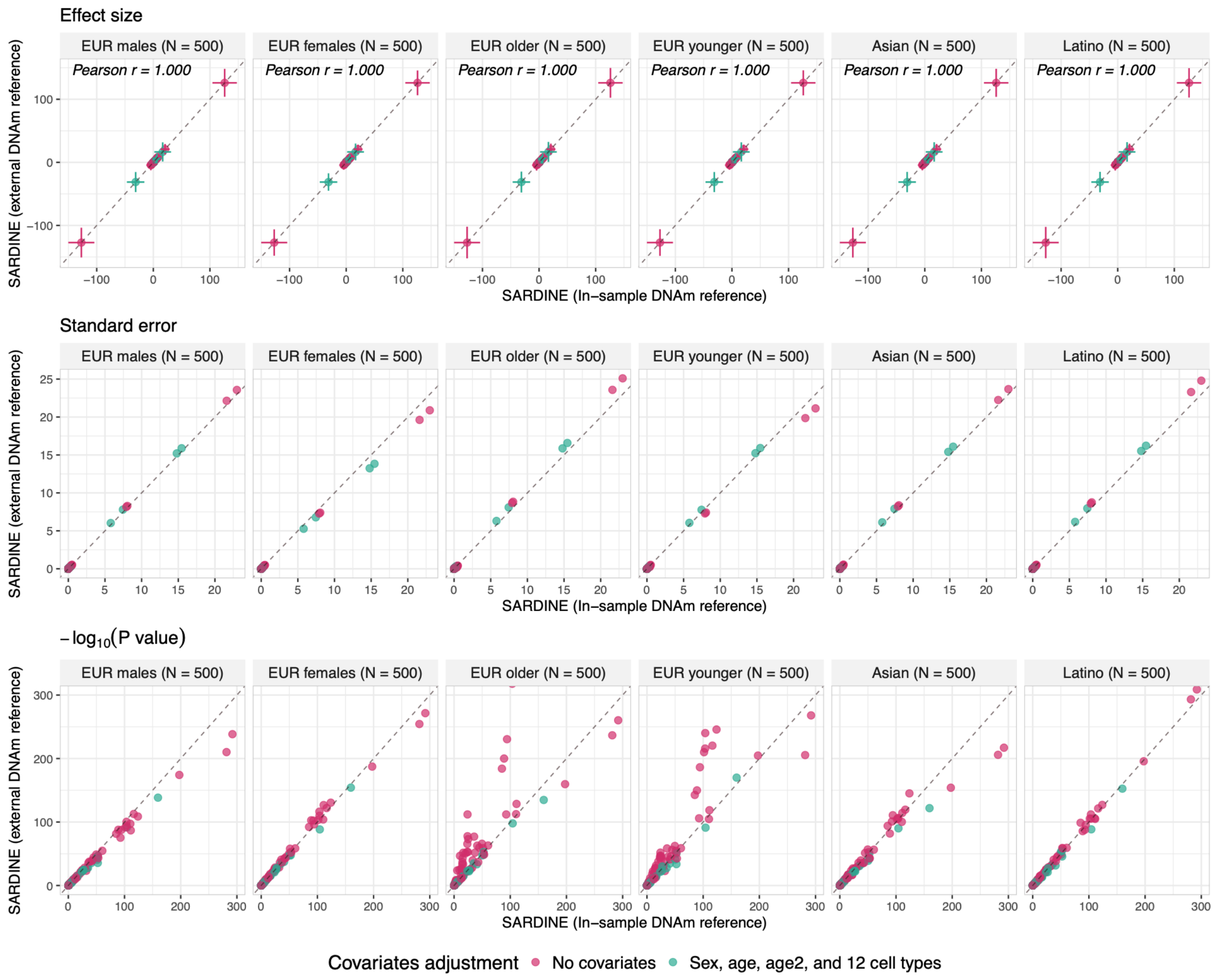


**Supplementary Figure 12. Sensitivity of SARDINE's estimates for $\lambda$ to the external DNAm reference panel.** All settings are identical with those in **Supplementary Figure 10**, except in this plot the external DNAm references were computed using independent TruDiagnostic samples with diverse demographics as shown by each panel's label.

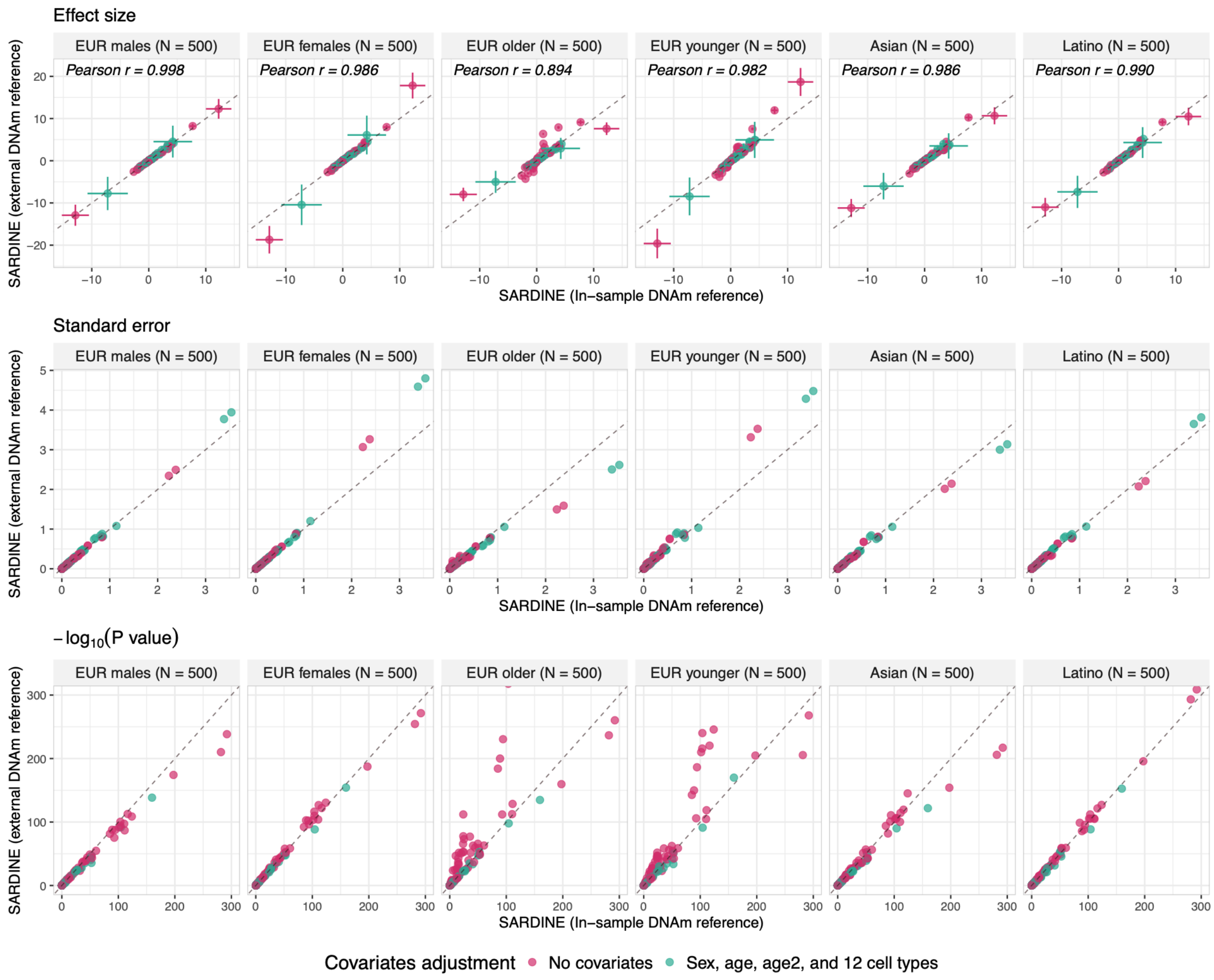


**Supplementary Figure 13. Sensitivity of SARDINE's estimates for $\gamma$ to the external DNAm reference panel.** All settings are identical with those in the **Supplementary Figure 12**, except in this plot we show the results for $\gamma$.

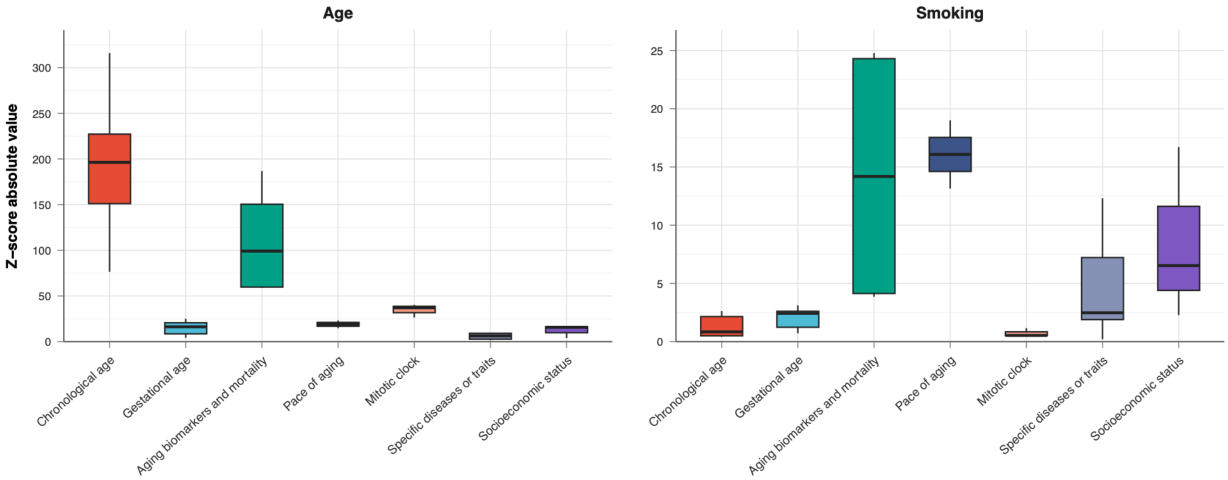


**Supplementary Figure 14. Distribution of clock-age and clock-smoking associations stratified by clock categories obtained in the TruDiagnostic sample.** Each box shows the distribution of the absolute values of association Z-scores between clock models in that category with the phenotype. Two panels present results for age and smoking, respectively. Full results are in **Supplementary Table 6**.

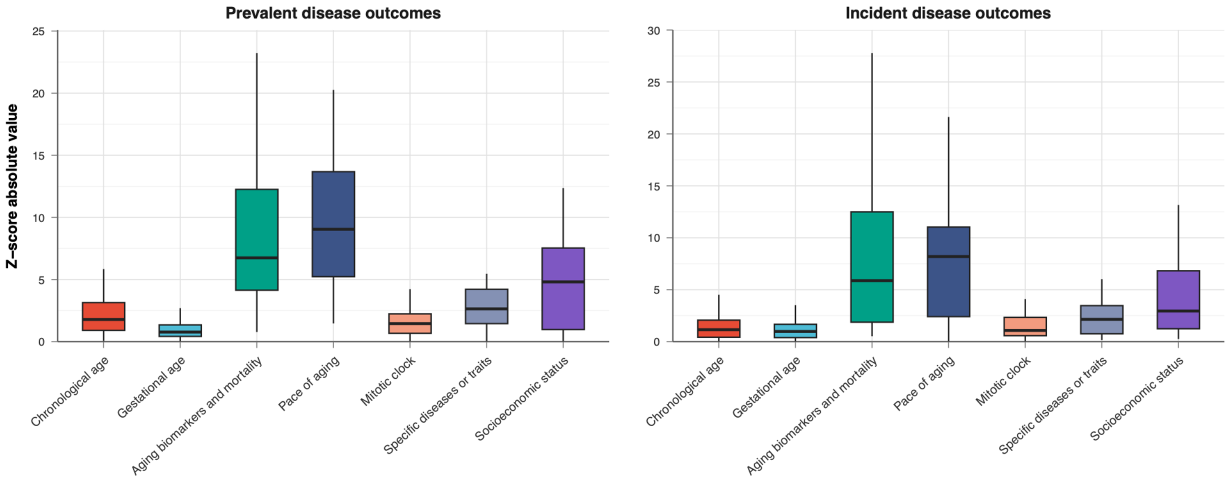


**Supplementary Figure 15. Distribution of clock-disease associations stratified by clock model categories.** Each box shows the distribution of association z-scores between clock models in that category with disease outcomes. Two panels present results for prevalent and incident disease outcomes, respectively.

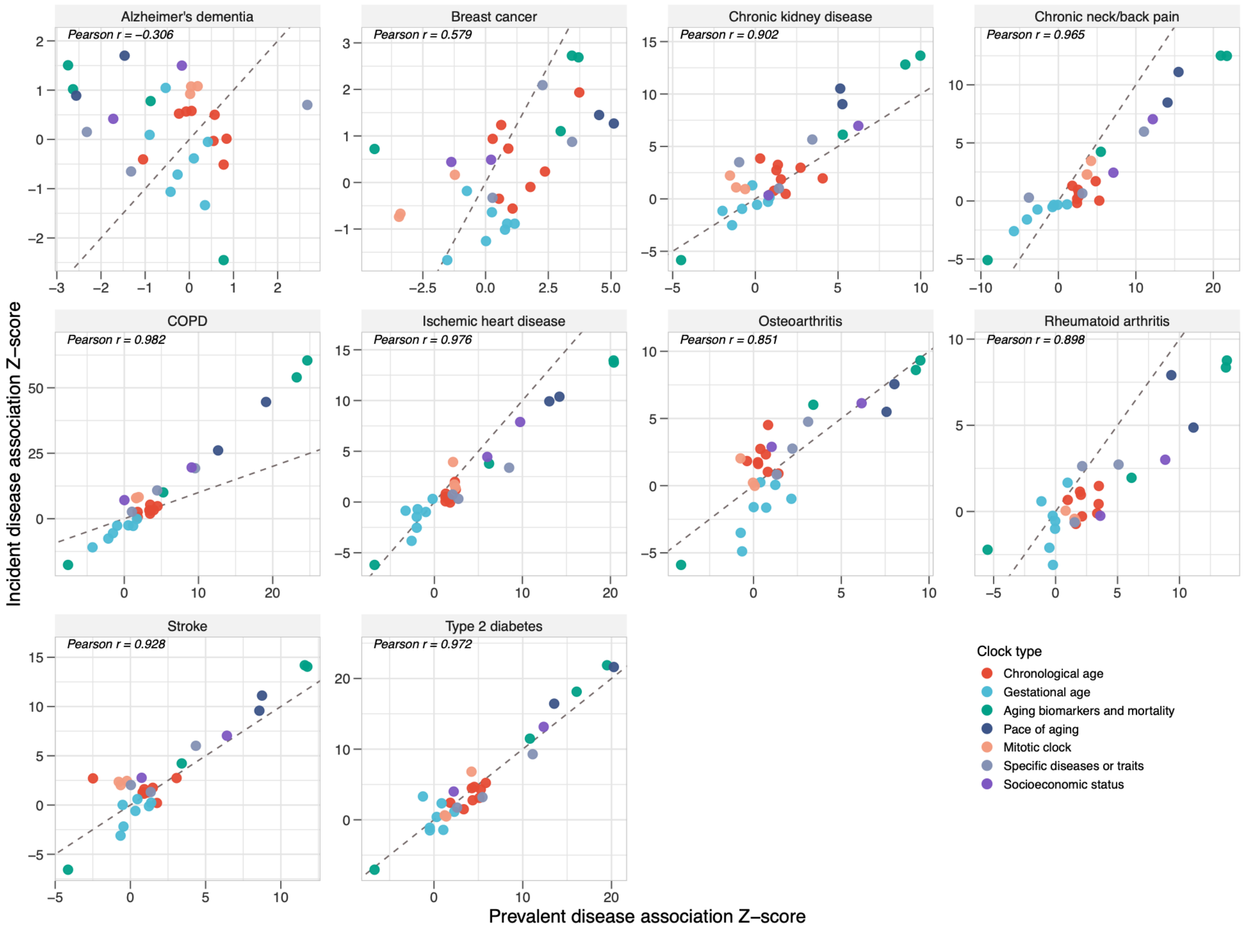


**Supplementary Figure 16. Scatter plot of clock associations with incident versus prevalent disease outcomes.** This is the same information presented in **Figure 5C** but results are stratified by disease. The dashed line in each panel is y = x.

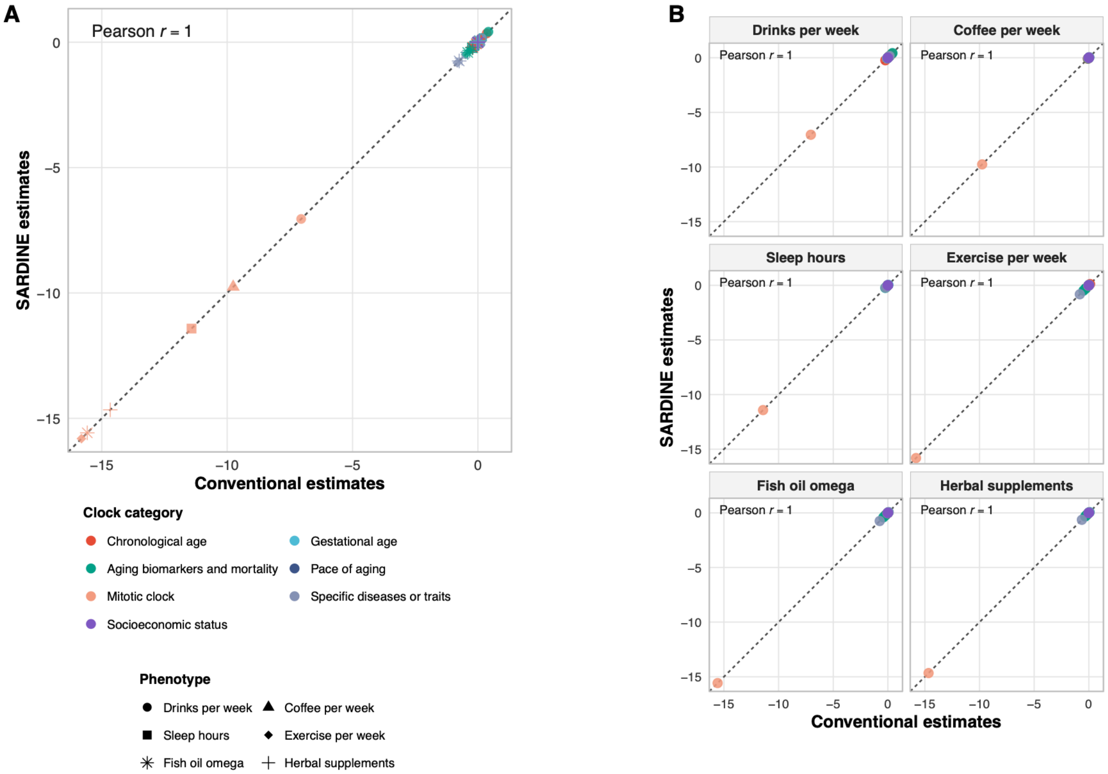


**Supplementary Figure 17. Validation of clock-phenotype associations across six TruD traits.** Panel A compares association estimates obtained using the conventional method and SARDINE, with each point representing a clock–trait pair. Panel B shows the same comparisons stratified by the six traits. Estimates from the two methods are identical, with a Pearson correlation of $r = 1$ for all traits.

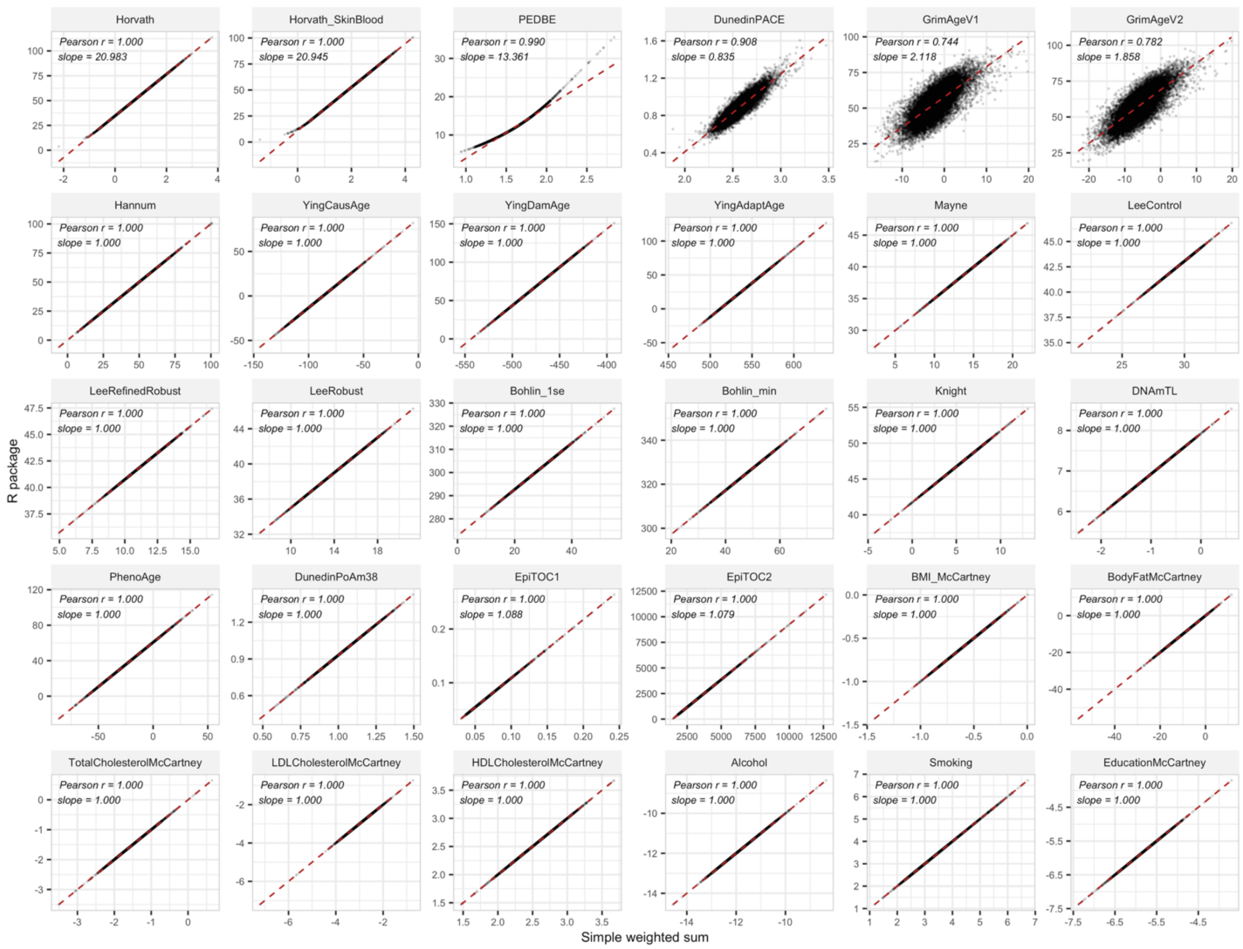


**Supplementary Figure 18. Epigenetic clock values computed as a simple CpG-weighted sum and by the meffonym R package.** The x-axis is the weighted sum of DNAm beta-values across clock CpGs without including intercept, sex, age, normalization, or transformation. The y-axis is the score returned by the "meffonym" R package. Each panel shows one clock, and each point is one TruDiagnostic sample (N = 15,677). The dashed line represents the fitted linear regression.

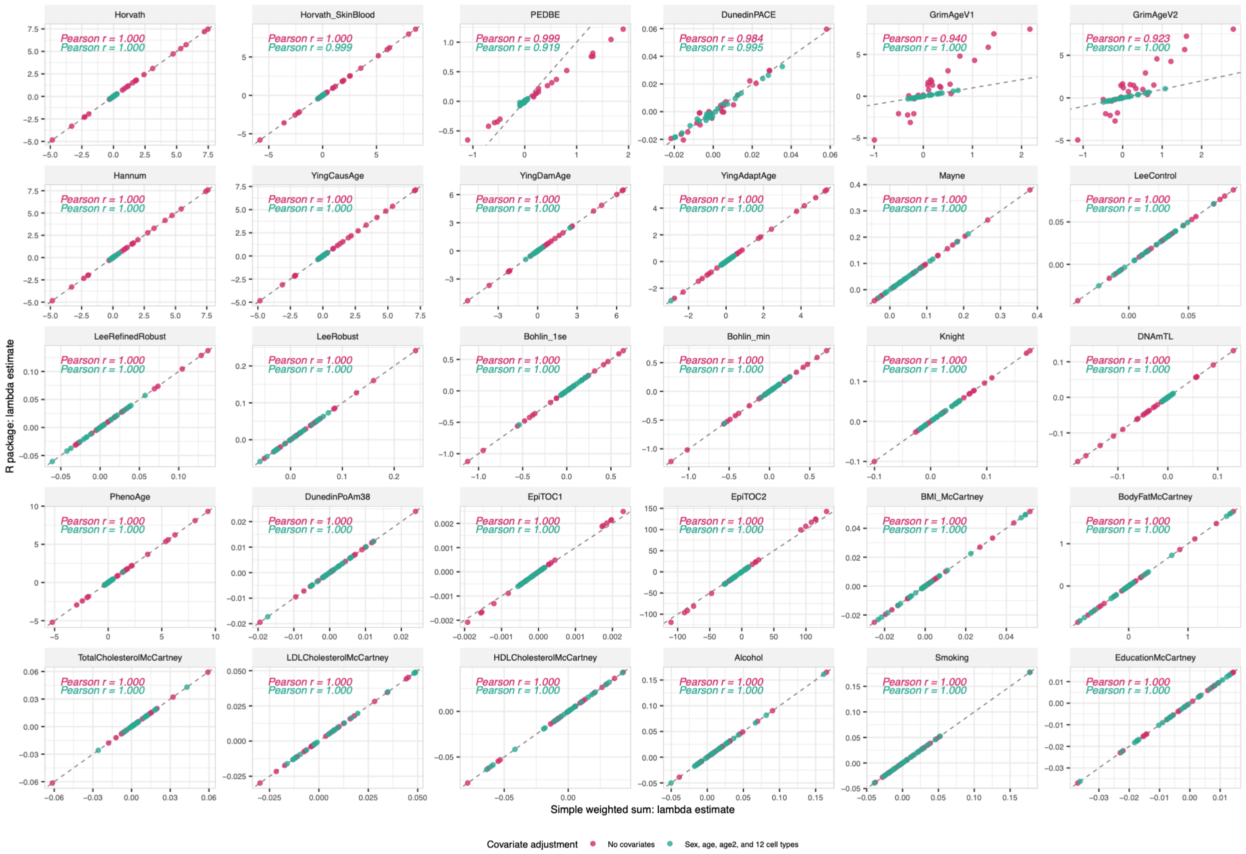


**Supplementary Figure 19. Estimates of $\lambda$ from weighted sum versus R package clock values.** Clocks were calculated either as a simple weighted sum across clock CpGs without including intercept, sex, age, normalization, or transformation; or by meffonym R package. Each point is one of 25 TruDiagnostic (N = 15,677) phenotypes under one of the two covariate sets. “Is_female” and “Age” were not included. The dashed line in each panel is y = x.

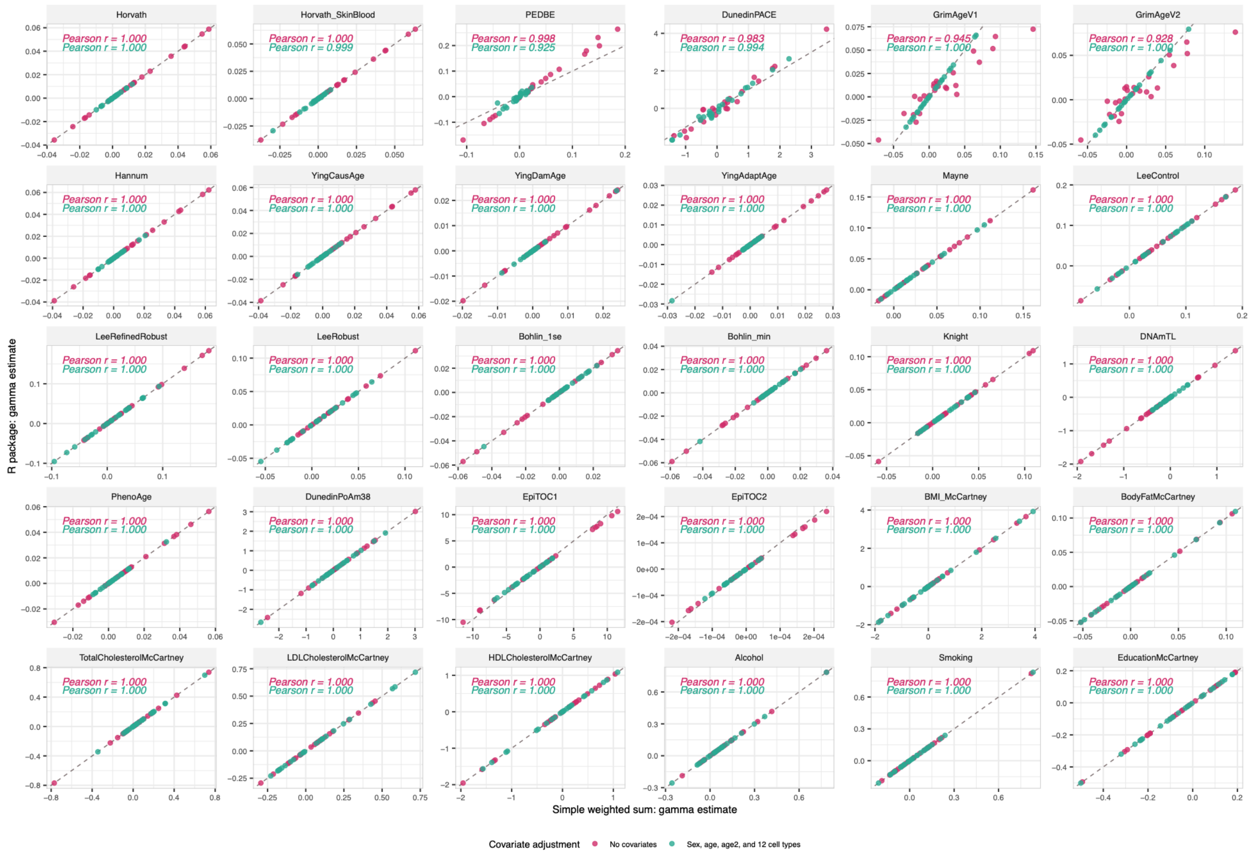


**Supplementary Figure 20. Estimates of $\gamma$ from weighted sum versus R package clock values.** Identical data as in **Supplementary Figure 19**. Binary phenotypes were analyzed by logistic regressions and the point estimates in the plots were log-odds ratios.

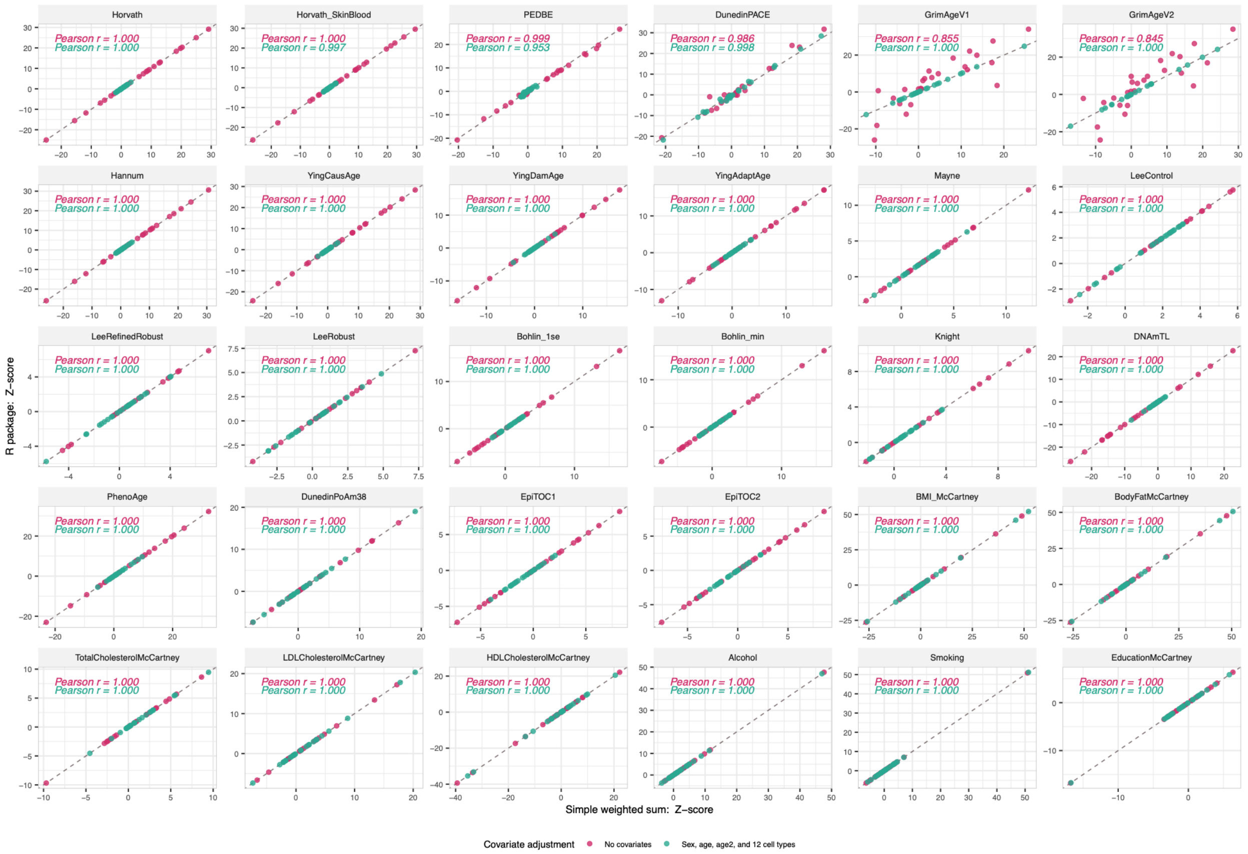


**Supplementary Figure 21. Z-scores from weighted sum versus R package clock values.** Same data as in **Supplementary Figure 19**. Z-scores are same for $\lambda$ or $\gamma$.

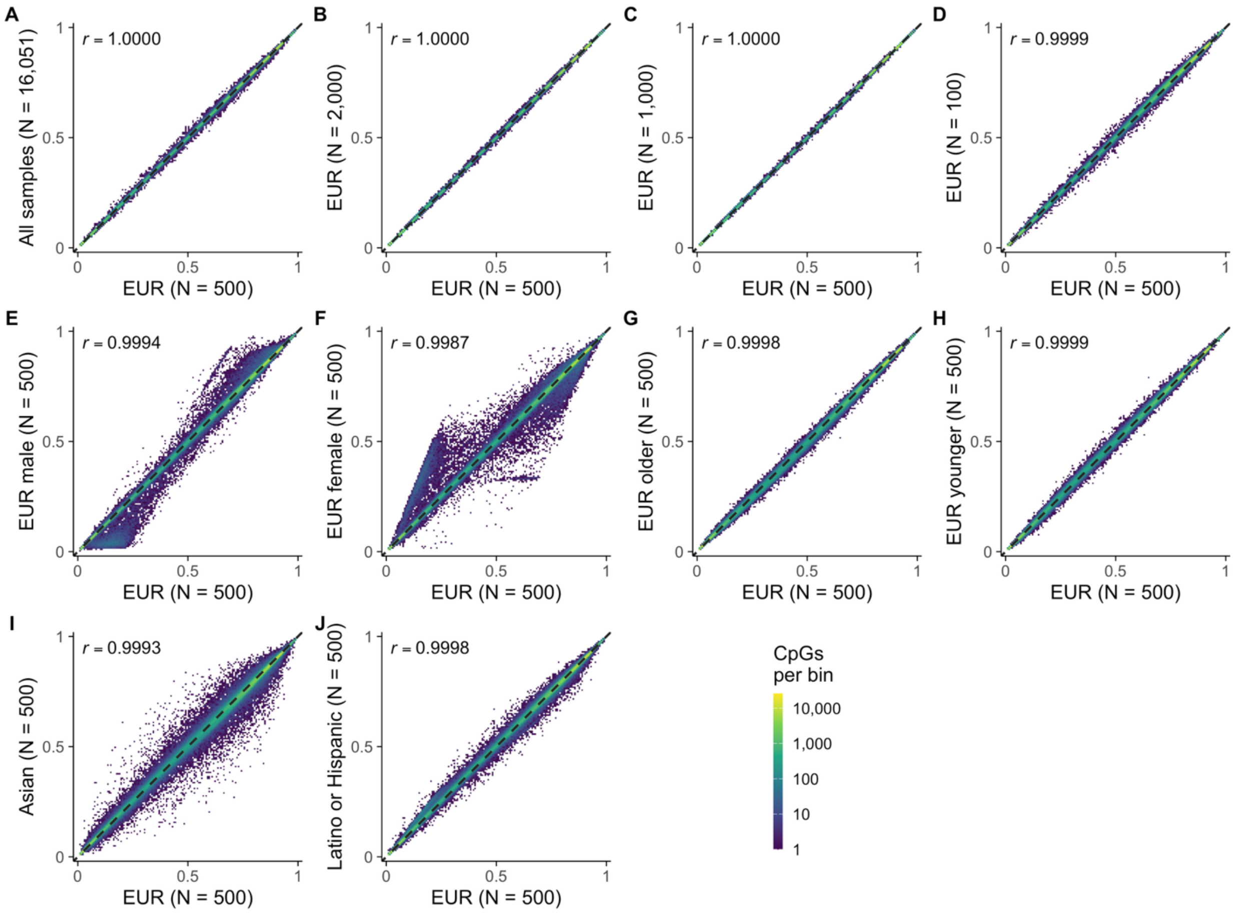


**Supplementary Figure 22. Per-CpG mean DNA methylation beta-value in each reference subset versus EUR (N = 500) subset.** Mean beta-values were computed for each CpG in each subset of the TruDiagnostic EPIC V1 data (N = 16,051). They are plotted against the means in EUR (N = 500) subset that was used for the external reference by SARDINE. A, all samples. B-D, EUR subsets with different sample sizes. E-H, EUR subsets by sex and age groups. I, J, other ancestries. Each panel is a two-dimensional histogram of 866,238 CpGs. Color shows the number of CpGs per bin on a log10 scale. The dashed line in each panel is y = x.
